# SAMP-Score: A morphology-based machine learning classification method for screening pro-senescence compounds in p16 positive cancer cells

**DOI:** 10.1101/2025.06.09.658585

**Authors:** Ryan Wallis, Bethany K. Hughes, Madeleine Moore, Emily A. O’Sullivan, Luke C. McIlvenna, Luke Gammon, Anthony Hope, Fiona Bellany, Claire Mackenzie, Charlotte Green, David Gray, Cleo L. Bishop

## Abstract

**Background:** Senescence identification is rendered challenging due to a lack of universally available biomarkers. This represents a bottleneck in efforts to develop pro-senescence therapeutics – agents designed to induce the arrest of cellular proliferation associated with a senescence response in cancer cells for therapeutic gain. This is particularly true in contexts such as basal-like breast cancer (BLBC), which often express high levels of widely reported senescence hallmarks, which has led to the designation of these subtypes as senescence marker positive (Sen-Mark+). Unfortunately, these are often cancers with the most limited treatment options, where novel pro-senescence compounds would be of potential clinical utility.

**Results:** To address these challenges, we have developed SAMP-Score, a machine learning classification tool for identifying senescence induction in Sen-Mark+ cancers. This technique builds upon our previous observation that senescent cells develop distinct senescence-associated morphological profiles (SAMPs), which can be assessed readily in traditionally challenging contexts for senescence identification, including high-throughput screens.

**Conclusions:** Through application of SAMP-Score, we have identified QM5928, a novel pro-senescence compound, that is able to induce senescence in a variety of Sen-Mark+ cancers and has potential utility as a tool molecule to explore the mechanisms and pathways through which senescence induction occurs in these cells.

## Background

Senescence is a term given to a set of related terminal cell fates; in which tumour suppressor mechanisms arrest proliferation following cellular insult to protect against malignant transformation^1^. The senescence programme is executed by cyclin-dependent kinase inhibitor (CDKi) signalling axes, most notably p53-p21 and p16, which act to prevent the phosphorylation of retinoblastoma (Rb) family members and consequently inhibit the production of E2F transcription factor targets, thus arresting cell cycle progression^2^. These tumour suppressors present a potent barrier to cancer development and, as such, are some of the most frequent mutations found in cancer cells^3^. This is reflected in the designation of senescence, and particularly its failure, as an emerging hallmark of cancer^4^. Therefore, the targeted activation of the senescence programme (so called pro-senescence) is an attractive therapeutic strategy in cancer.

Paradoxically, many cancers present with high expression of senescence markers, including the CDKi p16^5^. Importantly, these subtypes are often associated with aggressive forms of the disease leading to poor prognoses and clinical outcomes^6^. A notable example of this is basal-like breast cancer (BLBC), a subtype with significant (>90%) overlap with triple-negative histological classification^7^. These tumours stain positive for p16 so frequently, that it has been proposed as a surrogate subtype marker^8^. Crucially, this is a disease area with a profound unmet clinical need, with the triple negative status limiting the use of targeted treatment options^7^. Furthermore, modern pro-senescence therapeutic approaches, which act to mimic perturbed cell senescence pathways, including compounds such as the CDK4/6 inhibitor and p16 mimetic palbociclib, have been demonstrated to lack efficacy in cancers with high levels of p16^9^. Therefore, there is an outstanding challenge within the field to identify both pathways and compounds that can engage a senescence programme in these cancer subtypes – which we refer to as Sen-Mark+ cancers (senescence marker positive cancers)^10^.

A major obstacle to such endeavours is the reliable classification of a pro-senescence response in cancer subtypes already positive for a principal senescence marker. As standard, senescence classification requires the use of multiple complimentary hallmarks, as no single marker is universal^11^. Identifying senescence in cancer provides a particular challenge, as reliable markers such as initiation of a DNA damage response (DDR) or positive senescence-associated beta galactosidase (SA-ß-Gal) staining are often already observed in proliferating cancer cells^1,12^. Furthermore, loss of proliferation (a fundamental distinction between senescence and cancer) is insufficient alone to confidently classify a senescence phenotype, with experimental readouts such as reduced cell count attributable to other outcomes such as toxicity or slowed, rather than arrested, cell division^13,14^ . These issues are further exacerbated in contexts such as screening, which do not readily lend themselves to complex classification readouts with multiple marker stains, both practically and financially^13,15^. Therefore, there is a need for the development of scalable methods of senescence identification, particularly in Sen-Mark+ cancer subtypes^1,16^.

Previously, we utilised multiparameter assessment of high content microscopy images to demonstrate that senescent cells acquire distinctive senescence-associated morphological profiles (SAMPs)^17^. These can be visualised through simple nuclear and cell dyes (DAPI and Cell Mask) making such analysis readily applicable to high throughput contexts^18^. However, our previous use of SAMPs has focused on characterisation and subpopulation assessment of senescence models, as opposed to classification or identification *per se*. Recently, several tools have been developed, which successfully utilise machine learning (ML) algorithm assessment of cell morphologies for senescence classification^19–21^. However, it remains unclear how generalisable these methods are, for instance in contexts such as p16 positive cancer cells^22^. Furthermore, model development to date has relied on establishing ground truth through canonical senescence markers, to quantitate model performance^19,22^. Given the established limitations of senescence markers (particularly in Sen-Mark+ cancers), this represents a potential barrier to identifying novel senescence phenotypes where conventional markers prove insufficient to establish a ground truth. This underpins the discovery challenge facing the identification of pro-senescence approaches in p16 positive cancers.

Here, we build upon our previous observation that senescent cells are associated with distinct SAMPs^17^. By utilising unsupervised cluster analysis to assess the morphological profiles of a genome-wide siRNA screen in HeLa cells (p16+ cervical cancer) we developed a stacked meta-model classification tool incorporating prediction scores from multiple individual ML models which we term **SAMP-Score**. To demonstrate the potential application of SAMP-Score in p16 positive cancer therapeutic discovery, we assessed a diversity screen of 10,000 novel chemical entities in MDA-MB-468 cells (p16+ BLBC). Pro-senescence compound hits were identified through SAMP-Score classification and the effect of increasing concentration on senescence scoring was then assessed through a second dose response screen. SAMP-Score classification was then used to select a compound for validation of pro-senescence induction - QM0005928/DDD01293078 (**QM5928**), which was demonstrated to produce a senescence response in multiple Sen-Mark+ cancer lines. Collectively, SAMP-Score represents a versatile tool for therapeutic discovery and senescence classification across p16 positive cancers and has identified a promising novel compound in BLBC.

## Results

### Genome-wide siRNA screening for senescence labelling – Classical Screening Criteria

Sen-Mark+ cancer cells represent a particular challenge for developing ML classification tools for the identification of senescence. This is because the labelling of an initial training dataset generally requires a known “ground truth”, where observations are placed into the categories to allow the model to be both constructed and its effectiveness assessed ^22^. For Sen-Mark+ cancers, confidence in the labelling of any particular observation as either senescent or not is hindered by the lack of available biomarkers within the context of these cells, which, circularly, is the very reason such classification tools are needed ^10^. Furthermore, restricting the labelling of training data to be based upon a small set of hallmarks with known limitations also has the potential to miss uncommon, idiosyncratic or novel phenotypes, potentially limiting the application of such tools, particularly for identifying new senescence contexts.

Therefore, where traditional ML model development would first identify conditions of senescence, before measuring their morphologies, for our approach we worked in reverse. First, we generated and assessed as diverse a range of potential cell morphologies as possible before exploring different approaches to labelling those which resembled our previously observed senescence-associated morphological profiles (SAMPs) as senescent. This allowed us to capture the breadth of potential senescent cell morphologies, with the ultimate goal of serving as a training dataset for development of a flexible tool to identify senescence in the context of Sen-Mark+ cancers.

We performed a genome-wide siRNA screen in p16+ cervical cancer cells (HeLa). This screen consisted of 63 384-well plates, with a pool of 3 siRNA per well per target, giving a total of 21,658 treatments (excluding controls), representing a comprehensive perturbation of cellular pathways. To assess the morphology of these cells, we refined our previous SAMP methodology to limit the influence of co-linearity, aiding computing time and interpretability (Supplemental Figure 1). This was achieved by taking the morphology profiles from all individual siGLO control cells (∼1.16M cells) and removing redundant features, as identified by Pearson correlation assessment. This resulted in a parsimonious set of 36 features, which were then assessed for all siRNAs and control conditions (Figure 1A). As expected, a broad range of profiles were generated, with most siRNAs producing little to no change compared to the siGLO control profile (which would appear as an entirely black heatmap). However, a heterogenous set of altered morphologies was also produced by many treatments, which appeared similar to the SAMPs we have previously observed in senescence. We then explored several options for establishing senescence “hits”. First, we assessed the magnitude to which treatments produced a combined reduction in cell count and increase in cell area, which have previously been employed as readouts in senescence screens^25^. We filtered the profiles according to several thresholds in these two features, which served as high (≥3 Z-score change), medium (≥1.92 Z-score change) and low (≥1 Z-score change) stringency criteria (Supplemental Figure 2A-C). We observed that the potency of morphology changes followed the stringency of the senescence thresholding, suggesting that the altered profiles were associated with conditions that produce a reduction in cell count and general increase in cell area. Importantly, we also demonstrated no correlation between the overall cell number and the cell area, suggesting that morphological changes are not tied to a simple confounding influence such as confluency, and emphasising the limitation of relying on cell counts as a senescence readout alone (Supplemental Figure 2D-E). These, canonical screening thresholds were a useful starting point for suggesting altered morphology profiles are associated with conditions that would traditionally have considered senescent and corroborate our previous reports^25^. However, we previously observed significant heterogeneity in the composition of SAMP profiles between different senescence models. Given this, we sought to explore other methods for classifying profiles as senescent, in order to broaden the range of profiles within this class and potentially capture less conventional phenotypes.

**Figure 1:**
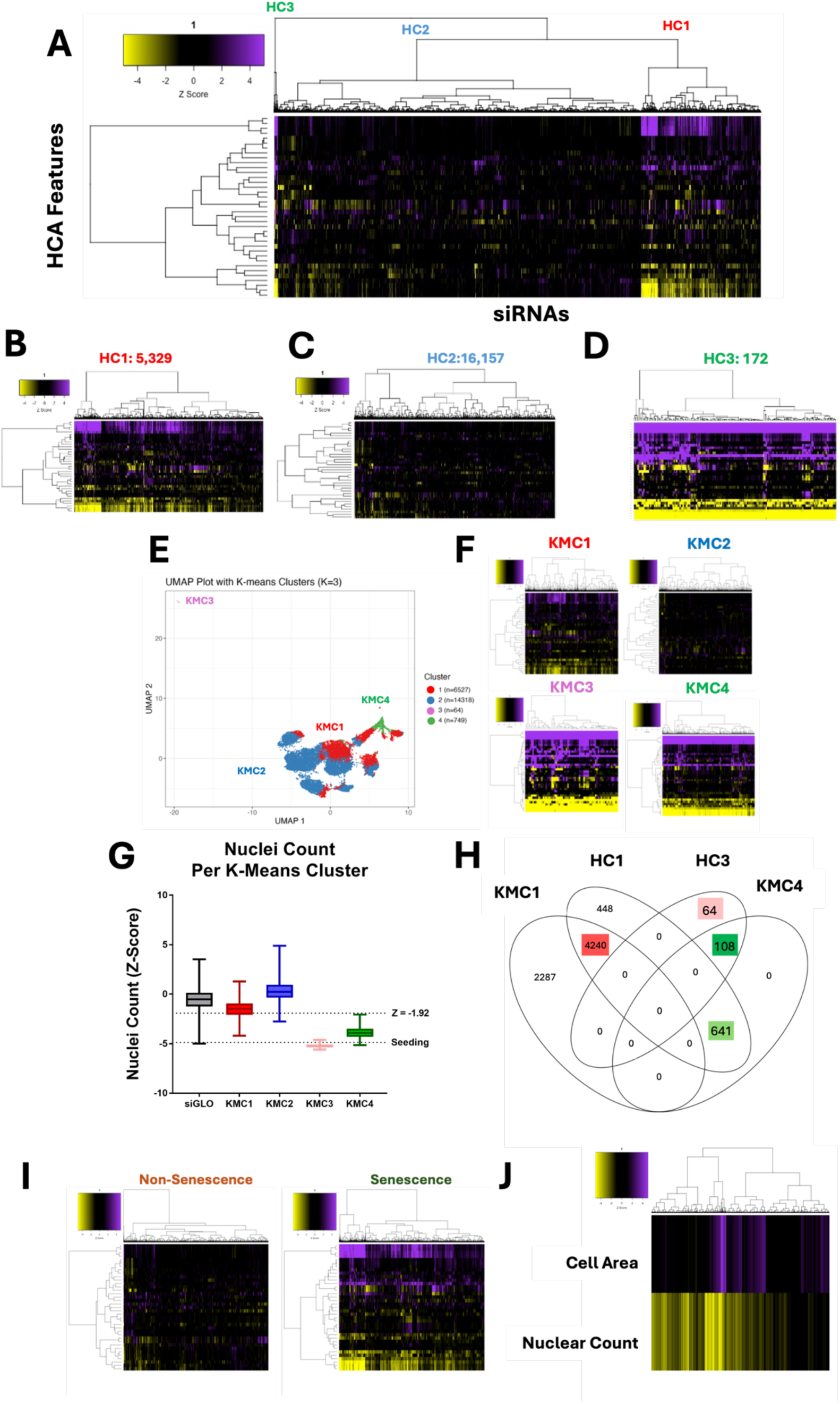
Genome-wide siRNA HeLa Screen and Cluster Based Senescence Labelling. A: Heatmap representing high content analysis feature (HCA; y-axis) profiles of a genome-wide siRNA screen. Treatments (siRNAs; x-axis) were grouped into three through hierarchical clustering. B-D: Heatmap of individual hierarchical clusters. E: UMAP plot showing 36 feature profiles of every treatment within the screen, labelled with K-means clustering groups. F: Heatmap profiles of K-means clusters. G: Nuclei counts for treatments in each K-means cluster H: Venn diagram showing overlap between hierarchical and k-means clustering. I: Heatmap profiles of treatments labelled as either Non-Senescence or Senescence. J: Heatmap profile showing cell area and nuclear count Z-Scores of all senescence conditions. In all heatmaps, purple indicates positive modulation and yellow negative modulation of greater than 1.96 Z-scores from siGlo control. Black indicates a Z-score between –1.96 and 1.96.

The advent of large-scale profiling techniques, such as RNA sequencing, has provided greater depth into our understanding of the pathways that are altered in senescence. Of pertinence to this work, the SENCAN classifier and SENESCopedia database, provide an insight into directionality of target expression in senescent cancer cells^2^ ^6^. We explored whether siRNAs against targets which are downregulated in SENCAN/SENESCopedia would lead to the generation of distinctive senescence phenotypes (Supplemental Figure 3A-B). We also performed this process for targets which fall within the KEGG gene ontology pathways “Cell Cycle” and “Senescence” (Supplemental Figure 3C-D). In general, we found this produced a far more mixed set of profiles than those we had generated with our previous thresholding strategy, with most profiles appearing unchanged compared to the control, suggesting they were not SAMPs. This may be due to factors such as variability between cancer types (which was strongly observed by the SENCAN authors) or that the downregulation of many of the targets may be a consequence but not a driver of senescence induction. Therefore, we concluded that a biased, pathway driven approach was inadequate for identifying profiles that could be labelled as senescent.

### Genome-wide siRNA screening for senescence labelling – Unsupervised ML Criteria

Next, we utilised several methods of unsupervised ML, in order to group profiles based on similarity, rather than through rigid screening thresholds. We then inspected these algorithm generated groups, to determine whether any resembled SAMP profiles. The first method used was unsupervised hierarchical clustering (HC). This algorithm groups together profiles based on pairwise distances and is represented by the dendrogram on the x-axis of the heatmap (Figure 1A). Whilst the clustering itself is unsupervised, the decision of where to cut the tree to determine the size of groupings is provided by the user. This could range from 1 cluster containing all siRNAs to 21658 clusters of 1 siRNA per group. We chose to break the screen into 3 clusters, which were the clearest from the dendrogram and which also appeared to have the longest branches, signifying the greatest difference between groups. When these clusters were visualised separately, their distinctions became obvious. HC1 (Figure 1B) and HC3 (Figure 1D) contained 5,329 and 172 siRNA respectively, and represented clear morphological changes from the siGLO control, reminiscent of SAMP profiles, with HC3 appearing to contain more potent profiles than those in HC1. HC2 (Figure 1C) in contrast, contained 16,157 siRNA profiles, which demonstrate comparatively little change from the siGLO control. Together, this suggested that the unsupervised hierarchical clustering had successfully split the siRNAs into those that change the morphology of the HeLa cells, and those that do not.

Next, to further explore the range of potential morphologies as well as additional subgroupings, we then plotted the profiles via UMAP dimensionality reduction, where each point on the chart represents the 36-feature profile from a single treatment (Figure 1E). We also employed an alternative K-means clustering (KMC) algorithm, to explore the effect of using different clustering methods on profile groupings. The UMAP allowed us to identify a small group of siRNAs that produced a phenotype which was distinct from the other profiles. However, isolating this group within its own cluster required increasing the total number of groups to four (K=4). The heatmap profiles (Figure 1F) shows the 64 siRNAs in this group, KMC3, generated a potent phenotype. The large reductions in cell count for treatments in this cluster, generally below the level of seeding, made it evident that these treatments had induced toxicity in the cells as opposed to senescence (Figure 1G). By contrast, KMC4 contained 749 profiles that had clear morphological changes reminiscent of the SAMPs we have described previously, whilst KMC1 contained a much larger range of 6,527 profiles that were variable in potency. The intensity of profile changes observed between these clusters also mirrored the relative reduction in their respective cell counts. As before, a large cluster, KMC2, contained 14,318 profiles that differed little from the control, indicating most treatments do not alter morphology. Interestingly when looking at a Venn diagram of how the two clustering algorithms overlap, we see that through KMC we have been able to split the 172 potent profiles from HC3 into 64 toxic profiles (pink box KMC3 - Figure 1H) and 108 strong senescent (dark green box KMC4 - Figure 1H). We can also see a large overlap between the HC1 and KM1 (red box-Figure 1H) which represent the profiles with more a modest change, but also with KM4 (light green box - Figure 1H) accounting for the more potent profiles observed in HC1. For the purposes of labelling our cluster-based training data, we took the entirety of KMC4, as well as those treatments that overlapped between HC1 and KMC1 (light green, dark green and red boxes - 4989) and labelled these as senescent. This strategy was deliberately designed to be inclusive in order to capture as many potential phenotypes as possible (Figure 1I). The trade-off with this approach was that any model developed based on this data may have been more prone to generating false positives, which was later accounted for and minimised through application of a stringent decision boundary (see below).

Next, we sought to understand how the clustering-based strategy for senescence labelling would compare the “traditional” approach based on rigid thresholds for nuclei count and cell area. 88.5% of those siRNAs which would have been considered hits according to the classic criteria of reduced count and increased area (using the medium stringency threshold of 1.92 Z-Score change) were contained within the senescence grouping. This suggests that the unsupervised methodology does not miss many hits that would previously have been identified. Of the 11.5% of siRNAs not included, the vast majority (53/60) were confined to KMC3, representing toxic conditions and supporting the concept that a clustering approach to data labelling has allowed these to be separated from senescence. Interestingly for our goal of identifying novel senescence morphologies, only 9.3% of the clustering approach labelled senescent conditions satisfy the Z-Score criteria for change in **both** reduced cell count and increased cell area. This means that as a screening criterion, the unsupervised clustering has expanded the list of potential senescence hits and thus broadened the range of phenotypes we are considering beyond those that are simply “large”. This is clear when the cell counts and areas of the senescence treatments are visualised (Figure 1J), with most satisfying the reduction in count fundamental to senescence but being lost as hits due to a failure to achieve the increased area threshold. Therefore, applying the additional sophistication of unsupervised morphological assessment to the hit detection process captures a greater number of potential senescence phenotypes, and allows distinction from both toxicity and proliferation. Validation of this is made difficult, by the nature of the problem we are trying to solve, the Sen-Mark+ status of the cells, which limits the utility of senescence markers. Given that our goal was to identify novel phenotypes that may not satisfy conventional markers anyway, we instead decided to explore the practical implications of our expanded definition and moved to develop classification tools based on this approach to identifying senescence.

### Developing an ML method for detecting senescence in Sen-Mark+ cancer cells

By establishing thresholds within the HeLa screen for what we consider senescence (SEN) or non-senescence (NonSen), we can utilise the data as a training set for developing ML classification tools. These have previously been developed through labelling according to canonical senescence markers in a range of contexts successfully ^19–21^. Here, we did so using the unsupervised clustering approach to data labelling. The details of this are outlined in (Figure 2) and methods above, but briefly: The labelled HeLa screening data was partitioned into 80% training and 20% testing. Due to heavy class imbalance in favour of the NonSen condition, this data was then randomly undersampled to achieve classes of a similar size. The excluded training data (which comprised only NonSen siRNAs) was then recycled into the testing data set. The undersampled training data was then split in half, with 50% dedicated to training individual ML models and 50% for testing. The prediction coefficients from the latter were then used to train an ensemble meta model. The individual models which comprised this covered a range of ML types including Logistic regression, Lasso regularisation, Elastic Nets, Support Vector Machines (SVM), Random Forrest (RF), Multiple Discriminant Analysis (MDA) and Neural Networks (NN). The meta model takes the predictions from each of these models and forms a consensus prediction (model stacking). Each model is individually used to assess the initially removed (20% + recycled) testing data and its performance assessed. The combined predictions are then evaluated with the meta model to determine a final prediction, the accuracy of which can be compared to each model individually. This approach was selected to be as comprehensive as possible, when assessing the applicability of different ML approaches to identifying senescent cells.

**Figure 2:**
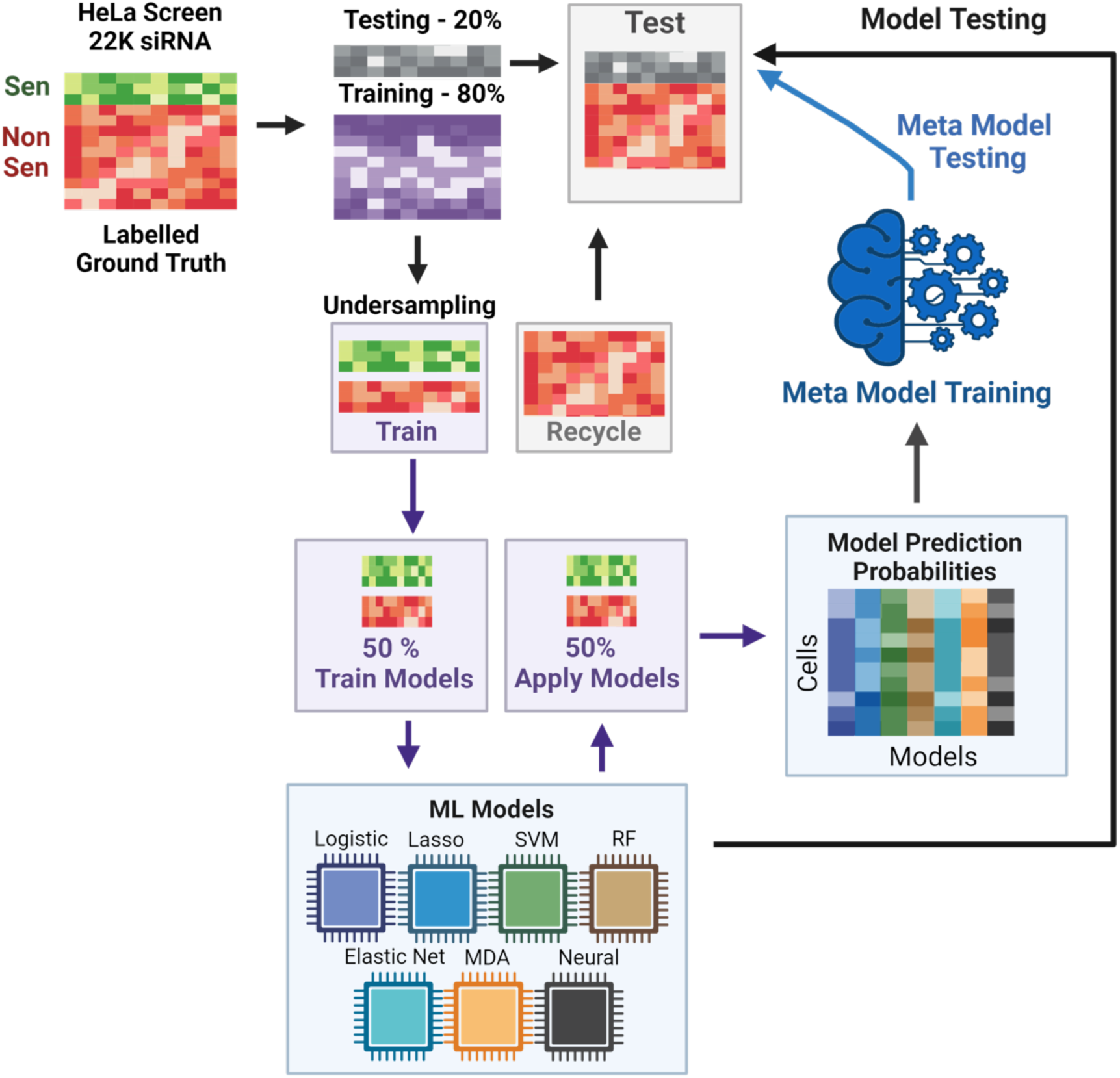
Overview of SAMP-Score Model Development. Each model was assessed according to a range of criteria, which are visualised in the model metrics (Figure 3A) and confusion matrix heatmaps (Figure 3B), as well as neural network map (Figure 3D) and ROC curves (Figure 3E; Supplemental Figure 4). The model metrics are nuanced and can be misleading when viewed in isolation. For instance, accuracy is a measure of correct predictions and is often relied upon as a single readout of model performance. But in a hypothetical example where there are 99 majority cases (e.g. NonSen) and 1 minority case (e.g. Sen) then a model may be 99% accurate by simply always predicting NonSen; but this would not be a useful tool. Therefore, particularly in senescence research where instances are likely to be imbalanced, particular care in assessing model performance must be taken.

Interestingly, we did observe high model accuracy in most cases (>90% accuracy in all individual models), which aligns with similarly high model performances previously reported^21^. Of the individual models, the SVM performed best, but by stacking the predictions from all models we were able to increase performance as seen in the meta model metrics (Figure 3A-B). However, each of the models is hindered by a high ratio of false positive (FP) to true positive (TP). This is recorded in the positive prediction value (PPV), which is the proportion of positive classifications that are TP – i.e. how good are the hits? These metrics suggest the models are somewhat “reckless” as whilst in general they will not miss many TPs, they do seem to “over predict” and pick up a high number of FPs. In a screening context this would equate to a high rate of false hits. It might be that in the context of senescence screening (where many more cases of NonSen are predicted) we can accept a lower PPV, because the class imbalance means that the ratio of TP vs FP is likely to always be low, given that there are many more negative cases to potentially misclassify than positive ones to classify. Importantly, the low rate of false negatives (FN), means that the models are generally not missing hits, and the very low ratio between TN and FP means that the FP rate is in fact very low, which contributes to the high accuracies observed. However, achieving as low a ratio of FP to TP as possible (high PPV) would limit the identification of false hits, and the model stacking approach appears to improve this slightly from 56% in the best performing individual model (SVM) to 58.7% in the meta model. This can be further improved by adjusting the decision boundary, a stringency measure that determines how confident the model must be to classify something as senescent. For the individual models this was set at the oft-used value of 0.5 but we see that increasing this value to a maximum of 0.95 improves both the accuracy (94.8 % to 97.4%) and PPV (58.7% to 81.5%). This comes from fewer FP classifications but at the expense of losing some TPs. To add further nuance, it is important to consider the process through which initial class labelling was performed. In our unbiased set up, we were *inclusive* with respect to senescence classification and thus consider it is more likely that there were instances of mislabelling siRNAs as senescent than missing those that should have been. This might mean the model has been misled at the training stage to label weak profiles as senescent, which would be anticipated to give rise to a greater raw number of FPs. Therefore, we think it is reasonable to accept a measure of TP loss in order to limit the number of FPs, thus leaving us with only the most confident predictions and avoiding low quality hits in any screening applications. As such we took a stringent decision boundary threshold of 0.9 when setting the final model.

**Figure 3:**
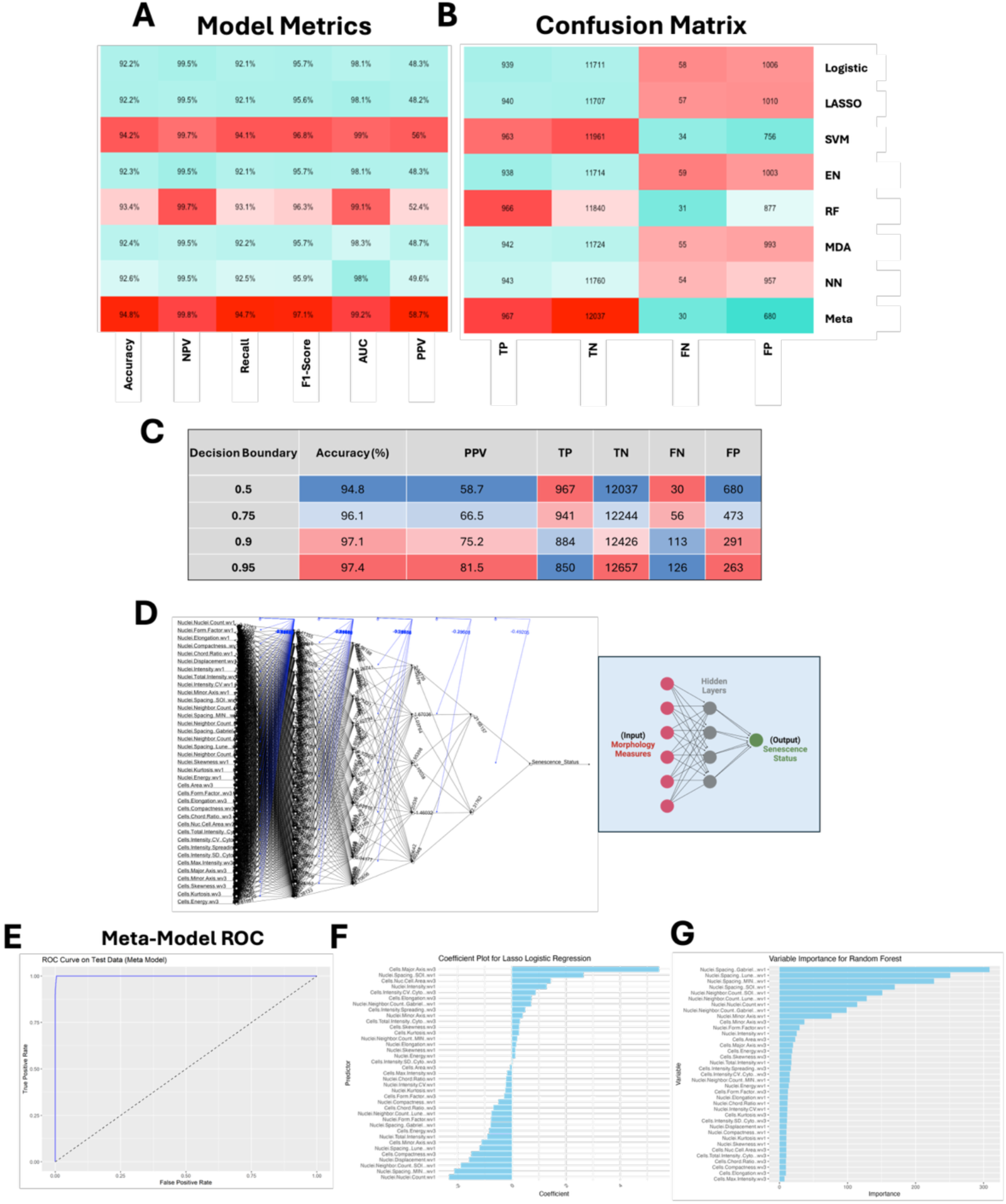
SAMP-Score Model Development and Metric Assessment. A-B: Machine learning (ML) model assessment metrics and confusion matrix for all individual ML models and stacked meta-model (SAMP-Score). NPV: Negative Prediction Value, AUC: Area under curve, PPV: Positive Prediction Value, TP: True Positive, TN: True Negative, FN: False Negative, FP: False Positive. C: Effect of altering decision boundary position on model metrics and confusion matrix. D: Neural network model. Features are input nodes and lead to a prediction of Senescence or Non-Senescence. E: Receiver operating characteristic (ROC) curve for stacked meta model. F-G: Model feature contributions to Lasso and Random Forest models.

The construction of the meta model relies on each of the composite models. These have a literally infinite number of hyperparameter combinations that could be changed, tweaked or optimised in each case (including the decision boundary), and the question of when to stop refining the models is one that is often debated in ML. Here, we have taken a heuristic approach, which is defined by a model only needing to be good enough to perform its intended task adequately, rather than the perfect version of what it could be. We have also included some insight into the decision-making process that were performed by a few of the models. The Lasso regularisation coefficients (Figure 3F) demonstrate the magnitude and directionality of individual features to the model performance, whilst the Random Forrest variable importance (Figure 3G) provides a measure of the contribution of each feature within the decision tree. This is an advantage of feature-based analysis over black-box approaches such as CNNs, aiding interpretability and leaving the potential to further refine model complexity^22^. Ultimately, we have developed a model that appears to perform well at its intended task of identifying senescent cells (as labelled by the unbiased approach). Individual model performance varies but is improved by stacking into an ensemble meta model. Whether the marginal gains in performance justify the significant additional complexity in construction will depend on the specific task to which the model is applied, but we chose to continue with the most sophisticated version of our ML classifiers (the meta model), which from here on is referred to as SAMP-Score – given that this produces a prediction value of senescence for any given treatment.

### Using SAMP Score to Identify Novel Pro-Senescence Compounds

The ultimate litmus test of any ML model is not whether it generates high performance metrics when applied to a testing dataset, but rather whether it is generalisable enough to serve the intended real-world function on unseen data sources. We set out to develop a model which would have utility as a screening tool to aid the identification of compounds that induce a senescence response in Sen-Mark+ cancer cells. To assess the utility of SAMP-Score towards this goal, and to assess its generalisability across different p16 positive cancer lines, we made use of screening data that had previously been generated in a MDA-MB-468 BLBC cell line, emphasising another advantage of SAMP-Score – the ability to re-mine and categorise archival image stacks.

The first of these screens was a diversity library of 10,000 novel chemical entities, which were screened at two doses (10µM and 50µM). As before, 36 feature HCA morphology profiles were generated for every compound and dose (Figure 4A). These profiles were then assessed using the SAMP-Score algorithm to classify the senescence state of each condition. In order to compare the SAMP-Score classifications to those that would have been generated through classical screening thresholds, we compared nuclear count and cell area for all conditions, whilst overlaying the SAMP-Score class predictions. It became clear that SAMP-Score avoids detecting proliferating conditions that would not have met a classic threshold for reduced nuclear count or increased cell area (Figure 4B). However, SAMP-Score is also able to make more nuanced classifications and appears to detect conditions that reduce nuclear count and increase cell area, but in the latter case not by the magnitude required by traditional screening cutoffs. Furthermore, SAMP-Score also avoids classifying conditions with extreme reductions in cell count and increases in cell area respectively, which likely represent conditions producing cytotoxicity rather than pro-senescence. To emphasise this further, we reproduced the SAMP-Score model as before but using classical screening thresholds to label our ground truth, rather than the unbiased clustering method (Supplemental Figure 5). When this model was applied to the compound screen, we see a very clear cut off in phenotype that aligns with the nuclear count and cell area screening thresholds used in the training data (Figure 4C). It is important to emphasise that we have not applied these thresholds here, but the model has learned to essentially replicate them. Additionally, we are able to see the particularly strong influence of cell area in the Lasso and Random Forrest feature assessments (Supplemental Figure 5C-D), highlighting the advantage of feature-based analysis for the purposes of interpretability over so called black box techniques such as CNNs^15,19,22^. Most crucially, we see that even if traditional screening thresholds are used to train a ML model rather than being rigidly applied, we do not see the exclusion of conditions that produce extreme phenotypes from being placed in the senescence class. This emphasises the value of our unbiased cluster-based labelling of senescence ground truth during model development, as it allows for far more nuanced classifications, distinguishing both toxicity and proliferation from senescence. This can be appreciated further if the morphology profiles are assessed according to senescence condition and dose. At the low dose (10µM), the Non-Sen condition comprises mostly very weak profiles comparable to the vehicle control (Figure 4D). However, at the high dose (50µM) the Non-Sen condition now also contains a large number of very potent profiles, reminiscent of the HeLa clustering in KM3 (Figure 4F). These are toxic conditions. Furthermore, we see the potency of profile in the senescence classifications increase with dose (Figure 4E and G), aligning with the principle that the cellular perturbations elicited by the compounds are enhanced with increased concentration.

**Figure 4:**
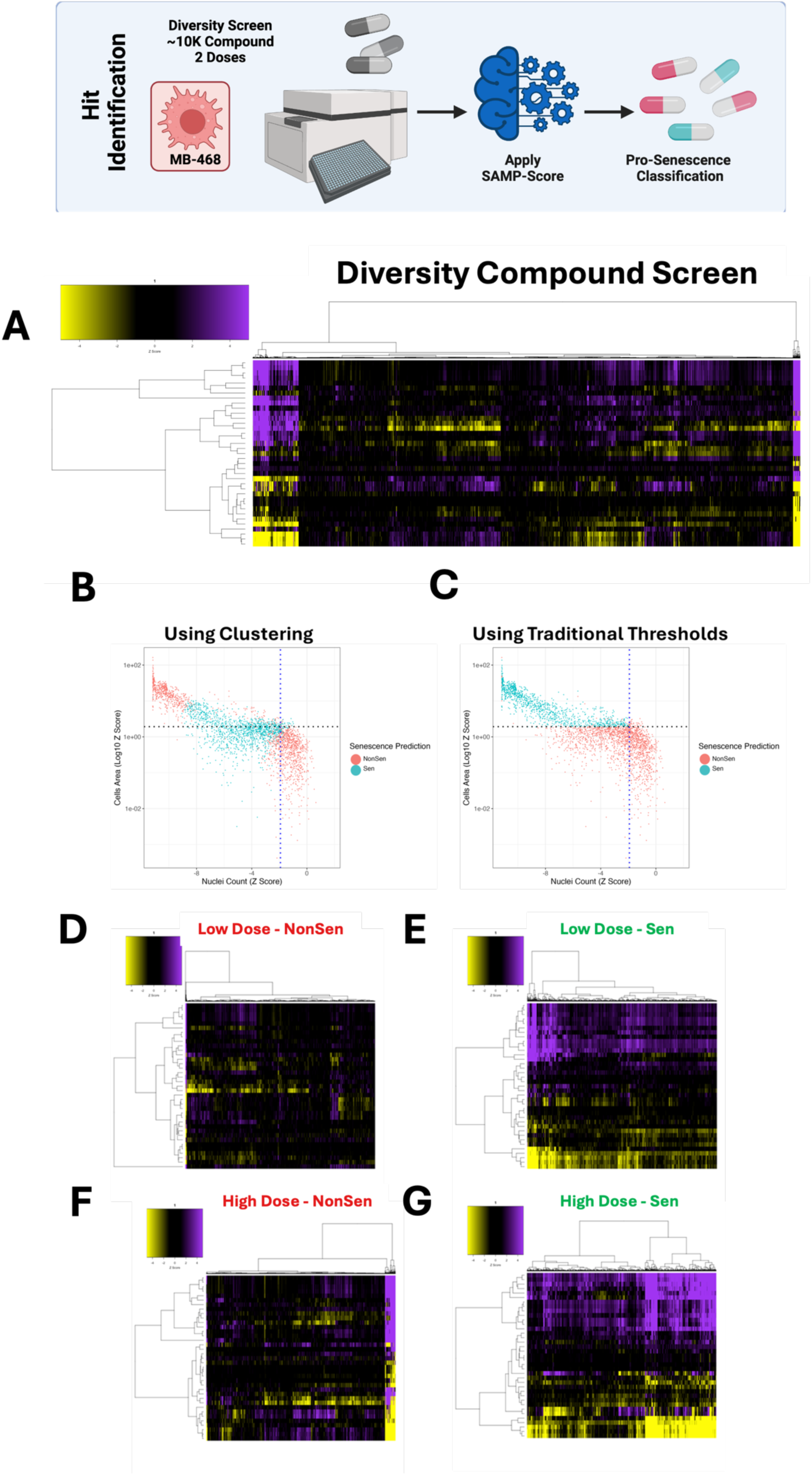
Diversity Library Compound Screen. A: Heatmap representing high content analysis feature (HCA; y-axis) profiles of a compound diversity library screen (Compounds; x-axis) performed in MB-468 cells. B-C: Scatter plots showing Z-scores of nuclear count and cell area (log10) for all compound treatments. Points are coloured Non-Senescent (red) or Senescent (blue) according to SAMP-Score classification based on models constructed with either cluster based or traditional threshold-based data labelling. D-G: Heatmap profiles for treatments classified as either Non-Senescent (NonSen) or Senescent (Sen) by SAMP-Score at both low (10 µ M) and high screening doses (50µM). In all heatmaps, purple indicates positive modulation and yellow negative modulation of greater than 1.96 Z-scores from DMSO vehicle control. Black indicates no change.

To further explore the influence of compound dose on SAMP-Score classification we next assessed a second compound screen comprising 10-point dose responses (DR) from 447 compounds that comprised a subset of the diversity screen. Whilst the compounds comprising this screen were selected before the establishment of the SAMP-Score methodology, it nevertheless represents a useful dataset, with 50% of compounds in the DR screen having been classified as senescent at either of the doses in the diversity screen according to SAMP-Score. Assessing morphology profiles by dose, we see a very clear increase in profile potency as dose increases (Figure 5A). As before, the association between nuclear count, cell area and SAMP-Score prediction demonstrates that only profiles that represent moderate change from the vehicle control are classed as senescent, with SAMP-Score once again avoiding detection of both proliferating and toxic doses (Figure 5B). This can be seen more clearly when SAMP-Score is visualised by dose, with a clear pattern for most compounds of moving from Non-Sen classification at low doses, through increased likelihood of being labelled as senescent and then back to Non-Sen as the concentration reaches toxicity (Figure 5C). This clearly supports our earlier observation, that SAMP-Score provides a means of separating senescence from both proliferation and toxicity, something not achievable with the application of standard screening thresholds. Therefore, we concluded that SAMP-Score was a tool readily applicable for pro-senescence screening and is able to identify compounds that elicit a phenotype comparable to that which we labelled as senescent ground truth in the HeLa cells.

**Figure 5:**
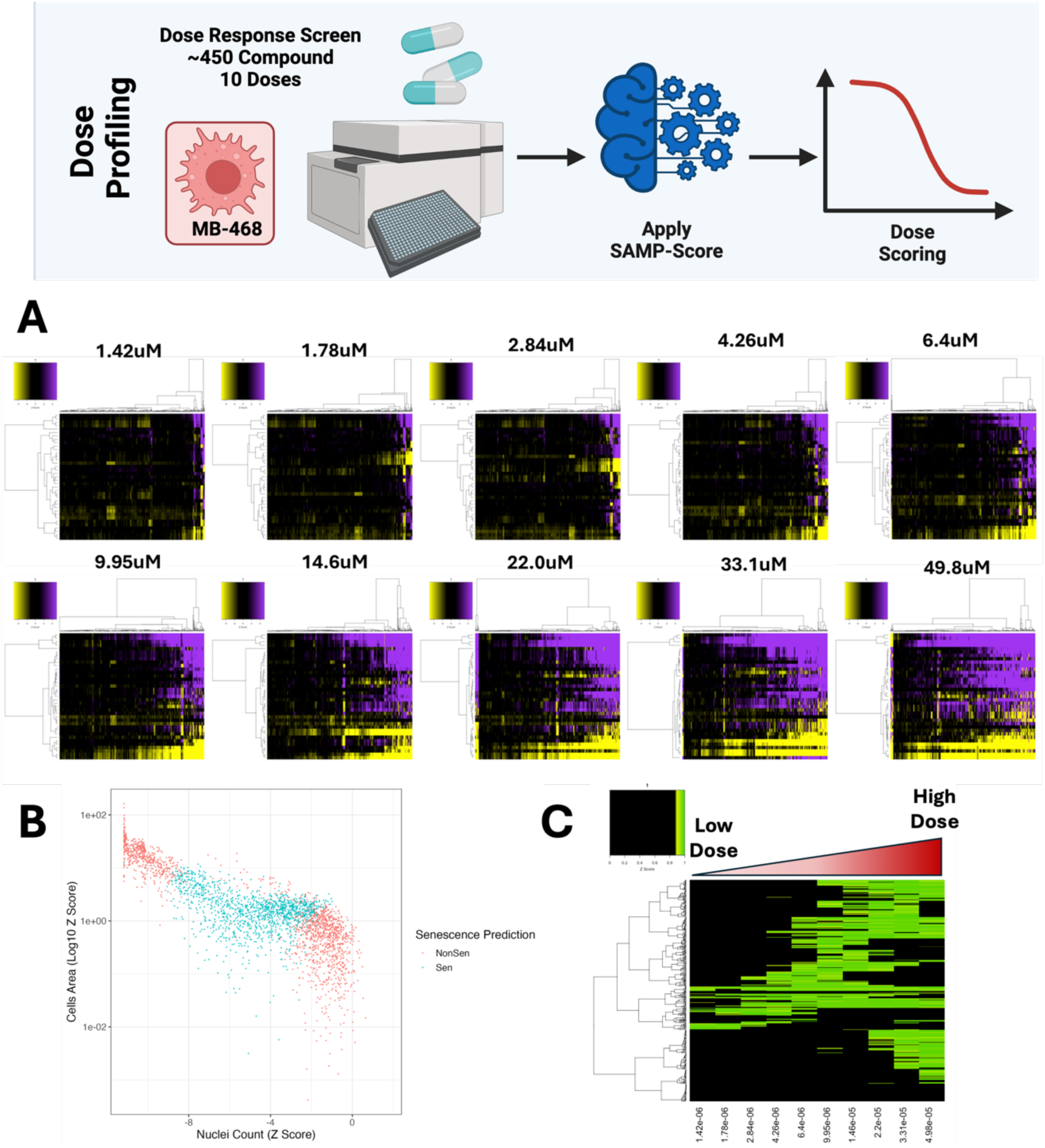
Dose Response Compound Screen. A: Heatmap representing high content analysis feature (HCA; y-axis) profiles of a dose response compound screen (Compounds; x-axis) performed in MB-468 cells. In all heatmaps, purple indicates positive modulation and yellow negative modulation of greater than 1.96 Z-scores from DMSO vehicle control. Black indicates no change. B: Scatter plot showing Z-scores of nuclear count and cell area (log10) for all compound treatments. Points are coloured Non-Senescent (red) or Senescent (blue) according to SAMP-Score classification. C: Heatmap showing SAMP-Score prediction co-efficient for all compounds (y-axis) and doses (x-axis). Black indicates score of <0.9 which is scored as non-senescence.

### SAMP-Score as a tool for pro-senescence therapeutic discovery

Through the application of SAMP-Score to the diversity screen, we were able to generate a list of compounds that produced a senescence response at one or both doses screened. In the DR screen we were also able to see that typically compounds which induce senescence do so in a dose dependent manner, before tipping over into cytotoxicity. To explore this response more thoroughly, we selected an individual compound for further evaluation. QM0005928/DDD01293078 (QM5928) was chosen for this purpose for several reasons (Figure 6A). Firstly, the SAMP-Score dose profile followed this typical proliferation-senescence-toxicity trajectory (Figure 6B-C), being classed as senescent in both screens at 10uM but neither at 50uM, providing a range of phenotypes to explore. Secondly, the 10uM dose produced a final cell number very close to the seeding number, whilst the following dose strayed just below this threshold (Figure 6D). This provided two conditions that are very similar in cell number, but which SAMP-Score has been able to distinguish, as senescence and toxicity. This sensitivity despite minimal signal to noise in terms of cell count would address a major challenge in senescence screening^14^. Thirdly, the compound at several doses was classed by SAMP-Score and nuclear count as senescent but failed to meet the cell area Z-score threshold (Figure 6E-F). This provided an opportunity to explore a compound producing senescence phenotypes that our cluster-based labelling identified, which move beyond the classically reported “large cells”. Finally, the compound itself had a series of appealing properties including a low IC50, high solubility, favourable physiochemistry, low toxicity (HepG2) and a high lipophilic efficiency (Supplemental Figure 6A). In summary, it represented a versatile chemical tool for exploring pro-senescence in Sen-Mark+ cancer.

**Figure 6:**
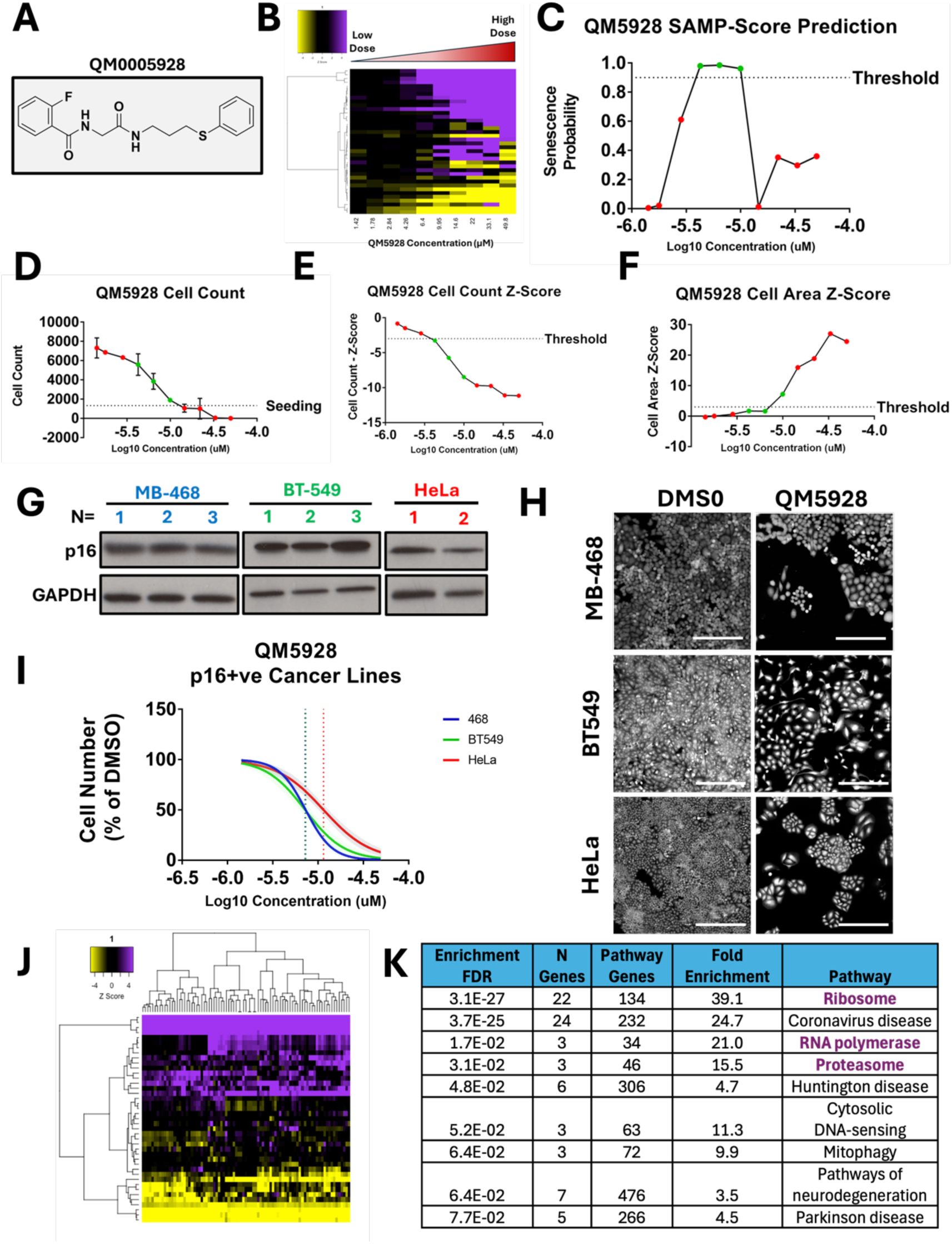
Validation of QM0005928 (QM5928) A: QM0005928 chemical structure. B: *Heatmap representing high content analysis feature (HCA; y-axis) profiles of a dose response of QM5928 (x-axis). C: Line Plot showing change in SAMP score with increasing doses of QM5928. D: Raw cell counts of increasing doses of QM5928 (Red = NonSen and Green = Sen SAMP-Score Classification). E-F: Z-scores for cell count and cell area increasing doses of QM5928. (Red = NonSen and Green = Sen SAMP-Score Classification). G: Western plot showing p16 expression in MB-468, BT549 and HeLa Sen-Mark+ cancer lines. H-I:* Cell counts for *MB-468, BT549 and HeLa Sen-Mark+ cancer lines in response to QM5928. N=3.* Scale bar = 100μm. *J: Heatmap representing high content analysis feature (HCA; y-axis) profiles of HeLa screen treatments that phenocopy QM5928 by hierarchical clustering. K: KEGG pathways analysis of HeLa screen treatments that phenocopy QM5928 by hierarchical clustering*.

As described above, validation of a senescence response in Sen-Mark+ cells is extremely challenging and was of course the motivation for our decision to take an unsupervised approach to senescence labelling when training SAMP-Score^10^. Indeed, as well as the characteristic expression of p16 in BLBC (Figure 6G) we also observed high levels of other canonical senescence hallmarks including high levels of SA-β-Gal, p21 and p53 (Supplemental Figure 6B-E), emphasising the classification challenge and the limitations of these markers more broadly^1^. Therefore, rather than characterising a wide range of senescence hallmarks, we sought to explore the efficacy of QM5928 in other Sen-Mark+ cancer lines (Figure 6H-I). We demonstrated that in both BT-549 (BLBC) and HeLa cells, QM5928 reduced cellular proliferation, in a dose manner similar to the MB-468s, indicating this compound has potential utility across a range of Sen-Mark+ contexts.

To further reinforce that the response being predicted by SAMP-Score is indeed senescence, we explored the SAMP-Score classification of BLBC cancer lines treated with compounds whose efficacy in senescence induction (or lack thereof) is well established. The CDK4/6 inhibitor palbociclib, has been previously demonstrated to be ineffective in cell lines already positive for p16 (whose effect the compound aims to mimic). This was observed in both p16 positive MB-468 and BT549 with no reduction in proliferation and a negative SAMP-Score classification (Supplemental Figure 7B-D). However, the p16-null MB-231 line (Supplemental Figure 7A) did respond canonically to palbociclib, with cell proliferation inhibited (Supplemental Figure 7E). Importantly, this treatment was classified as senescent by SAMP-Score, with a clear SAMP profile produced, that was not seen in the other lines (Supplemental Figure 7F-G). This emphasises the utility of the model across a range of cancer cell contexts and supports the principle that the phenotype being predicted is indeed senescence, demonstrating that SAMP-Score is a versatile tool for identifying senescence in cancer cells.

Finally, we utilised a technique that has previously been demonstrated to provide insight into potential mechanisms of actions of novel compounds by comparing the phenotypes elicited to a reference data set^18^. This is known as phenocopying and is based on the principle that perturbations in the same pathway by different methods will establish similar phenotypes. Here, we compared the morphology profile of QM5928 in the DR screen to the original siRNA screen. Utilising hierarchical clustering once more, we were able to identify the siRNAs that produced phenotypes comparable to that of QM5928 (Figure 6J). KEGG pathway analysis showed that QM5928 phenocopied pathways whose inhibition has been previously established to lead to senescence in cancer cells including the Ribosome^27,28^, RNA Polymerase^29^ and Proteasome^30,31^ (Figure 6K). Whilst this does not mean QM5928 acts via these mechanisms specifically, it does suggest that the response the compound is eliciting can be considered senescence and could also be an indication that targeting these pathways might be of therapeutic benefit in Sen-Mark+ cancers.

## Discussion

Sen-Mark+ cancer cells represent a particular challenge to identify the establishment of a senescence response, due to their intrinsically high levels of the most common hallmarks of senescence^10^. These cancers are often associated with particularly aggressive forms of the disease, where unmet clinical need is greatest, as in the case of BLBC. The tumour suppressive role of senescence makes re-instatement of a senescence programme (pro-senescence) in these Sen-Mark+ cells an attractive therapeutic strategy, but the challenge of identification has limited the application of standard screening approaches^1^. More broadly, it is becoming increasingly appreciated that senescence exists in contexts which do not conform to many established hallmarks, making the need for novel tools for senescence classification ever more important^16^.

Here, we utilised multiple ML approaches to develop SAMP-Score, a model that is able to readily identify induction of a senescence response in multiple Sen-Mark+ cancer cell lines and conditions. By utilising unbiased clustering algorithms, we have been able to establish a breadth of senescence morphology profiles by building on our previous observation of senescence-associated morphology profiles (SAMPs). Utilising these, we have developed a model that is able to distinguish senescence from two opposing ends of the cell fate spectrum – proliferation and cytotoxicity. This represents an advantage over traditional screening readouts, which typically rely on more rigid cutoffs to identify hits. By applying SAMP-Score to both diversity library and dose response screening data, we have been able to demonstrate its utility as a drug discovery tool through identification of QM5928, a novel pro-senescence compound in Sen-Mark+ cancer cells. This compound is effective at inducing senescence in a range of BLBC lines and phenocopies pathways previously linked to senescence induction. Future work will focus on understanding the mechanisms through which QM5928 is able to exert its pro-senescence effects, with a view to forming the “first-punch” of a paired pro-senescence-senolytic therapeutic approach.

## Conclusions

Overall, this study further demonstrates the utility of high-content morphological analysis as a tool for the identification of senescent cells. This is particularly powerful when paired with modern ML approaches, expanding the possible contexts into which pro-senescence drug discovery can be performed.

## Materials and Methods

### Code and Model Availability

All scripts, workflows, packages and models used to develop SAMP-Score are available at https://github.com/Phenotypic-Screening-QMUL/SAMP-Score. These are broken up into sequential steps and can be run from the R Markdown files, constituting ∼8,500 lines of code. To contextualise this code and the generated outputs, HTML guides have also been included (Supplemental Data 1) – these are provided as a line-by-line walkthrough of the workflow. Due to the large size of the single target high-content analysis screening data (Supplemental Data 2), this is available on reasonable request from the corresponding author. Files required to run scripts are included in Supplemental Data 1. All data analysis was performed in R version 4.2.1 using an x86_64-apple-darwin17.0 (64-bit) platform unless otherwise stated.

### Cell culture and reagents

Unless specified, all reagents were from Sigma, UK. HeLa (CRUK), MDA-MB-468 (ATCC, HTB-132) BT549 (ATCC, HTB-122) and MDA-MB-231 (ATCC, HTB-26) cells were cultured in high-glucose DMEM (Life Technologies, UK). HeLa medium was supplemented with 5% FBS (Labtech.com, UK), 1mM sodium pyruvate, and 2mM L-glutamine (Life Technologies, UK), while MDA-MB-468, BT549 and MDA-MB-231 medium contained 10% FBS, 1mM sodium pyruvate, and 2mM L-glutamine. Cells were maintained at 37°C/5% CO₂ without antibiotics. Immunoblotting was performed as previously described using antibodies against p16 (Santa Cruz, sc-56330; 1:100), p21 (Cell Signalling, 12D1; 1:2,000), p53 (Cell Signalling, 2527S; 1:1,000), ß-tubulin (EnoGene, E1C601-1; 1:20,000) and GAPDH (Abcam, ab9485; 1:5,000)^23^. All compounds were solubilised in 0.5% DMSO.

### Screening

#### Automation and Liquid Handling

Cell culture conditions (cell seeding, media changes etc.) were adapted for use with a CyBio Vario liquid handing robotics system, utilising a 384-tip set-up. This system was also used to perform fixation, permeabilization and staining procedures described below.

#### Genome Wide siRNA screen

HeLa cells were seeded in 384-well plates at 5,000 cells/cm^2^. Reverse transfection at 30nM was performed using a genome-wide siRNA library containing 1:1:1 ratio pools of 3 siRNAs (Ambion) using HiPerFect transfection reagent. Media was changed after 48hrs and cell fixation performed following another 72hrs. Negative control wells containing an siRNA targeting cyclophilin B (siGLO, D-001610-01, Dharmacon) were also included. These are found in columns 23 and 24. In total 21,658 siRNA conditions were tested (excluding controls) across 63 384-well plates which were screened in 3 batches.

#### Compound diversity library screen

A 10,000-compound diversity screening library was supplied for hit identification by the Dundee Drug Discovery Unit (DDU). The compounds represent novel chemical entities with no established target or mechanism of action and are identified by their QMCode designation (An alternative DDU designation may be found in the Supplemental Table S1). These compounds were solubilised in 0.5% DMSO vehicle, which also served as a negative control (located in columns 11, 12, 23 and 24). Compounds were screened at two doses - 10µM and 50µM. 10µl of each compound dilution was prepared per 384-well plate well and MDA-MB-468 cells seeded on top at 1,320 cells/well in 60µl of medium. Media was changed after 48hrs and cells fixed following a further 72hrs. In total, 60 384-well plates which were screened in 4 batches.

#### Compound dose response screen

A second compound screen containing 447 compounds in 10-point dose response curves was supplied by DDU. These compounds represent a subset of the original diversity screen but were not selected based on the current SAMP-Score methodology, but through stringent (≥3) Z-Score thresholds for nuclear count and cell area. Conditions, procedures and data processing was identical to the first screen with compounds being screened at the following doses: 49.80µM, 33.10µM, 22.00µM, 14.60µM, 9.95µM, 6.40µM, 4.26µM, 2.84µM, 1.78µM, 1.42 µM. The screen was performed in two identical batches representing two technical replicates for each compound/dose combination.

### Immunofluorescence staining and high content imaging

Immunofluorescence staining and high content analysis (HCA) microscopy has been described previously^17,23^. Briefly, fixation was carried out using 3.7% paraformaldehyde + 5% sucrose, with subsequent permeabilization using 0.1% Triton X-100. Nuclei and whole cells were stained for 2 hours at room with diamidino-2 phenylindole (DAPI) (Sigma UK, D8417, 1:1,000) and HCS Cell Mask Deep

Red (Thermo-Fisher UK, H32721, 1:100,000). High throughput automated imaging was then performed using an INCell Analyser 2200. For SA-β-Galactosidase activity assay, CellEvent Senescence Green Detection Kit (ThermoFisher, C10850) was used for 2hr at 37°C without CO2. Quantitation of cell positivity was set at the 95th-percentile threshold in the unstained control.

### High-content analysis (HCA) and Feature Selection

InCarta high-content image analysis software (Molecular Devices) was used to assess nuclear and cellular features; with object masks generated from DAPI and Cell Mask staining. Bespoke detection protocols were developed for all cell lines, but the same set of curated morphological features were assessed in each case (Supplemental Figure 1). This set of features consisted of a refined list based on our previous work; with the removal of those that were determined to be highly correlated according to a Pearson correlation index threshold of >0.9^17^. The input data into the correlation calculation was the morphological profiles from all individual siGlo control cells (∼1.16M cells) within the HeLa screen. The decision of which of the highly correlated features to retain was determined by the average overall correlation of both features to all other features within the dataset; with the feature with the highest score (and thus producing least variance on average) discarded.

#### Z-Score Data Processing

Z-score data scaling was performed for all high-content imaging features (Supplemental Figure 1) as previously reported^17^. This allows for both normalisation to a batch control condition (siGLO or DMSO) and data scaling across features. Z-Scores were calculated according to the following equation:

- *Z-Score **=** value of experimental condition (siRNA or Compound) – mean value of control condition / Standard Deviation (SD) of control condition*

Z-Scores for all high-content features were displayed as heatmaps; where black represented no-change relative to the control condition. Z-score changes of greater than 1.96 from the control condition (95% confidence level) in either positive or negative direction were visualised as purple or yellow respectively. For a condition where no change was observed relative to the control, an entirely black profile would be produced.

### Unsupervised Machine Learning - HeLa Screen Cluster Analysis for Senescence Labelling

Morphological Z-Score profiles from all siRNAs were assessed via unsupervised hierarchical clustering; with distance matrices constructed using Euclidean distances via the dist() function and clustering via the ward.D2 method argument within the hclust() function from the “stats” package. The number of clusters was determined through visual inspection of the dendrogram and heatmap generated via the heatmap.2 function within the “gplots” package. K-means clustering was performed using the kmeans() function from base R, with the number of centres iteratively altered upon data inspection. Clusters were overlaid on individual treatment morphological profiles, which in turn were visualised via uniform manifold approximation and projection (UMAP) dimensionality reduction from the “umap” package. Manual cluster labelling was then performed through assessment of heatmap profiles and cluster membership comparisons between the two methods made via Venn diagrams.

### Supervised Machine Learning – SAMP-Score Classification Model Development

#### Data Organisation

Data labelling of the HeLa screen siRNA treatments (Senescent vs Non-Senescent) was performed according to the unsupervised cluster-based approach. Data partitioning was then performed with 80% being allocated to the training dataset and 20% to the testing. To account for the large class imbalance between the Senescent (Sen) and Non-Senescent (NonSen) classes, the NonSen class was then randomly undersampled to produce an equal number of observations in each class. Those NonSen conditions that were removed were “recycled” into the testing dataset. Crucially, these data were never used for model training, preventing data leakage. The training data was then further split 50:50 into two separate training datasets. One set was used to train individual ML models and the other for “testing”, to produce prediction coefficients. These prediction values then became the training data variables for the **SAMP-Score** ensemble meta model. Each individual ML model was then used to assess the testing data (original 20% plus recycled) and individual model performance assessed. The prediction coefficients from these models were then combined into a testing dataset for the meta model to determine a final prediction. For the alternative model constructed using traditional screening thresholds (+/- 1.96 Z-score changing in nuclear count and cell area) the only change was at the data labelling stage.

#### Model Development

All models were constructed in R using the following packages: glmnet (logistic regression, Lasso regularisation and elastic nets); randomForest (random forest); e1071 (support vector machine; SVM); mda (multiple discriminant analysis; MDA); neuralnet (neural network; NN). The stacked meta model is a lasso regularisation model which takes the prediction coefficients from all seven models as input data. For the individual models, default hyperparameters were selected unless otherwise indicated and a decision threshold of 0.5 was applied. Further details of model parameters are found in the html guides (Supporting Data Set 2). Model performance was assessed via the generation of receiver operating characteristic (ROC) curves and assessment of model accuracy (proportion of correct predictions), Negative Predictive Value (NPV; proportion of true negatives out of all negative predictions), recall (proportion of true positives out of all positives), F1 score (harmonic mean of PPV and recall), AUC (Area Under ROC Curve) and positive prediction value (PPV; proportion of true positives among positive predictions). Prediction rates were recorded as confusion matrices.

#### Phenocopying

Z-Score morphology profiles from the HeLa screen were combined with that of QM5928 from the dose response screen. The dose selected for QM5928 was the highest that scored as a senescence hit according to SAMP-Score (9.95µM), in order to explore the most well-established phenotype. Hierarchical clustering was then performed with the data partitioned into 50 clusters, in order to refine the number of treatments within individual clusters. Targets of the 77 siRNA treatments that produced phenotypes that clustered alongside QM5928 (Supplemental Data 3) were then assessed by KEGG pathway analysis using ShinyGO 0.81^24^.

## Statistical analysis

Statistical analysis was performed using GraphPad Prism 7. An unpaired Student’s t-test was used to compare the means of two groups unless specified. Data ≥2 independent experiments unless otherwise stated. Error bars represent SD. Scale bars = 250µm.

## Declarations

### Ethics approval and consent to participate

Not applicable.

### Consent for publication

Not applicable.

### Availability of data and materials

All packages and versions used in this code are available at the following URL: https://github.com/Phenotypic-Screening-QMUL/SAMP-Score (Note: this will be made available upon acceptance, and we can provide access to reviewers with pleasure).

### Authors’ contributions

RW and CLB developed the initial concept. RW, BKH, MM, EO’S, LCM, LG, AH, FB, CM, CG, DG and CLB finalised the concept, developing and designed experiments. RW and CLB developed SAMP-Score. RW, BKH, MM, EO’S, LCM, LG, AH, FB performed all experiments. RW and CLB wrote the initial manuscript draft with all authors contributing to revisions of manuscript prior to submission. All authors (with the exception of AH) read and approved the final manuscript.

## Supporting information

HTML Walkthrough Guides for Code

Phenocopying Cluster Lists

QM and DDU Compound Codes

## Acknowledgements

Dedicated to the memory of Anthony (Tony) Hope – a great scientist and friend.

## Authors’ information (optional)

Not applicable.

