## Supplemental Figures for "SAMP-Score: A morphology-based machine learning classification method for screening pro-senescence compounds in p16 positive cancer cells"

### Supplemental Information

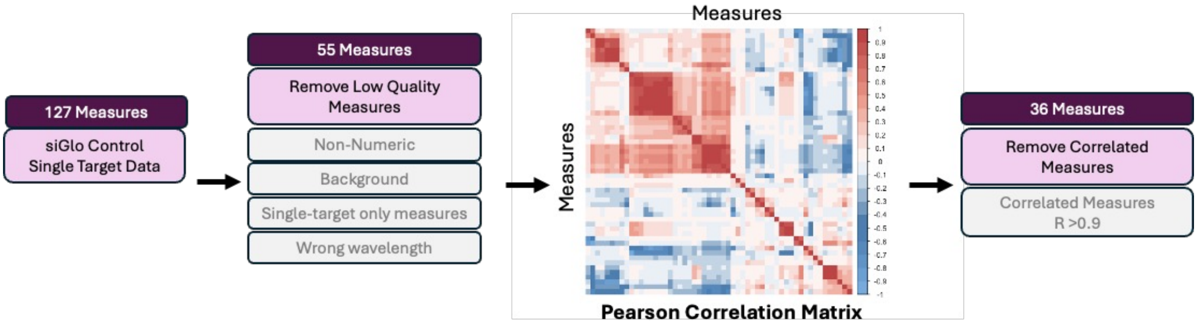


| Measures Retained | | | | | | | | Measures Removed | | | |
|---|---|---|---|---|---|---|---|---|---|---|---|
| 1 | Nuclei.Form.Factor | 11 | Nuclei.Neighbor.Count..SOI.. | 21 | Cells.Area | 31 | Cells.Max.Intensity | 1 | Cells.Gyration.Radius | 11 | Nuclei.Entropy |
| 2 | Nuclei.Elongation | 12 | Nuclei.Spacing..MIN.. | 22 | Cells.Form.Factor. | 32 | Cells.Major.Axis | 2 | Cells.Diameter | 12 | Cells.Entropy |
| 3 | Nuclei.Compactness | 13 | Nuclei.Neighbor.Count..MIN. | 23 | Cells.Elongation | 33 | Cells.Minor.Axis | 3 | Cells.Perimeter | 13 | Cells.Intensity.CV..Cell.. |
| 4 | Nuclei.Chord.Ratio | 14 | Nuclei.Spacing..Gabriel.. | 24 | Cells.Compactness | 34 | Cells.Skewness | 4 | Cells.Light.Flux. | 14 | Cells.Nuc.Cyto.Intensity. |
| 5 | Nuclei.Displacement | 15 | Nuclei.Neighbor.Count..Gabriel.. | 25 | Cells.Chord.Ratio. | 35 | Cells.Kurtosis | 5 | Nuclei.Light.Flux. | 15 | Cells.Total.Intensity..Cell.. |
| 6 | Nuclei.Intensity | 16 | Nuclei.Spacing..Lune.. | 26 | Cells.Nuc.Cell.Area | 36 | Cells.Energy | 6 | Nuclei.Diameter | 16 | Cells.Intensity.SD..Cell. |
| 7 | Nuclei.Total.Intensity | 17 | Nuclei.Neighbor.Count..Lune.. | 27 | Cells.Total.Intensity..Cyto.. | | | 7 | Nuclei.Major.Axis | 17 | Cells.Intensity..Cyto. |
| 8 | Nuclei.Intensity.CV | 18 | Nuclei.Skewness | 28 | Cells.Intensity.CV..Cyto.. | | | 8 | Nuclei.Gyration.Radius | 18 | Cells.Intensity..Cell. |
| 9 | Nuclei.Major.Axis.Angle | 19 | Nuclei.Kurtosis | 29 | Cells.Intensity.Spreading. | | | 9 | Nuclei.Perimeter | 19 | Nuclei.Intensity.SD |
| 10 | Nuclei.Spacing..SOI.. | 20 | Nuclei.Energy | 30 | Cells.Intensity.SD..Cyto. | | | 10 | Nuclei.Area | | |

*Supplemental Figure 1: Feature reduction via person correlation assessment*

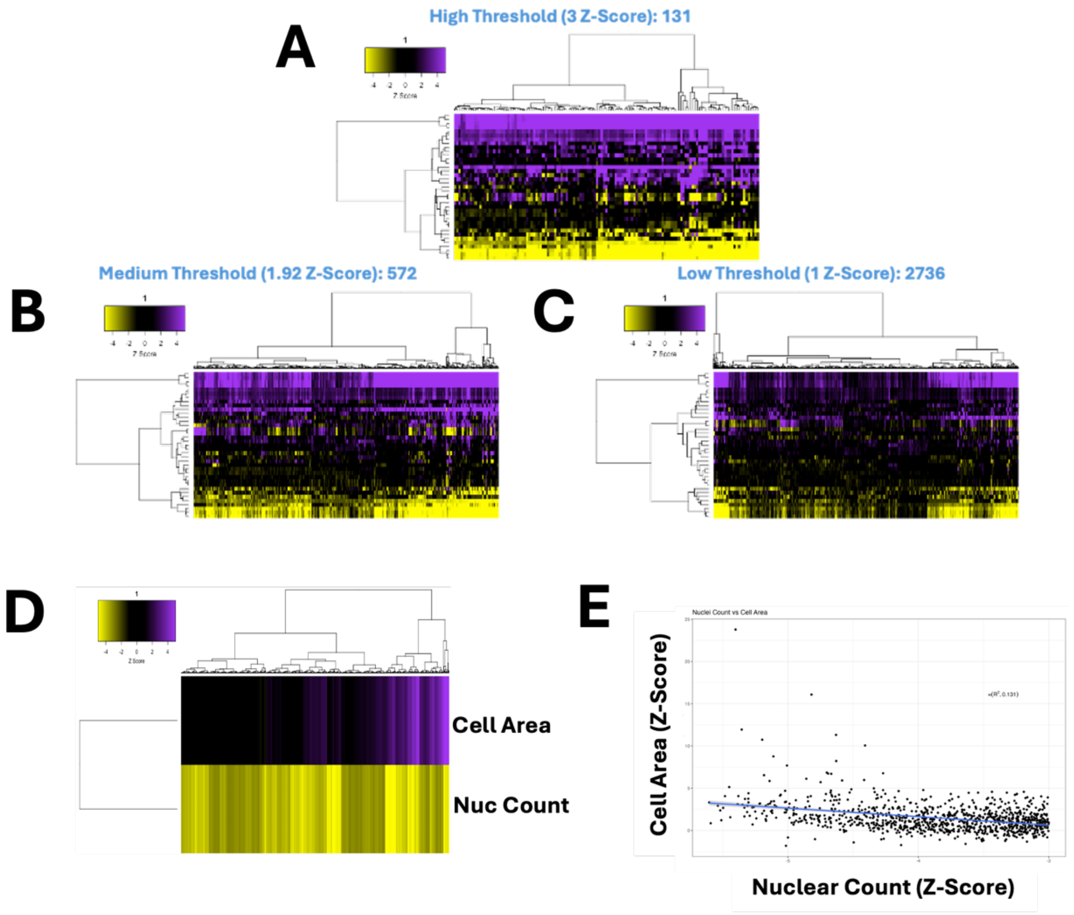


***Supplemental Figure 2: Genome-wide siRNA HeLa Screen – Traditional Screening Readouts***

*A-C: Heatmaps representing high content analysis feature (HCA; y-axis) profiles of a genome-wide siRNA screen treatments (siRNAs; x-axis) that reduce nuclear count and increase nuclear area by high, medium and low stringency thresholds. D: Heatmap profile showing cell area and nuclear count Z-scores for all treatments. E: Scatter plot showing nuclear count vs cell area Z-scores. In all heatmaps, purple indicates positive modulation and yellow negative modulation of greater than 1.96 Z-scores from siGlo control. Black indicates no change.*

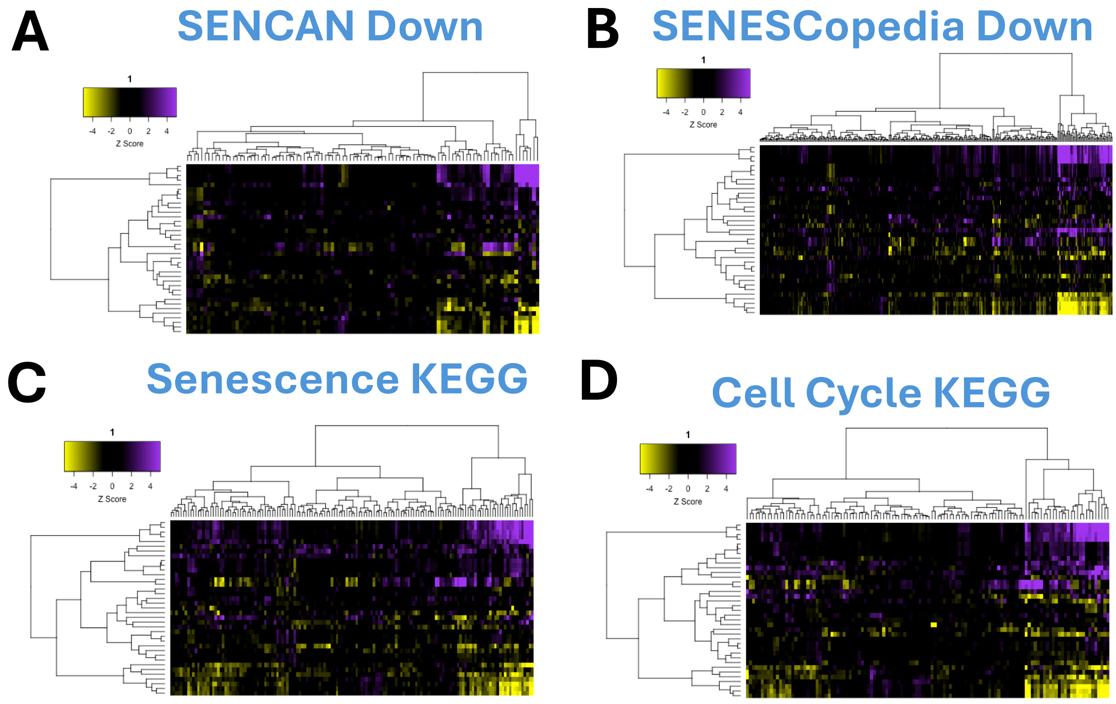


***Supplemental Figure 3: Genome-wide siRNA HeLa Screen – Biased Pathway Labelling***

*A-D: Heatmaps representing high content analysis feature (HCA; y-axis) profiles of a genome-wide siRNA screen treatments (siRNAs; x-axis) selected through identification as downregulated in SENCAN and SENESCopedia database or Senescence/Cell Cycle KEGG pathway analysis terms. In all heatmaps, purple indicates positive modulation and yellow negative modulation of greater than 1.96 Z-scores from DMSO vehicle control. Black indicates no change.*

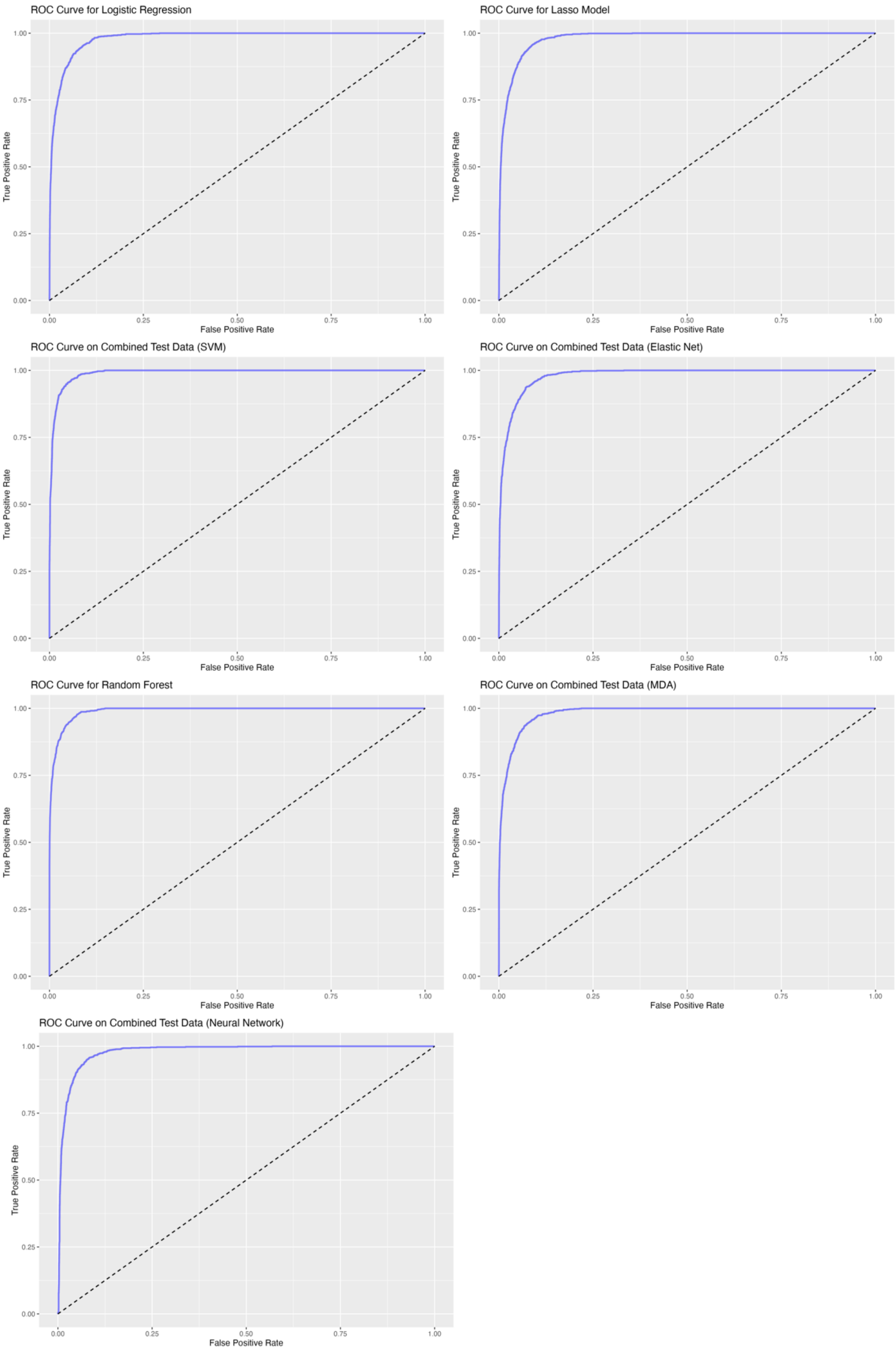


***Supplemental Figure 4:*** **Receiver operating characteristic (ROC)** ***curves for all composite models of SAMP-Score***

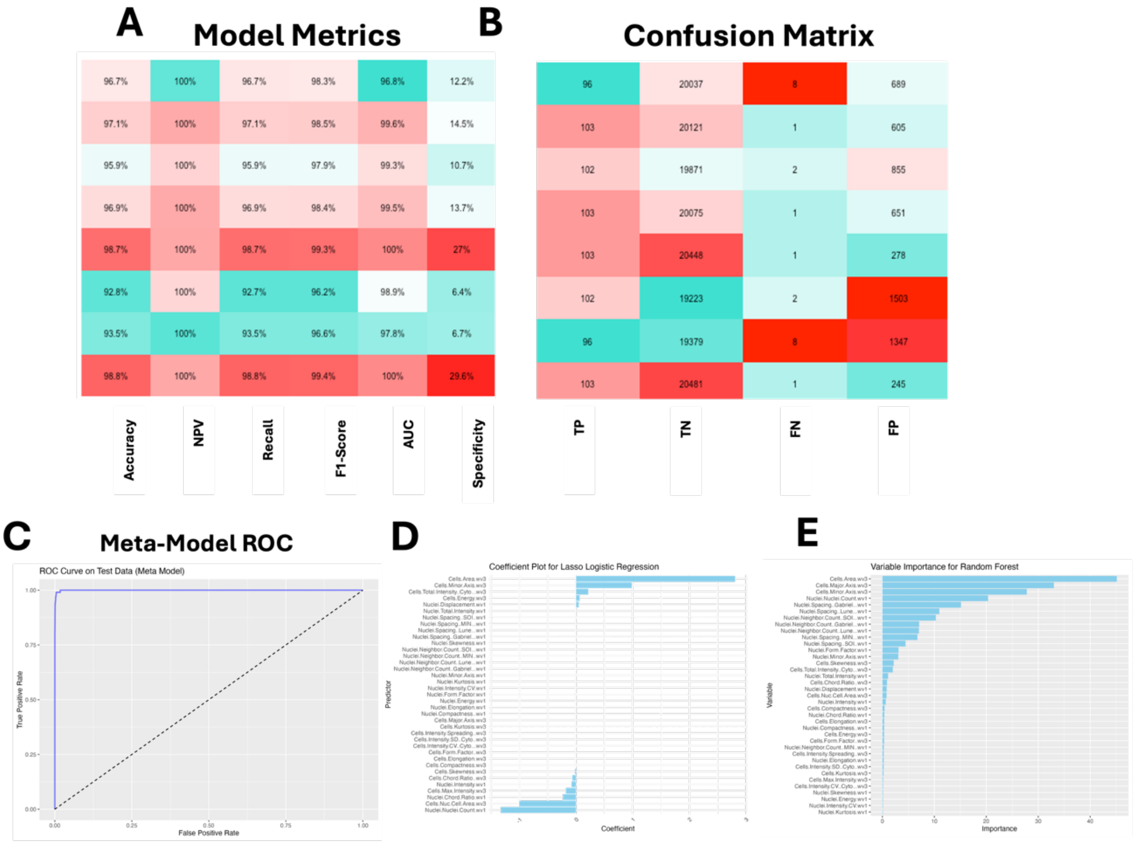


***Supplemental Figure 5:*** **Model Development and Metric Assessment using traditional threshold-based senescence labelling of training data**

*A-B: Machine learning (ML) model assessment metrics and confusion matrix for all individual ML models and stacked meta-model (SAMP-Score). NPV: Negative Prediction Value, AUC: Area under curve, PPV: Positive Prediction Value, TP: True Positive, TN: True Negative, FN: False Negative, FP: False Positive. C: Receiver operating characteristic (ROC) curve for stacked meta model. D-E: Model feature contributions to Lasso and Random Forest models.*

| A | | QM0005928 |
|---|---|---|
| | | 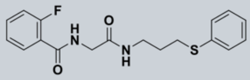 |
| Toxicity | HepG2 pIC50 | **4.4** |
| Solubility | ReaSol (uM) | **180** |
| DMPK | Clearance (ml/min/g) | **8.3** |
| | MDCK (Papp, nm/s) | **235** |
| Physiochemical Properties | MWt | **346.42** |
| | logP/CHI-logD | **2.6/2.6** |
| | tPSA | **58** |
| Metrics | LE/LLE | **0.30/ 2.6** |
| | IFI | **0.1** |

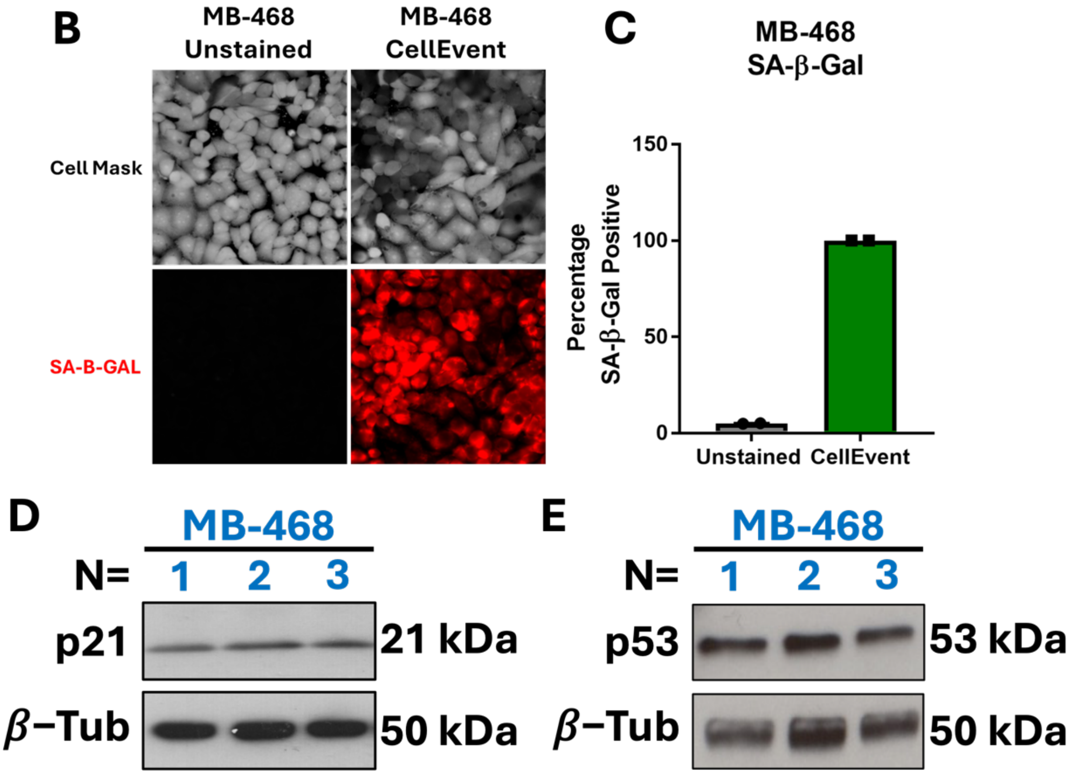


***Supplemental Figure 6: QM0005928 physicochemical properties and SenMark+ cancer cell senescence markers.***

*A: Standard chemical assessment panel for QM5928. B-C: Senescence-associated beta galactosidase staining of MB-468 cells. N=1 with 2 technical replicates. D-E: Immunoblotting of MB-468 cells for p21 and p53. N=3.*

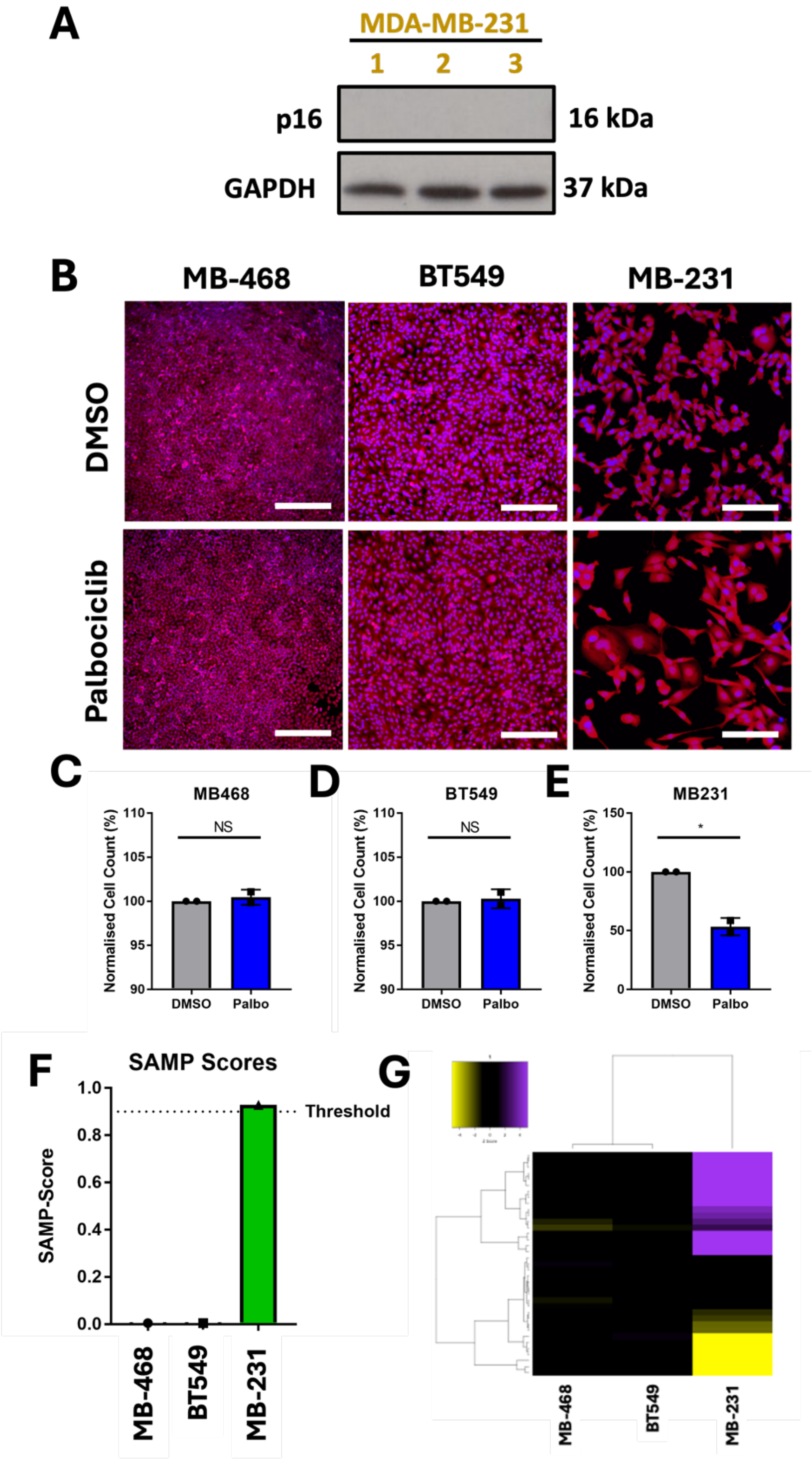


***Supplemental Figure 7:*** **SAMP-Score Assessment of BLBC Response to CDK Inhibition**

*A:* Immunoblotting of MB-231 cells for p16. N-3. B-E: Response to 1uM palbociclib treatment or vehicle control (DMSO) in MB-468, BT549 and MB-231 BLBC lines. F: SAMP-Score classification coefficient for MB-468, BT549 and MB-231 BLBC lines treated with palbociclib. G: *Heatmap representing high content analysis feature (HCA; y-axis) profiles of* MB-468, BT549 and MB-231 BLBC lines treated with palbociclib *(x-axis). In all heatmaps, purple indicates positive modulation and yellow negative modulation of greater than 1.96 Z-scores from DMSO vehicle control. Black indicates no change.*
