## Supplementary material for "SAMP-Score: A morphology-based machine learning classification method for screening pro-senescence compounds in p16 positive cancer cells": HTML Walkthrough Guides for Code: Supplemental Data 1_0_Feature_Selection_Correlations.html


### 0\_Feature\_Selection\_Correlations

###### Ryan Wallis

#### 2024-11-20

```
library("gplots")
```

```
## 
## Attaching package: 'gplots'
```

```
## The following object is masked from 'package:stats':
## 
##     lowess
```

```
library("RColorBrewer")
library("matrixStats")
library("plyr")
```

```
## 
## Attaching package: 'plyr'
```

```
## The following object is masked from 'package:matrixStats':
## 
##     count
```

```
library("dplyr")
```

```
## 
## Attaching package: 'dplyr'
```

```
## The following objects are masked from 'package:plyr':
## 
##     arrange, count, desc, failwith, id, mutate, rename, summarise,
##     summarize
```

```
## The following object is masked from 'package:matrixStats':
## 
##     count
```

```
## The following objects are masked from 'package:stats':
## 
##     filter, lag
```

```
## The following objects are masked from 'package:base':
## 
##     intersect, setdiff, setequal, union
```

```
library("data.table")
```

```
## 
## Attaching package: 'data.table'
```

```
## The following objects are masked from 'package:dplyr':
## 
##     between, first, last
```

```
library("stringr")
library("ggplot2")
library("Rtsne")


#Import siGlo Control ST Data
All_ST_Clean <- read.csv("All_ST_Clean.csv") # Note this takes a long time to run

#Remove Final Low Quality Measures
numdata <- All_ST_Clean [,-c(1,2,17,18,19,21,56,57,60)]
numdatadf <- data.frame(numdata)
sapply(numdatadf,is.numeric)
```

```
##                      Nuclei.Area.wv1               Nuclei.Form.Factor.wv1 
##                                 TRUE                                 TRUE 
##                Nuclei.Elongation.wv1              Nuclei.Compactness..wv1 
##                                 TRUE                                 TRUE 
##               Nuclei.Chord.Ratio.wv1           Nuclei.Gyration.Radius.wv1 
##                                 TRUE                                 TRUE 
##              Nuclei.Displacement.wv1                  Nuclei.Diameter.wv1 
##                                 TRUE                                 TRUE 
##                 Nuclei.Perimeter.wv1                 Nuclei.Intensity.wv1 
##                                 TRUE                                 TRUE 
##           Nuclei.Total.Intensity.wv1              Nuclei.Intensity.CV.wv1 
##                                 TRUE                                 TRUE 
##               Nuclei.Light.Flux..wv1              Nuclei.Intensity.SD.wv1 
##                                 TRUE                                 TRUE 
##                Nuclei.Major.Axis.wv1          Nuclei.Major.Axis.Angle.wv1 
##                                 TRUE                                 TRUE 
##             Nuclei.Spacing..SOI..wv1     Nuclei.Neighbor.Count..SOI...wv1 
##                                 TRUE                                 TRUE 
##            Nuclei.Spacing..MIN...wv1      Nuclei.Neighbor.Count..MIN..wv1 
##                                 TRUE                                 TRUE 
##        Nuclei.Spacing..Gabriel...wv1 Nuclei.Neighbor.Count..Gabriel...wv1 
##                                 TRUE                                 TRUE 
##           Nuclei.Spacing..Lune...wv1    Nuclei.Neighbor.Count..Lune...wv1 
##                                 TRUE                                 TRUE 
##                  Nuclei.Skewness.wv1                  Nuclei.Kurtosis.wv1 
##                                 TRUE                                 TRUE 
##                    Nuclei.Energy.wv1                   Nuclei.Entropy.wv1 
##                                 TRUE                                 TRUE 
##                       Cells.Area.wv3               Cells.Form.Factor..wv3 
##                                 TRUE                                 TRUE 
##                 Cells.Elongation.wv3                Cells.Compactness.wv3 
##                                 TRUE                                 TRUE 
##               Cells.Chord.Ratio..wv3           Cells.Gyration.Radius..wv3 
##                                 TRUE                                 TRUE 
##              Cells.Nuc.Cell.Area.wv3                   Cells.Diameter.wv3 
##                                 TRUE                                 TRUE 
##                  Cells.Perimeter.wv3           Cells.Intensity..Cell..wv3 
##                                 TRUE                                 TRUE 
##           Cells.Intensity..Cyto..wv3    Cells.Total.Intensity..Cell...wv3 
##                                 TRUE                                 TRUE 
##    Cells.Total.Intensity..Cyto...wv3       Cells.Intensity.CV..Cell...wv3 
##                                 TRUE                                 TRUE 
##       Cells.Intensity.CV..Cyto...wv3       Cells.Intensity.Spreading..wv3 
##                                 TRUE                                 TRUE 
##                Cells.Light.Flux..wv3        Cells.Nuc.Cyto.Intensity..wv3 
##                                 TRUE                                 TRUE 
##        Cells.Intensity.SD..Cell..wv3        Cells.Intensity.SD..Cyto..wv3 
##                                 TRUE                                 TRUE 
##              Cells.Max.Intensity.wv3                 Cells.Major.Axis.wv3 
##                                 TRUE                                 TRUE 
##                 Cells.Minor.Axis.wv3                   Cells.Skewness.wv3 
##                                 TRUE                                 TRUE 
##                   Cells.Kurtosis.wv3                     Cells.Energy.wv3 
##                                 TRUE                                 TRUE 
##                    Cells.Entropy.wv3 
##                                 TRUE
```

```
#Generate Correlation Matrix 

library(corrplot)
```

```
## corrplot 0.92 loaded
```

```
col <- colorRampPalette(c( "#77AADD", "#4477AA", "#FFFFFF", "#EE9988","#BB4444" ))
df_cor <- cor(numdatadf, use=c("pairwise.complete.obs")) # This will also take some time to run 
#makes labels fit in margins
par(xpd=TRUE)
corrplot(df_cor, method="color", col=col(20),  
         type="full", order="hclust", tl.cex = 0.01, # Add coefficient of correlation
         tl.col="black", tl.srt=90, #Text label color and rotation
         # hide correlation coefficient on the principal diagonal
         diag=T,mar = c(2, 0, 1, 0))
```

```
#Generate Correlation Tables
library("caret")
```

```
## Loading required package: lattice
```

```
High_corr <- findCorrelation(df_cor, cutoff = .9, verbose = TRUE, names = TRUE)
```

```
## Compare row 34  and column  36 with corr  0.99 
##   Means:  0.376 vs 0.205 so flagging column 34 
## Compare row 36  and column  37 with corr  0.975 
##   Means:  0.364 vs 0.199 so flagging column 36 
## Compare row 37  and column  50 with corr  0.948 
##   Means:  0.347 vs 0.193 so flagging column 37 
## Compare row 45  and column  13 with corr  0.901 
##   Means:  0.327 vs 0.188 so flagging column 45 
## Compare row 13  and column  6 with corr  0.904 
##   Means:  0.287 vs 0.182 so flagging column 13 
## Compare row 8  and column  15 with corr  0.975 
##   Means:  0.272 vs 0.178 so flagging column 8 
## Compare row 15  and column  6 with corr  0.969 
##   Means:  0.258 vs 0.175 so flagging column 15 
## Compare row 6  and column  9 with corr  0.995 
##   Means:  0.238 vs 0.171 so flagging column 6 
## Compare row 9  and column  1 with corr  0.978 
##   Means:  0.219 vs 0.169 so flagging column 9 
## Compare row 1  and column  11 with corr  0.916 
##   Means:  0.193 vs 0.167 so flagging column 1 
## Compare row 28  and column  27 with corr  0.951 
##   Means:  0.184 vs 0.167 so flagging column 28 
## Compare row 55  and column  54 with corr  0.954 
##   Means:  0.254 vs 0.165 so flagging column 55 
## Compare row 42  and column  46 with corr  0.959 
##   Means:  0.255 vs 0.16 so flagging column 42 
## Compare row 46  and column  43 with corr  0.91 
##   Means:  0.23 vs 0.155 so flagging column 46 
## Compare row 40  and column  41 with corr  0.955 
##   Means:  0.208 vs 0.153 so flagging column 40 
## Compare row 47  and column  48 with corr  0.962 
##   Means:  0.2 vs 0.15 so flagging column 47 
## Compare row 39  and column  38 with corr  0.998 
##   Means:  0.182 vs 0.147 so flagging column 39 
## Compare row 38  and column  49 with corr  0.978 
##   Means:  0.156 vs 0.147 so flagging column 38 
## Compare row 12  and column  14 with corr  0.963 
##   Means:  0.119 vs 0.146 so flagging column 14 
## All correlations <= 0.9
```

```
hc = findCorrelation(df_cor, cutoff=0.9) # putt any value as a "cutoff" 
hc = sort(hc)
reduced_Data = df_cor[,-c(hc)]
reduced_Data <- as.data.frame(reduced_Data)

#Full Correlation Matrix
write.csv(df_cor, "Correlation Matrix.csv")
#Retained Features Matrix - Columns 
write.csv(reduced_Data, "Reduced Data.csv")
#Removed Features
write.csv(High_corr, "Correlated Measures.csv")
```
