## Supplementary material for "SAMP-Score: A morphology-based machine learning classification method for screening pro-senescence compounds in p16 positive cancer cells": HTML Walkthrough Guides for Code: Supplemental Data 1_1_Data_Wrangling_HeLa_Screen.html

Data\_Wrangling\_HeLa\_Screen.knit


### 1. Import Data

```
# Load the libraries 
library("gplots")
```

```
## 
## Attaching package: 'gplots'
```

```
## The following object is masked from 'package:stats':
## 
##     lowess
```

```
library("RColorBrewer")
library("dplyr")
```

```
## 
## Attaching package: 'dplyr'
```

```
## The following objects are masked from 'package:stats':
## 
##     filter, lag
```

```
## The following objects are masked from 'package:base':
## 
##     intersect, setdiff, setequal, union
```

```
library("matrixStats")
```

```
## 
## Attaching package: 'matrixStats'
```

```
## The following object is masked from 'package:dplyr':
## 
##     count
```

```
# Upload files - remove BOM
D001 <- read.csv("D01.csv",fileEncoding = 'UTF-8-BOM')
D002 <- read.csv("D02.csv",fileEncoding = 'UTF-8-BOM')
D003 <- read.csv("D03.csv",fileEncoding = 'UTF-8-BOM')
D004 <- read.csv("D04.csv",fileEncoding = 'UTF-8-BOM')
D005 <- read.csv("D05.csv",fileEncoding = 'UTF-8-BOM')
D006 <- read.csv("D06.csv",fileEncoding = 'UTF-8-BOM')
D007 <- read.csv("D07.csv",fileEncoding = 'UTF-8-BOM')
D008 <- read.csv("D08.csv",fileEncoding = 'UTF-8-BOM')
D009 <- read.csv("D09.csv",fileEncoding = 'UTF-8-BOM')
D010 <- read.csv("D10.csv",fileEncoding = 'UTF-8-BOM')
D011 <- read.csv("D11.csv",fileEncoding = 'UTF-8-BOM')
D012 <- read.csv("D12.csv",fileEncoding = 'UTF-8-BOM')
D013 <- read.csv("D13.csv",fileEncoding = 'UTF-8-BOM')
D014 <- read.csv("D14.csv",fileEncoding = 'UTF-8-BOM')
D015 <- read.csv("D15.csv",fileEncoding = 'UTF-8-BOM')
D016 <- read.csv("D16.csv",fileEncoding = 'UTF-8-BOM')
G001 <- read.csv("G001.csv",fileEncoding = 'UTF-8-BOM')
G002 <- read.csv("G002.csv",fileEncoding = 'UTF-8-BOM')
G003 <- read.csv("G003.csv",fileEncoding = 'UTF-8-BOM')
G004 <- read.csv("G004.csv",fileEncoding = 'UTF-8-BOM')
G005 <- read.csv("G005.csv",fileEncoding = 'UTF-8-BOM')
G006 <- read.csv("G006.csv",fileEncoding = 'UTF-8-BOM')
G007 <- read.csv("G007.csv",fileEncoding = 'UTF-8-BOM')
G008 <- read.csv("G008.csv",fileEncoding = 'UTF-8-BOM')
G009 <- read.csv("G009.csv",fileEncoding = 'UTF-8-BOM')
G010 <- read.csv("G010.csv",fileEncoding = 'UTF-8-BOM')
G011 <- read.csv("G011.csv",fileEncoding = 'UTF-8-BOM')
G012 <- read.csv("G012.csv",fileEncoding = 'UTF-8-BOM')
G013 <- read.csv("G013.csv",fileEncoding = 'UTF-8-BOM')
G014 <- read.csv("G014.csv",fileEncoding = 'UTF-8-BOM')
G015 <- read.csv("G015.csv",fileEncoding = 'UTF-8-BOM')
G016 <- read.csv("G016.csv",fileEncoding = 'UTF-8-BOM')
G017 <- read.csv("G017.csv",fileEncoding = 'UTF-8-BOM')
G018 <- read.csv("G018.csv",fileEncoding = 'UTF-8-BOM')
G019 <- read.csv("G019.csv",fileEncoding = 'UTF-8-BOM')
G020 <- read.csv("G020.csv",fileEncoding = 'UTF-8-BOM')
G021 <- read.csv("G021.csv",fileEncoding = 'UTF-8-BOM')
G022 <- read.csv("G022.csv",fileEncoding = 'UTF-8-BOM')
G023 <- read.csv("G023.csv",fileEncoding = 'UTF-8-BOM')
G024 <- read.csv("G024.csv",fileEncoding = 'UTF-8-BOM')
G025 <- read.csv("G025.csv",fileEncoding = 'UTF-8-BOM')
G026 <- read.csv("G026.csv",fileEncoding = 'UTF-8-BOM')
G027 <- read.csv("G027.csv",fileEncoding = 'UTF-8-BOM')
G028 <- read.csv("G028.csv",fileEncoding = 'UTF-8-BOM')
G029 <- read.csv("G029.csv",fileEncoding = 'UTF-8-BOM')
G030 <- read.csv("G030.csv",fileEncoding = 'UTF-8-BOM')
G031 <- read.csv("G031.csv",fileEncoding = 'UTF-8-BOM')
G032 <- read.csv("G032.csv",fileEncoding = 'UTF-8-BOM')
G033 <- read.csv("G033.csv",fileEncoding = 'UTF-8-BOM')
G034 <- read.csv("G034.csv",fileEncoding = 'UTF-8-BOM')
G035 <- read.csv("G035.csv",fileEncoding = 'UTF-8-BOM')
G036 <- read.csv("G036.csv",fileEncoding = 'UTF-8-BOM')
G037 <- read.csv("G037.csv",fileEncoding = 'UTF-8-BOM')
G038 <- read.csv("G038.csv",fileEncoding = 'UTF-8-BOM')
G039 <- read.csv("G039.csv",fileEncoding = 'UTF-8-BOM')
G040 <- read.csv("G040.csv",fileEncoding = 'UTF-8-BOM')
G041 <- read.csv("G041.csv",fileEncoding = 'UTF-8-BOM')
G042 <- read.csv("G042.csv",fileEncoding = 'UTF-8-BOM')
G043 <- read.csv("G043.csv",fileEncoding = 'UTF-8-BOM')
G044 <- read.csv("G044.csv",fileEncoding = 'UTF-8-BOM')
G045 <- read.csv("G045.csv",fileEncoding = 'UTF-8-BOM')
G046 <- read.csv("G046.csv",fileEncoding = 'UTF-8-BOM')
G047 <- read.csv("G047.csv",fileEncoding = 'UTF-8-BOM')
```

### 2.Tidy Measures

```
#Column Number for Measures -SAMP Measures plus Nuclei Count
column_measures <- c(3,6,8,9,10,11,13,16,17,18,29,31:41,58:62,64,70,72,73,86,87,90:91,93:95)

#Select only measures in each column 
D001 <- D001 %>%
  select(column_measures)
```

```
## Warning: Using an external vector in selections was deprecated in tidyselect 1.1.0.
## ℹ Please use `all_of()` or `any_of()` instead.
##   # Was:
##   data %>% select(column_measures)
## 
##   # Now:
##   data %>% select(all_of(column_measures))
## 
## See <https://tidyselect.r-lib.org/reference/faq-external-vector.html>.
## This warning is displayed once every 8 hours.
## Call `lifecycle::last_lifecycle_warnings()` to see where this warning was
## generated.
```

```
D002 <- D002 %>%
  select(column_measures)
D003 <- D003 %>%
  select(column_measures)
D004 <- D004 %>%
  select(column_measures)
D005 <- D005 %>%
  select(column_measures)
D006 <- D006 %>%
  select(column_measures)
D007 <- D007 %>%
  select(column_measures)
D008 <- D008 %>%
  select(column_measures)
D009 <- D009 %>%
  select(column_measures)
D010 <- D010 %>%
  select(column_measures)
D011 <- D011 %>%
  select(column_measures)
D012 <- D012 %>%
  select(column_measures)
D013 <- D013 %>%
  select(column_measures)
D014 <- D014 %>%
  select(column_measures)
D015 <- D015 %>%
  select(column_measures)
D016 <- D016%>%
  select(column_measures)
G001 <- G001 %>%
  select(column_measures)
G002 <- G002 %>%
  select(column_measures)
G003 <- G003 %>%
  select(column_measures)
G004 <- G004 %>%
  select(column_measures)
G005 <- G005 %>%
  select(column_measures)
G006 <- G006 %>%
  select(column_measures)
G007 <- G007 %>%
  select(column_measures)
G008 <- G008 %>%
  select(column_measures)
G009 <- G009 %>%
  select(column_measures)
G010 <- G010 %>%
  select(column_measures)
G011 <- G011 %>%
  select(column_measures)
G012 <- G012 %>%
  select(column_measures)
G013 <- G013 %>%
  select(column_measures)
G014 <- G014 %>%
  select(column_measures)
G015 <- G015 %>%
  select(column_measures)
G016 <- G016 %>%
  select(column_measures)
G017 <- G017 %>%
  select(column_measures)
G018 <- G018 %>%
  select(column_measures)
G019 <- G019 %>%
  select(column_measures)
G020 <- G020 %>%
  select(column_measures)
G021 <- G021 %>%
  select(column_measures)
G022 <- G022 %>%
  select(column_measures)
G023 <- G023 %>%
  select(column_measures)
G024 <- G025 %>%
  select(column_measures)
G025 <- G025 %>%
  select(column_measures)
G026 <- G026 %>%
  select(column_measures)
G027 <- G027 %>%
  select(column_measures)
G028 <- G028 %>%
  select(column_measures)
G029 <- G029 %>%
  select(column_measures)
G030 <- G030 %>%
  select(column_measures)
G031 <- G031 %>%
  select(column_measures)
G032 <- G032 %>%
  select(column_measures)
G033 <- G033 %>%
  select(column_measures)
G034 <- G034 %>%
  select(column_measures)
G035 <- G035 %>%
  select(column_measures)
G036 <- G036 %>%
  select(column_measures)
G037 <- G037 %>%
  select(column_measures)
G038 <- G038 %>%
  select(column_measures)
G039 <- G039 %>%
  select(column_measures)
G040 <- G040 %>%
  select(column_measures)
G041 <- G041 %>%
  select(column_measures)
G042 <- G042 %>%
  select(column_measures)
G043 <- G043 %>%
  select(column_measures)
G044 <- G044 %>%
  select(column_measures)
G045 <- G045 %>%
  select(column_measures)
G046 <- G046 %>%
  select(column_measures)
G047 <- G047 %>%
  select(column_measures)
```

### 3. Extract siGLO Controls

```
#Filter for siGlo control wells
D001_siGlo <- D001 %>%
  filter(WELL.LABEL %in% c("A - 23","B - 23","C - 23","D - 23"))
D002_siGlo <- D002 %>%
  filter(WELL.LABEL %in% c("A - 23","B - 23","C - 23","D - 23"))
D003_siGlo <- D003 %>%
  filter(WELL.LABEL %in% c("A - 23","B - 23","C - 23","D - 23"))
D004_siGlo <- D004 %>%
  filter(WELL.LABEL %in% c("A - 23","B - 23","C - 23","D - 23"))
D005_siGlo <- D005 %>%
  filter(WELL.LABEL %in% c("A - 23","B - 23","C - 23","D - 23"))
D006_siGlo <- D006 %>%
  filter(WELL.LABEL %in% c("A - 23","B - 23","C - 23","D - 23"))
D007_siGlo <- D007 %>%
  filter(WELL.LABEL %in% c("A - 23","B - 23","C - 23","D - 23"))
D008_siGlo <- D008 %>%
  filter(WELL.LABEL %in% c("A - 23","B - 23","C - 23","D - 23"))
D009_siGlo <- D009 %>%
  filter(WELL.LABEL %in% c("A - 23","B - 23","C - 23","D - 23"))
D010_siGlo <- D010 %>%
  filter(WELL.LABEL %in% c("A - 23","B - 23","C - 23","D - 23"))
D011_siGlo <- D011 %>%
  filter(WELL.LABEL %in% c("A - 23","B - 23","C - 23","D - 23"))
D012_siGlo <- D012 %>%
  filter(WELL.LABEL %in% c("A - 23","B - 23","C - 23","D - 23"))
D013_siGlo <- D013 %>%
  filter(WELL.LABEL %in% c("A - 23","B - 23","C - 23","D - 23"))
D014_siGlo <- D014 %>%
  filter(WELL.LABEL %in% c("A - 23","B - 23","C - 23","D - 23"))
D015_siGlo <- D015 %>%
  filter(WELL.LABEL %in% c("A - 23","B - 23","C - 23","D - 23"))
D016_siGlo <- D015 %>%
  filter(WELL.LABEL %in% c("A - 23","B - 23","C - 23","D - 23"))
G001_siGlo <- G001 %>%
  filter(WELL.LABEL %in% c("A - 23","B - 23","C - 23","D - 23"))
G002_siGlo <- G002 %>%
  filter(WELL.LABEL %in% c("A - 23","B - 23","C - 23","D - 23"))
G003_siGlo <- G003 %>%
  filter(WELL.LABEL %in% c("A - 23","B - 23","C - 23","D - 23"))
G004_siGlo <- G004 %>%
  filter(WELL.LABEL %in% c("A - 23","B - 23","C - 23","D - 23"))
G005_siGlo <- G005 %>%
  filter(WELL.LABEL %in% c("A - 23","B - 23","C - 23","D - 23"))
G006_siGlo <- G006 %>%
  filter(WELL.LABEL %in% c("A - 23","B - 23","C - 23","D - 23"))
G007_siGlo <- G007 %>%
  filter(WELL.LABEL %in% c("A - 23","B - 23","C - 23","D - 23"))
G008_siGlo <- G008 %>%
  filter(WELL.LABEL %in% c("A - 23","B - 23","C - 23","D - 23"))
G009_siGlo <- G009 %>%
  filter(WELL.LABEL %in% c("A - 23","B - 23","C - 23","D - 23"))
G010_siGlo <- G010 %>%
  filter(WELL.LABEL %in% c("A - 23","B - 23","C - 23","D - 23"))
G011_siGlo <- G011 %>%
  filter(WELL.LABEL %in% c("A - 23","B - 23","C - 23","D - 23"))
G012_siGlo <- G012 %>%
  filter(WELL.LABEL %in% c("A - 23","B - 23","C - 23","D - 23"))
G013_siGlo <- G013 %>%
  filter(WELL.LABEL %in% c("A - 23","B - 23","C - 23","D - 23"))
G014_siGlo <- G014 %>%
  filter(WELL.LABEL %in% c("A - 23","B - 23","C - 23","D - 23"))
G015_siGlo <- G015 %>%
  filter(WELL.LABEL %in% c("A - 23","B - 23","C - 23","D - 23"))
G016_siGlo <- G016 %>%
  filter(WELL.LABEL %in% c("A - 23","B - 23","C - 23","D - 23"))
G017_siGlo <- G017 %>%
  filter(WELL.LABEL %in% c("A - 23","B - 23","C - 23","D - 23"))
G018_siGlo <- G018 %>%
  filter(WELL.LABEL %in% c("A - 23","B - 23","C - 23","D - 23"))
G019_siGlo <- G019 %>%
  filter(WELL.LABEL %in% c("A - 23","B - 23","C - 23","D - 23"))
G020_siGlo <- G020 %>%
  filter(WELL.LABEL %in% c("A - 23","B - 23","C - 23","D - 23"))
G021_siGlo <- G021 %>%
  filter(WELL.LABEL %in% c("A - 23","B - 23","C - 23","D - 23"))
G022_siGlo <- G022 %>%
  filter(WELL.LABEL %in% c("A - 23","B - 23","C - 23","D - 23"))
G023_siGlo <- G023 %>%
  filter(WELL.LABEL %in% c("A - 23","B - 23","C - 23","D - 23"))
G024_siGlo <- G024 %>%
  filter(WELL.LABEL %in% c("A - 23","B - 23","C - 23","D - 23"))
G025_siGlo <- G025 %>%
  filter(WELL.LABEL %in% c("A - 23","B - 23","C - 23","D - 23"))
G026_siGlo <- G026 %>%
  filter(WELL.LABEL %in% c("A - 23","B - 23","C - 23","D - 23"))
G027_siGlo <- G027 %>%
  filter(WELL.LABEL %in% c("A - 23","B - 23","C - 23","D - 23"))
G028_siGlo <- G028 %>%
  filter(WELL.LABEL %in% c("A - 23","B - 23","C - 23","D - 23"))
G029_siGlo <- G029 %>%
  filter(WELL.LABEL %in% c("A - 23","B - 23","C - 23","D - 23"))
G030_siGlo <- G030 %>%
  filter(WELL.LABEL %in% c("A - 23","B - 23","C - 23","D - 23"))
G031_siGlo <- G031 %>%
  filter(WELL.LABEL %in% c("A - 23","B - 23","C - 23","D - 23"))
G032_siGlo <- G032 %>%
  filter(WELL.LABEL %in% c("A - 23","B - 23","C - 23","D - 23"))
G033_siGlo <- G033 %>%
  filter(WELL.LABEL %in% c("A - 23","B - 23","C - 23","D - 23"))
G034_siGlo <- G034 %>%
  filter(WELL.LABEL %in% c("A - 23","B - 23","C - 23","D - 23"))
G035_siGlo <- G035 %>%
  filter(WELL.LABEL %in% c("A - 23","B - 23","C - 23","D - 23"))
G036_siGlo <- G036 %>%
  filter(WELL.LABEL %in% c("A - 23","B - 23","C - 23","D - 23"))
G037_siGlo <- G037 %>%
  filter(WELL.LABEL %in% c("A - 23","B - 23","C - 23","D - 23"))
G038_siGlo <- G038 %>%
  filter(WELL.LABEL %in% c("A - 23","B - 23","C - 23","D - 23"))
G039_siGlo <- G039 %>%
  filter(WELL.LABEL %in% c("A - 23","B - 23","C - 23","D - 23"))
G040_siGlo <- G040 %>%
  filter(WELL.LABEL %in% c("A - 23","B - 23","C - 23","D - 23"))
G041_siGlo <- G041 %>%
  filter(WELL.LABEL %in% c("A - 23","B - 23","C - 23","D - 23"))
G042_siGlo <- G042 %>%
  filter(WELL.LABEL %in% c("A - 23","B - 23","C - 23","D - 23"))
G043_siGlo <- G043 %>%
  filter(WELL.LABEL %in% c("A - 23","B - 23","C - 23","D - 23"))
G044_siGlo <- G044 %>%
  filter(WELL.LABEL %in% c("A - 23","B - 23","C - 23","D - 23"))
G045_siGlo <- G045 %>%
  filter(WELL.LABEL %in% c("A - 23","B - 23","C - 23","D - 23"))
G046_siGlo <- G046 %>%
  filter(WELL.LABEL %in% c("A - 23","B - 23","C - 23","D - 23"))
G047_siGlo <- G047 %>%
  filter(WELL.LABEL %in% c("A - 23","B - 23","C - 23","D - 23"))
```

### 4. Create Batches

```
#Separate into screening batches

Batch_1_siGlo <- bind_rows(D001_siGlo,D002_siGlo,D003_siGlo,D004_siGlo,D005_siGlo,
                           D006_siGlo,D007_siGlo,D008_siGlo,D009_siGlo,D010_siGlo,
                           D011_siGlo,D012_siGlo,D013_siGlo,D014_siGlo,D015_siGlo,
                           D016_siGlo,G001_siGlo,G002_siGlo,G003_siGlo,G004_siGlo,
                           G005_siGlo)

Batch_2_siGlo <- bind_rows(G006_siGlo,G007_siGlo,G008_siGlo,G009_siGlo,G010_siGlo,
                           G011_siGlo,G012_siGlo,G013_siGlo,G014_siGlo,G015_siGlo,
                           G016_siGlo,G017_siGlo,G018_siGlo,G019_siGlo,G020_siGlo,
                           G021_siGlo,G022_siGlo,G023_siGlo,G024_siGlo,G025_siGlo,
                           G026_siGlo)

Batch_3_siGlo <- bind_rows(G027_siGlo,G028_siGlo,G029_siGlo,G030_siGlo,G031_siGlo,
                           G011_siGlo,G032_siGlo,G033_siGlo,G034_siGlo,G035_siGlo,
                           G036_siGlo,G037_siGlo,G038_siGlo,G039_siGlo,G040_siGlo,
                           G041_siGlo,G042_siGlo,G043_siGlo,G044_siGlo,G045_siGlo,
                           G046_siGlo)
```

### 5. Create Batch Means

```
#Batch1
Batch_1_Means <- colMeans(Batch_1_siGlo[,-1])
Batch_1_Means <- c("Mean", Batch_1_Means)
Batch_1_Means_DF <- rbind(Batch_1_siGlo,Batch_1_Means, stringsAsFactors = F)

Batch_1_SD <- Batch_1_siGlo %>%
  select_if(is.numeric) %>%
  summarise_all(sd, na.rm = TRUE) %>%
  unlist(use.names = FALSE) %>%
  c("SD", .)
Batch_1_siGlo_Complete <- rbind(Batch_1_Means_DF,Batch_1_SD, stringsAsFactors = F)


#Batch2
Batch_2_Means <- colMeans(Batch_2_siGlo[,-1])
Batch_2_Means <- c("Mean", Batch_2_Means)
Batch_2_Means_DF <- rbind(Batch_2_siGlo,Batch_2_Means, stringsAsFactors = F)

Batch_2_SD <- Batch_2_siGlo %>%
  select_if(is.numeric) %>%
  summarise_all(sd, na.rm = TRUE) %>%
  unlist(use.names = FALSE) %>%
  c("SD", .)
Batch_2_siGlo_Complete <- rbind(Batch_2_Means_DF,Batch_2_SD, stringsAsFactors = F)


#Batch3
Batch_3_Means <- colMeans(Batch_3_siGlo[,-1])
Batch_3_Means <- c("Mean", Batch_3_Means)
Batch_3_Means_DF <- rbind(Batch_3_siGlo,Batch_3_Means, stringsAsFactors = F)

Batch_3_SD <- Batch_3_siGlo %>%
  select_if(is.numeric) %>%
  summarise_all(sd, na.rm = TRUE) %>%
  unlist(use.names = FALSE) %>%
  c("SD", .)
Batch_3_siGlo_Complete <- rbind(Batch_3_Means_DF,Batch_3_SD, stringsAsFactors = F)
```

### 6. Write siGLO Batch Controls

```
write.csv(Batch_1_siGlo_Complete,"Batch_1_Complete.csv", row.names = FALSE)

write.csv(Batch_2_siGlo_Complete,"Batch_2_Complete.csv", row.names = FALSE)

write.csv(Batch_3_siGlo_Complete,"Batch_3_Complete.csv", row.names = FALSE)
```

### 7. Add Batch Means

```
#Add Batch 1 Mean/SD
D001 <- rbind(D001, Batch_1_Means,Batch_1_SD)
D002 <- rbind(D002, Batch_1_Means,Batch_1_SD) 
D003 <- rbind(D003, Batch_1_Means,Batch_1_SD) 
D004 <- rbind(D004, Batch_1_Means,Batch_1_SD) 
D005 <- rbind(D005, Batch_1_Means,Batch_1_SD) 
D006 <- rbind(D006, Batch_1_Means,Batch_1_SD) 
D007 <- rbind(D007, Batch_1_Means,Batch_1_SD)
D008 <- rbind(D008, Batch_1_Means,Batch_1_SD) 
D009 <- rbind(D009, Batch_1_Means,Batch_1_SD) 
D010 <- rbind(D010, Batch_1_Means,Batch_1_SD) 
D011 <- rbind(D011, Batch_1_Means,Batch_1_SD) 
D012 <- rbind(D012, Batch_1_Means,Batch_1_SD) 
D013 <- rbind(D013, Batch_1_Means,Batch_1_SD)
D014 <- rbind(D014, Batch_1_Means,Batch_1_SD) 
D015 <- rbind(D015, Batch_1_Means,Batch_1_SD) 
D016 <- rbind(D016, Batch_1_Means,Batch_1_SD) 
G001 <- rbind(G001, Batch_1_Means,Batch_1_SD)
G002 <- rbind(G002, Batch_1_Means,Batch_1_SD)
G003 <- rbind(G003, Batch_1_Means,Batch_1_SD)
G004 <- rbind(G004, Batch_1_Means,Batch_1_SD)
G005 <- rbind(G005, Batch_1_Means,Batch_1_SD)

#Add Batch 2 Mean/SD
G006 <- rbind(G006, Batch_2_Means,Batch_2_SD)
G007 <- rbind(G007, Batch_2_Means,Batch_2_SD)
G008 <- rbind(G008, Batch_2_Means,Batch_2_SD)
G009 <- rbind(G009, Batch_2_Means,Batch_2_SD)
G010 <- rbind(G010, Batch_2_Means,Batch_2_SD)
G011 <- rbind(G011, Batch_2_Means,Batch_2_SD)
G012 <- rbind(G012, Batch_2_Means,Batch_2_SD)
G013 <- rbind(G013, Batch_2_Means,Batch_2_SD)
G014 <- rbind(G014, Batch_2_Means,Batch_2_SD)
G015 <- rbind(G015, Batch_2_Means,Batch_2_SD)
G016 <- rbind(G016, Batch_2_Means,Batch_2_SD)
G017 <- rbind(G017, Batch_2_Means,Batch_2_SD)
G018 <- rbind(G018, Batch_2_Means,Batch_2_SD)
G019 <- rbind(G019, Batch_2_Means,Batch_2_SD)
G020 <- rbind(G020, Batch_2_Means,Batch_2_SD)
G021 <- rbind(G021, Batch_2_Means,Batch_2_SD)
G022 <- rbind(G022, Batch_2_Means,Batch_2_SD)
G023 <- rbind(G023, Batch_2_Means,Batch_2_SD)
G024 <- rbind(G024, Batch_2_Means,Batch_2_SD)
G025 <- rbind(G025, Batch_2_Means,Batch_2_SD)
G026 <- rbind(G026, Batch_2_Means,Batch_2_SD)


#Add Batch 1 Mean/SD
G027 <- rbind(G027, Batch_3_Means,Batch_3_SD)
G028 <- rbind(G028, Batch_3_Means,Batch_3_SD)
G029 <- rbind(G029, Batch_3_Means,Batch_3_SD)
G030 <- rbind(G030, Batch_3_Means,Batch_3_SD)
G031 <- rbind(G031, Batch_3_Means,Batch_3_SD)
G032 <- rbind(G032, Batch_3_Means,Batch_3_SD)
G033 <- rbind(G033, Batch_3_Means,Batch_3_SD)
G034 <- rbind(G034, Batch_3_Means,Batch_3_SD)
G035 <- rbind(G035, Batch_3_Means,Batch_3_SD)
G036 <- rbind(G036, Batch_3_Means,Batch_3_SD)
G037 <- rbind(G037, Batch_3_Means,Batch_3_SD)
G038 <- rbind(G038, Batch_3_Means,Batch_3_SD)
G039 <- rbind(G039, Batch_3_Means,Batch_3_SD)
G040 <- rbind(G040, Batch_3_Means,Batch_3_SD)
G041 <- rbind(G041, Batch_3_Means,Batch_3_SD)
G042 <- rbind(G042, Batch_3_Means,Batch_3_SD)
G043 <- rbind(G043, Batch_3_Means,Batch_3_SD)
G044 <- rbind(G044, Batch_3_Means,Batch_3_SD)
G045 <- rbind(G045, Batch_3_Means,Batch_3_SD)
G046 <- rbind(G046, Batch_3_Means,Batch_3_SD)
G047 <- rbind(G047, Batch_3_Means,Batch_3_SD)
```

### 8. Create Z-Score Function

```
# Z Score = (value - batch DMSO mean) / batch DMSO sd

transform_dataframe <- function(input_df) {
  # Remove the first column and convert the remaining columns to numeric
  D001_New <- input_df[,-1]
  D001_New <- D001_New %>% mutate_all(as.numeric)
  
  # Create a new dataframe for the transformed values
  D001_Normalized <- D001_New  # Create a copy of the original dataframe
  
  # Divide each column by its corresponding value in row 385
  for (i in 1:ncol(D001_Normalized)) {
    D001_Normalized[, i] <- D001_Normalized[, i] - D001_New[385, i]
  }
  
  for (i in 1:ncol(D001_Normalized)) {
    D001_Normalized[, i] <- D001_Normalized[, i] / D001_New[386, i]
  }
  
  # Add the first column back to the transformed dataframe with the name "WELL.LABEL"
  D001_Normalized <- cbind("Plate Name" = deparse(substitute(input_df)), WELL.LABEL = input_df[, 1], D001_Normalized)
  
  return(D001_Normalized)
}

# Example usage:
# D001_PhenoMeasures <- transform_dataframe(D001)
```

### 9. Z-Score Transformations

```
#Transform data using Z_Function
D001_PhenoMeasures <- transform_dataframe(D001)
D002_PhenoMeasures <- transform_dataframe(D002)
D003_PhenoMeasures <- transform_dataframe(D003)
D004_PhenoMeasures <- transform_dataframe(D004)
D005_PhenoMeasures <- transform_dataframe(D005)
D006_PhenoMeasures <- transform_dataframe(D006)
D007_PhenoMeasures <- transform_dataframe(D007)
D008_PhenoMeasures <- transform_dataframe(D008)
D009_PhenoMeasures <- transform_dataframe(D009)
D010_PhenoMeasures <- transform_dataframe(D010)
D011_PhenoMeasures <- transform_dataframe(D011)
D012_PhenoMeasures <- transform_dataframe(D012)
D013_PhenoMeasures <- transform_dataframe(D013)
D014_PhenoMeasures <- transform_dataframe(D014)
D015_PhenoMeasures <- transform_dataframe(D015)
D016_PhenoMeasures <- transform_dataframe(D016)

G001_PhenoMeasures <- transform_dataframe(G001)
G002_PhenoMeasures <- transform_dataframe(G002)
G003_PhenoMeasures <- transform_dataframe(G003)
G004_PhenoMeasures <- transform_dataframe(G004)
G005_PhenoMeasures <- transform_dataframe(G005)
G006_PhenoMeasures <- transform_dataframe(G006)
G007_PhenoMeasures <- transform_dataframe(G007)
G008_PhenoMeasures <- transform_dataframe(G008)
G009_PhenoMeasures <- transform_dataframe(G009)
G010_PhenoMeasures <- transform_dataframe(G010)
G011_PhenoMeasures <- transform_dataframe(G011)
G012_PhenoMeasures <- transform_dataframe(G012)
G013_PhenoMeasures <- transform_dataframe(G013)
G014_PhenoMeasures <- transform_dataframe(G014)
G015_PhenoMeasures <- transform_dataframe(G015)
G016_PhenoMeasures <- transform_dataframe(G016)
G017_PhenoMeasures <- transform_dataframe(G017)
G018_PhenoMeasures <- transform_dataframe(G018)
G019_PhenoMeasures <- transform_dataframe(G019)
G020_PhenoMeasures <- transform_dataframe(G020)
G021_PhenoMeasures <- transform_dataframe(G021)
G022_PhenoMeasures <- transform_dataframe(G022)
G023_PhenoMeasures <- transform_dataframe(G023)
G024_PhenoMeasures <- transform_dataframe(G024)
G025_PhenoMeasures <- transform_dataframe(G025)
G026_PhenoMeasures <- transform_dataframe(G026)
G027_PhenoMeasures <- transform_dataframe(G027)
G028_PhenoMeasures <- transform_dataframe(G028)
G029_PhenoMeasures <- transform_dataframe(G029)
G030_PhenoMeasures <- transform_dataframe(G030)
G031_PhenoMeasures <- transform_dataframe(G031)
G032_PhenoMeasures <- transform_dataframe(G032)
G033_PhenoMeasures <- transform_dataframe(G033)
G034_PhenoMeasures <- transform_dataframe(G034)
G035_PhenoMeasures <- transform_dataframe(G035)
G036_PhenoMeasures <- transform_dataframe(G036)
G037_PhenoMeasures <- transform_dataframe(G037)
G038_PhenoMeasures <- transform_dataframe(G038)
G039_PhenoMeasures <- transform_dataframe(G039)
G040_PhenoMeasures <- transform_dataframe(G040)
G041_PhenoMeasures <- transform_dataframe(G041)
G042_PhenoMeasures <- transform_dataframe(G042)
G043_PhenoMeasures <- transform_dataframe(G043)
G044_PhenoMeasures <- transform_dataframe(G044)
G045_PhenoMeasures <- transform_dataframe(G045)
G046_PhenoMeasures <- transform_dataframe(G046)
G047_PhenoMeasures <- transform_dataframe(G047)
```

### 10. Export Pheno Measures

```
# Save D001_PhenoMeasures to CSV
write.csv(D001_PhenoMeasures, "D001_PhenoMeasures.csv", row.names = FALSE)

# Save D002_PhenoMeasures to CSV
write.csv(D002_PhenoMeasures, "D002_PhenoMeasures.csv", row.names = FALSE)

# Save D003_PhenoMeasures to CSV
write.csv(D003_PhenoMeasures, "D003_PhenoMeasures.csv", row.names = FALSE)

# Save D004_PhenoMeasures to CSV
write.csv(D004_PhenoMeasures, "D004_PhenoMeasures.csv", row.names = FALSE)

# Save D005_PhenoMeasures to CSV
write.csv(D005_PhenoMeasures, "D005_PhenoMeasures.csv", row.names = FALSE)

# Save D006_PhenoMeasures to CSV
write.csv(D006_PhenoMeasures, "D006_PhenoMeasures.csv", row.names = FALSE)

# Save D007_PhenoMeasures to CSV
write.csv(D007_PhenoMeasures, "D007_PhenoMeasures.csv", row.names = FALSE)

# Save D008_PhenoMeasures to CSV
write.csv(D008_PhenoMeasures, "D008_PhenoMeasures.csv", row.names = FALSE)

# Save D009_PhenoMeasures to CSV
write.csv(D009_PhenoMeasures, "D009_PhenoMeasures.csv", row.names = FALSE)

# Save D010_PhenoMeasures to CSV
write.csv(D010_PhenoMeasures, "D010_PhenoMeasures.csv", row.names = FALSE)

# Save D011_PhenoMeasures to CSV
write.csv(D011_PhenoMeasures, "D011_PhenoMeasures.csv", row.names = FALSE)

# Save D012_PhenoMeasures to CSV
write.csv(D012_PhenoMeasures, "D012_PhenoMeasures.csv", row.names = FALSE)

# Save D013_PhenoMeasures to CSV
write.csv(D013_PhenoMeasures, "D013_PhenoMeasures.csv", row.names = FALSE)

# Save D014_PhenoMeasures to CSV
write.csv(D014_PhenoMeasures, "D014_PhenoMeasures.csv", row.names = FALSE)

# Save D015_PhenoMeasures to CSV
write.csv(D015_PhenoMeasures, "D015_PhenoMeasures.csv", row.names = FALSE)

# Save D016_PhenoMeasures to CSV
write.csv(D016_PhenoMeasures, "D016_PhenoMeasures.csv", row.names = FALSE)

# Continue this pattern for the G001 to G047 dataframes.

# Save G001_PhenoMeasures to CSV
write.csv(G001_PhenoMeasures, "G001_PhenoMeasures.csv", row.names = FALSE)

# Save G002_PhenoMeasures to CSV
write.csv(G002_PhenoMeasures, "G002_PhenoMeasures.csv", row.names = FALSE)

# Save G003_PhenoMeasures to CSV
write.csv(G003_PhenoMeasures, "G003_PhenoMeasures.csv", row.names = FALSE)

# Save G004_PhenoMeasures to CSV
write.csv(G004_PhenoMeasures, "G004_PhenoMeasures.csv", row.names = FALSE)

# Save G005_PhenoMeasures to CSV
write.csv(G005_PhenoMeasures, "G005_PhenoMeasures.csv", row.names = FALSE)

# Save G006_PhenoMeasures to CSV
write.csv(G006_PhenoMeasures, "G006_PhenoMeasures.csv", row.names = FALSE)

# Save G007_PhenoMeasures to CSV
write.csv(G007_PhenoMeasures, "G007_PhenoMeasures.csv", row.names = FALSE)

# Save G008_PhenoMeasures to CSV
write.csv(G008_PhenoMeasures, "G008_PhenoMeasures.csv", row.names = FALSE)

# Save G009_PhenoMeasures to CSV
write.csv(G009_PhenoMeasures, "G009_PhenoMeasures.csv", row.names = FALSE)

# Save G010_PhenoMeasures to CSV
write.csv(G010_PhenoMeasures, "G010_PhenoMeasures.csv", row.names = FALSE)

# Save G011_PhenoMeasures to CSV
write.csv(G011_PhenoMeasures, "G011_PhenoMeasures.csv", row.names = FALSE)

# Save G012_PhenoMeasures to CSV
write.csv(G012_PhenoMeasures, "G012_PhenoMeasures.csv", row.names = FALSE)

# Save G013_PhenoMeasures to CSV
write.csv(G013_PhenoMeasures, "G013_PhenoMeasures.csv", row.names = FALSE)

# Save G014_PhenoMeasures to CSV
write.csv(G014_PhenoMeasures, "G014_PhenoMeasures.csv", row.names = FALSE)

# Save G015_PhenoMeasures to CSV
write.csv(G015_PhenoMeasures, "G015_PhenoMeasures.csv", row.names = FALSE)

# Save G016_PhenoMeasures to CSV
write.csv(G016_PhenoMeasures, "G016_PhenoMeasures.csv", row.names = FALSE)

# Save G017_PhenoMeasures to CSV
write.csv(G017_PhenoMeasures, "G017_PhenoMeasures.csv", row.names = FALSE)

# Save G018_PhenoMeasures to CSV
write.csv(G018_PhenoMeasures, "G018_PhenoMeasures.csv", row.names = FALSE)

# Save G019_PhenoMeasures to CSV
write.csv(G019_PhenoMeasures, "G019_PhenoMeasures.csv", row.names = FALSE)

# Save G020_PhenoMeasures to CSV
write.csv(G020_PhenoMeasures, "G020_PhenoMeasures.csv", row.names = FALSE)

# Save G021_PhenoMeasures to CSV
write.csv(G021_PhenoMeasures, "G021_PhenoMeasures.csv", row.names = FALSE)

# Save G022_PhenoMeasures to CSV
write.csv(G022_PhenoMeasures, "G022_PhenoMeasures.csv", row.names = FALSE)

# Save G023_PhenoMeasures to CSV
write.csv(G023_PhenoMeasures, "G023_PhenoMeasures.csv", row.names = FALSE)

# Save G024_PhenoMeasures to CSV
write.csv(G024_PhenoMeasures, "G024_PhenoMeasures.csv", row.names = FALSE)

# Save G025_PhenoMeasures to CSV
write.csv(G025_PhenoMeasures, "G025_PhenoMeasures.csv", row.names = FALSE)

# Save G026_PhenoMeasures to CSV
write.csv(G026_PhenoMeasures, "G026_PhenoMeasures.csv", row.names = FALSE)

# Save G027_PhenoMeasures to CSV
write.csv(G027_PhenoMeasures, "G027_PhenoMeasures.csv", row.names = FALSE)

# Save G028_PhenoMeasures to CSV
write.csv(G028_PhenoMeasures, "G028_PhenoMeasures.csv", row.names = FALSE)

# Save G029_PhenoMeasures to CSV
write.csv(G029_PhenoMeasures, "G029_PhenoMeasures.csv", row.names = FALSE)

# Save G030_PhenoMeasures to CSV
write.csv(G030_PhenoMeasures, "G030_PhenoMeasures.csv", row.names = FALSE)

# Save G031_PhenoMeasures to CSV
write.csv(G031_PhenoMeasures, "G031_PhenoMeasures.csv", row.names = FALSE)

# Save G032_PhenoMeasures to CSV
write.csv(G032_PhenoMeasures, "G032_PhenoMeasures.csv", row.names = FALSE)

# Save G033_PhenoMeasures to CSV
write.csv(G033_PhenoMeasures, "G033_PhenoMeasures.csv", row.names = FALSE)

# Save G034_PhenoMeasures to CSV
write.csv(G034_PhenoMeasures, "G034_PhenoMeasures.csv", row.names = FALSE)

# Save G035_PhenoMeasures to CSV
write.csv(G035_PhenoMeasures, "G035_PhenoMeasures.csv", row.names = FALSE)

# Save G036_PhenoMeasures to CSV
write.csv(G036_PhenoMeasures, "G036_PhenoMeasures.csv", row.names = FALSE)

# Save G037_PhenoMeasures to CSV
write.csv(G037_PhenoMeasures, "G037_PhenoMeasures.csv", row.names = FALSE)

# Save G038_PhenoMeasures to CSV
write.csv(G038_PhenoMeasures, "G038_PhenoMeasures.csv", row.names = FALSE)

# Save G039_PhenoMeasures to CSV
write.csv(G039_PhenoMeasures, "G039_PhenoMeasures.csv", row.names = FALSE)

# Save G040_PhenoMeasures to CSV
write.csv(G040_PhenoMeasures, "G040_PhenoMeasures.csv", row.names = FALSE)

# Save G041_PhenoMeasures to CSV
write.csv(G041_PhenoMeasures, "G041_PhenoMeasures.csv", row.names = FALSE)

# Save G042_PhenoMeasures to CSV
write.csv(G042_PhenoMeasures, "G042_PhenoMeasures.csv", row.names = FALSE)

# Save G043_PhenoMeasures to CSV
write.csv(G043_PhenoMeasures, "G043_PhenoMeasures.csv", row.names = FALSE)

# Save G044_PhenoMeasures to CSV
write.csv(G044_PhenoMeasures, "G044_PhenoMeasures.csv", row.names = FALSE)

# Save G045_PhenoMeasures to CSV
write.csv(G045_PhenoMeasures, "G045_PhenoMeasures.csv", row.names = FALSE)

# Save G046_PhenoMeasures to CSV
write.csv(G046_PhenoMeasures, "G046_PhenoMeasures.csv", row.names = FALSE)

# Save G047_PhenoMeasures to CSV
write.csv(G047_PhenoMeasures, "G047_PhenoMeasures.csv", row.names = FALSE)
```

### 11. Data Export

```
# List of PhenoMeasures dataframe names
df_names <- c("D001_PhenoMeasures", "D002_PhenoMeasures", "D003_PhenoMeasures", "D004_PhenoMeasures", "D005_PhenoMeasures",
              "D006_PhenoMeasures", "D007_PhenoMeasures", "D008_PhenoMeasures", "D009_PhenoMeasures", "D010_PhenoMeasures",
              "D011_PhenoMeasures", "D012_PhenoMeasures", "D013_PhenoMeasures", "D014_PhenoMeasures", "D015_PhenoMeasures",
              "D016_PhenoMeasures", "G001_PhenoMeasures", "G002_PhenoMeasures", "G003_PhenoMeasures", "G004_PhenoMeasures",
              "G005_PhenoMeasures", "G006_PhenoMeasures", "G007_PhenoMeasures", "G008_PhenoMeasures", "G009_PhenoMeasures",
              "G010_PhenoMeasures", "G011_PhenoMeasures", "G012_PhenoMeasures", "G013_PhenoMeasures", "G014_PhenoMeasures",
              "G015_PhenoMeasures", "G016_PhenoMeasures", "G017_PhenoMeasures", "G018_PhenoMeasures", "G019_PhenoMeasures",
              "G020_PhenoMeasures", "G021_PhenoMeasures", "G022_PhenoMeasures", "G023_PhenoMeasures", "G024_PhenoMeasures",
              "G025_PhenoMeasures", "G026_PhenoMeasures", "G027_PhenoMeasures", "G028_PhenoMeasures", "G029_PhenoMeasures",
              "G030_PhenoMeasures", "G031_PhenoMeasures", "G032_PhenoMeasures", "G033_PhenoMeasures", "G034_PhenoMeasures",
              "G035_PhenoMeasures", "G036_PhenoMeasures", "G037_PhenoMeasures", "G038_PhenoMeasures", "G039_PhenoMeasures",
              "G040_PhenoMeasures", "G041_PhenoMeasures", "G042_PhenoMeasures", "G043_PhenoMeasures", "G044_PhenoMeasures",
              "G045_PhenoMeasures", "G046_PhenoMeasures", "G047_PhenoMeasures")

# Initialize an empty list to store the dataframes
combined_dataframes <- list()

# Loop through the dataframe names
for (df_name in df_names) {
  # Get the dataframe
  df <- get(df_name)
  # Exclude the last two rows
  df <- df[1:(nrow(df) - 2), ]
  # Add the dataframe to the list
  combined_dataframes[[df_name]] <- df
}

# Combine the dataframes into one large dataframe
All_Z_Scores <- do.call(rbind, combined_dataframes)

siRNAs <- read.csv("siRNA Locations.csv",fileEncoding = 'UTF-8-BOM')

All_Z_Scores <- cbind(siRNAs,All_Z_Scores)

All_Z_Scores <- All_Z_Scores %>% na.omit()

# Save All_Z_Scores to CSV
write.csv(All_Z_Scores, "All_Z_Scores.csv", row.names = FALSE)
```
