## Supplementary material for "SAMP-Score: A morphology-based machine learning classification method for screening pro-senescence compounds in p16 positive cancer cells": HTML Walkthrough Guides for Code: Supplemental Data 1_2_HeLa_Screen_SAMP.html

2\_HeLa-Screen-SAMP.knit


### 1\_HeLa Heatmap.R

```
library("gplots")
```

```
## 
## Attaching package: 'gplots'
```

```
## The following object is masked from 'package:stats':
## 
##     lowess
```

```
library("RColorBrewer")
library("matrixStats")
library("plyr")
```

```
## 
## Attaching package: 'plyr'
```

```
## The following object is masked from 'package:matrixStats':
## 
##     count
```

```
library("dplyr")
```

```
## 
## Attaching package: 'dplyr'
```

```
## The following objects are masked from 'package:plyr':
## 
##     arrange, count, desc, failwith, id, mutate, rename, summarise,
##     summarize
```

```
## The following object is masked from 'package:matrixStats':
## 
##     count
```

```
## The following objects are masked from 'package:stats':
## 
##     filter, lag
```

```
## The following objects are masked from 'package:base':
## 
##     intersect, setdiff, setequal, union
```

```
library("data.table")
```

```
## 
## Attaching package: 'data.table'
```

```
## The following objects are masked from 'package:dplyr':
## 
##     between, first, last
```

```
library("stringr")
library("ggplot2")


# Upload HeLa Screen file 
all <- read.csv("All_Z_Scores.csv")

all_small <- all %>%
  filter(!grepl("Blank", Gene.Symbol))

### Sanity check - NA removed in previous stage 
all_removed <- all %>%
  filter(grepl("Blank", Gene.Symbol))

# Update DF ###
all <- all_small

#Deal with duplicate Gene Entries (12 in total)#
all <- all %>%
  group_by(RefSeq.Accession.Number) %>%
  mutate(RefSeq.Accession.Number = ifelse(row_number() == 1, RefSeq.Accession.Number, paste0(RefSeq.Accession.Number, "_A")))

row_names <- all$RefSeq.Accession.Number

# Remove unnecessary measures (i.e. extra label columns)
all <- all[,6:42]

#Convert to data matrix 
all_matrix <- ((as.matrix(all)))
rownames(all_matrix)<- row_names 

# Create heat map
breaks <- unique(c(seq(-5,-1,length=100),seq(-1,1,length=100), seq(1,5,length=100)))
my_palette <- colorRampPalette(c("yellow","black","black","purple"))(length(breaks)-1)
heatmap.2(t(all_matrix),
          Rowv = T,
          Colv = T,
          col=my_palette,
          breaks=breaks,
          density.info="none",
          trace="none",
          #main = "199 compounds; Length or Prev",
          dendrogram=c("both"), 
          symm=F,symkey=F,symbreaks=T,
          labRow=F,
          labCol=F,
          cexRow=0.8,
          cexCol=0.1,
          margins = c(8,2),
          key.title = "1" , key.xlab="Z Score", 
          #rowsep =c(0,4,12,19,31,37,43),
          #colsep =c(0,233),
          sepcolor = c("black"),sepwidth = c(0.05, 0.05),
          #ColSideColors=condition_colors, scale="none",
          distfun = function(x) dist(x, method = "euclidean"),
          hclust=function(x) hclust(x,method="ward.D2"))
```

```
# Look at Nuclei Count vs Cell Area - Might Remove
plot <- ggplot(data = all, aes(x = Nuclei.Nuclei.Count.wv1, y = Cells.Area.wv3)) + 
  geom_point() + 
  geom_smooth(method = "lm") +  theme_bw() +
  scale_fill_manual(values = c("#999999", "#E69F00")) +
  labs(title = "Nuclei Count vs Cell Area", x = "Nuclei Count (Z-Score)", y = "Cell Area (Z-Score)")

# Fit linear regression model
lm_model <- lm(Cells.Area.wv3 ~ Nuclei.Nuclei.Count.wv1, data = all)

# Extract R-squared value
r_squared <- summary(lm_model)$r.squared

# Add text annotation for R-squared value
plot <- plot + 
  annotate("text", x = max(all$Nuclei.Nuclei.Count.wv1), y = max(all$Cells.Area.wv3),
           label = paste("R^2 =", round(r_squared, 3)), parse = TRUE, hjust = 2, vjust = 10) 

#plot

#Save plot 
ggsave(plot = plot,"Full_Count_vs_Area_Corr.png")
```

```
## Saving 7 x 5 in image
```

```
## `geom_smooth()` using formula = 'y ~ x'
```

### 2\_Hits Nuclei Count and Area Only.R

```
library("gplots") 
library("RColorBrewer")
library("matrixStats")
library("plyr")
library("dplyr")
library("data.table")
library("stringr")
library("ggplot2")
library("Rtsne")

# Upload HeLa Screen file 
all <- read.csv("All_Z_Scores.csv")

# Define Hit Parameters - This can be Adjusted # 
all <- all %>%
  filter(`Nuclei.Nuclei.Count.wv1`  < -3)

all_small <- all %>%
  filter(!grepl("Blank", Gene.Symbol))

### Sanity check - NA removed in previous stage
all_removed <- all %>%
  filter(grepl("Blank", Gene.Symbol))

# Update DF ###
all <- all_small

#Deal with duplicate Gene Entries (12 in total)#
all <- all %>%
  group_by(RefSeq.Accession.Number) %>%
  mutate(RefSeq.Accession.Number = ifelse(row_number() == 1, RefSeq.Accession.Number, paste0(RefSeq.Accession.Number, "_A")))

row_names <- all$RefSeq.Accession.Number

# Remove unnecessary measures (i.e. extra label columns)
all <- all[,c(6,27)]


#Convert to data matrix 
all_matrix <- ((as.matrix(all)))
rownames(all_matrix)<- row_names 

# Create heat map
breaks <- unique(c(seq(-5,-1,length=100),seq(-1,0.1,length=100), seq(1,5,length=100)))
my_palette <- colorRampPalette(c("yellow","black","black","purple"))(length(breaks)-1)
heatmap.2(t(all_matrix),
          Rowv = T,
          Colv = T,
          col=my_palette,
          breaks=breaks,
          density.info="none",
          trace="none",
          #main = "199 compounds; Length or Prev",
          dendrogram=c("both"), 
          symm=F,symkey=F,symbreaks=T,
          labRow=F,
          labCol=F,
          cexRow=0.8,
          cexCol=0.1,
          margins = c(8,2),
          key.title = "1" , key.xlab="Z Score", 
          #rowsep =c(0,4,12,19,31,37,43),
          #colsep =c(0,233),
          sepcolor = c("black"),sepwidth = c(0.05, 0.05),
          #ColSideColors=condition_colors, scale="none",
          distfun = function(x) dist(x, method = "euclidean"),
          hclust=function(x) hclust(x,method="ward.D2"))
```

```
Hits <- rownames(all_matrix)
write.csv(Hits, "Hits_Nuclei_Count_Only.csv")

# Look at Nuclei Count vs Cell Area
plot <- ggplot(data = all, aes(x = Nuclei.Nuclei.Count.wv1, y = Cells.Area.wv3)) + 
  geom_point() + 
  geom_smooth(method = "lm") +  theme_bw() +
  scale_fill_manual(values = c("#999999", "#E69F00")) +
  labs(title = "Nuclei Count vs Cell Area", x = "Nuclei Count (Z-Score)", y = "Cell Area (Z-Score)")

# Fit linear regression model
lm_model <- lm(Cells.Area.wv3 ~ Nuclei.Nuclei.Count.wv1, data = all)

# Extract R-squared value
r_squared <- summary(lm_model)$r.squared

# Add text annotation for R-squared value
plot <- plot + 
  annotate("text", x = max(all$Nuclei.Nuclei.Count.wv1), y = max(all$Cells.Area.wv3),
           label = paste("R^2 =", round(r_squared, 3)), parse = TRUE, hjust = 2, vjust = 10) 

plot
```

```
## `geom_smooth()` using formula = 'y ~ x'
```

```
#Save plot 
ggsave(plot = plot,"Count_Threshold_Count_vs_Area_Corr.png")
```

```
## Saving 7 x 5 in image
## `geom_smooth()` using formula = 'y ~ x'
```

### 3\_Filtered Heatmap Count and Area.R

```
library("gplots") 
library("RColorBrewer")
library("matrixStats")
library("plyr")
library("dplyr")
library("data.table")
library("stringr")
library("ggplot2")
library("Rtsne")

# Upload your file 
all <- read.csv("All_Z_Scores.csv")

# Define Hit Parameters - This can be Adjusted # 
all <- all %>%
  filter(`Nuclei.Nuclei.Count.wv1` < -3, Cells.Area.wv3 >3)


all_small <- all %>%
  filter(!grepl("Blank", Gene.Symbol))

### Sanity check - NA removed in previous stage 
all_removed <- all %>%
  filter(grepl("Blank", Gene.Symbol))

# Update DF ###
all <- all_small

#Deal with duplicate Gene Entries (12 in total)#
all <- all %>%
  group_by(RefSeq.Accession.Number) %>%
  mutate(RefSeq.Accession.Number = ifelse(row_number() == 1, RefSeq.Accession.Number, paste0(RefSeq.Accession.Number, "_A")))

row_names <- all$RefSeq.Accession.Number

# Remove unnecessary measures (i.e. extra label columns)
all <- all[,6:42]


#Convert to data matrix 
all_matrix <- ((as.matrix(all)))
rownames(all_matrix)<- row_names 

# Create heat map
breaks <- unique(c(seq(-5,-1,length=100),seq(-1,0.1,length=100), seq(1,5,length=100)))
my_palette <- colorRampPalette(c("yellow","black","black","purple"))(length(breaks)-1)
heatmap.2(t(all_matrix),
          Rowv = T,
          Colv = T,
          col=my_palette,
          breaks=breaks,
          density.info="none",
          trace="none",
          #main = "199 compounds; Length or Prev",
          dendrogram=c("both"), 
          symm=F,symkey=F,symbreaks=T,
          labRow=F,
          labCol=F,
          cexRow=0.8,
          cexCol=0.1,
          margins = c(8,2),
          key.title = "1" , key.xlab="Z Score", 
          #rowsep =c(0,4,12,19,31,37,43),
          #colsep =c(0,233),
          sepcolor = c("black"),sepwidth = c(0.05, 0.05),
          #ColSideColors=condition_colors, scale="none",
          distfun = function(x) dist(x, method = "euclidean"),
          hclust=function(x) hclust(x,method="ward.D2"))
```

```
Hits <- rownames(all_matrix)
write.csv(Hits, "Hits.csv")
```

### 4\_Hits 1.96.R

```
library("gplots") 
library("RColorBrewer")
library("matrixStats")
library("plyr")
library("dplyr")
library("data.table")
library("stringr")
library("ggplot2")
library("Rtsne")

# Upload your file 
all <- read.csv("All_Z_Scores.csv")

# Define Hit Parameters - This can be Adjusted # 
all <- all %>%
  filter(`Nuclei.Nuclei.Count.wv1` < -1.96, Cells.Area.wv3 >1.96)

all_small <- all %>%
  filter(!grepl("Blank", Gene.Symbol))

### Sanity check - NA removed in previous stage 
all_removed <- all %>%
  filter(grepl("Blank", Gene.Symbol))

# Update DF ###
all <- all_small

#Deal with duplicate Gene Entries (12 in total)#
all <- all %>%
  group_by(RefSeq.Accession.Number) %>%
  mutate(RefSeq.Accession.Number = ifelse(row_number() == 1, RefSeq.Accession.Number, paste0(RefSeq.Accession.Number, "_A")))

row_names <- all$RefSeq.Accession.Number

# Remove unnecessary measures (i.e. extra label columns)
all <- all[,6:42]

#Convert to data matrix 
all_matrix <- ((as.matrix(all)))
rownames(all_matrix)<- row_names 

# Create heat map
breaks <- unique(c(seq(-5,-1,length=100),seq(-1,0.1,length=100), seq(1,5,length=100)))
my_palette <- colorRampPalette(c("yellow","black","black","purple"))(length(breaks)-1)
heatmap.2(t(all_matrix),
          Rowv = T,
          Colv = T,
          col=my_palette,
          breaks=breaks,
          density.info="none",
          trace="none",
          #main = "199 compounds; Length or Prev",
          dendrogram=c("both"), 
          symm=F,symkey=F,symbreaks=T,
          labRow=F,
          labCol=F,
          cexRow=0.8,
          cexCol=0.1,
          margins = c(8,2),
          key.title = "1" , key.xlab="Z Score", 
          #rowsep =c(0,4,12,19,31,37,43),
          #colsep =c(0,233),
          sepcolor = c("black"),sepwidth = c(0.05, 0.05),
          #ColSideColors=condition_colors, scale="none",
          distfun = function(x) dist(x, method = "euclidean"),
          hclust=function(x) hclust(x,method="ward.D2"))
```

```
Hits <- rownames(all_matrix)
write.csv(Hits, "Hits_1.96.csv")
```

### 5\_Hits 1.R

```
library("gplots") 
library("RColorBrewer")
library("matrixStats")
library("plyr")
library("dplyr")
library("data.table")
library("stringr")
library("ggplot2")
library("Rtsne")

# Upload your file 
all <- read.csv("All_Z_Scores.csv")

# Define Hit Parameters - This can be Adjusted # 
all <- all %>%
  filter(`Nuclei.Nuclei.Count.wv1` < -1, Cells.Area.wv3 >1)


all_small <- all %>%
  filter(!grepl("Blank", Gene.Symbol))

### Sanity check - NA removed in previous stage 
all_removed <- all %>%
  filter(grepl("Blank", Gene.Symbol))

# Update DF ###
all <- all_small

#Deal with duplicate Gene Entries (12 in total)#
all <- all %>%
  group_by(RefSeq.Accession.Number) %>%
  mutate(RefSeq.Accession.Number = ifelse(row_number() == 1, RefSeq.Accession.Number, paste0(RefSeq.Accession.Number, "_A")))

row_names <- all$RefSeq.Accession.Number

# Remove unnecessary measures (i.e. extra label columns)
all <- all[,6:42]


#Convert to data matrix 
all_matrix <- ((as.matrix(all)))
rownames(all_matrix)<- row_names 

# Create heat map
breaks <- unique(c(seq(-5,-1,length=100),seq(-1,0.1,length=100), seq(1,5,length=100)))
my_palette <- colorRampPalette(c("yellow","black","black","purple"))(length(breaks)-1)
heatmap.2(t(all_matrix),
          Rowv = T,
          Colv = T,
          col=my_palette,
          breaks=breaks,
          density.info="none",
          trace="none",
          #main = "199 compounds; Length or Prev",
          dendrogram=c("both"), 
          symm=F,symkey=F,symbreaks=T,
          labRow=F,
          labCol=F,
          cexRow=0.8,
          cexCol=0.1,
          margins = c(8,2),
          key.title = "1" , key.xlab="Z Score", 
          #rowsep =c(0,4,12,19,31,37,43),
          #colsep =c(0,233),
          sepcolor = c("black"),sepwidth = c(0.05, 0.05),
          #ColSideColors=condition_colors, scale="none",
          distfun = function(x) dist(x, method = "euclidean"),
          hclust=function(x) hclust(x,method="ward.D2"))
```

```
Hits <- rownames(all_matrix)
write.csv(Hits, "Hits_1.csv")
```

### 6\_SENCAN Down.R

```
library("gplots") 
library("RColorBrewer")
library("matrixStats")
library("plyr")
library("dplyr")
library("data.table")
library("stringr")
library("ggplot2")
library("Rtsne")

# Upload your file 
all <- read.csv("All_Z_Scores.csv")
Filter <- read.csv("SENCAN Down.csv")

Filter<-Filter$converted_alias

# Define Hit Parameters - This can be Adjusted # 
all <- all %>%
  filter(RefSeq.Accession.Number %in% Filter)

all_small <- all %>%
  filter(!grepl("Blank", Gene.Symbol))

### Sanity check - NA removed in previous stage
all_removed <- all %>%
  filter(grepl("Blank", Gene.Symbol))

# Update DF ###
all <- all_small

#Deal with duplicate Gene Entries (12 in total)#
all <- all %>%
  group_by(RefSeq.Accession.Number) %>%
  mutate(RefSeq.Accession.Number = ifelse(row_number() == 1, RefSeq.Accession.Number, paste0(RefSeq.Accession.Number, "_A")))

row_names <- all$RefSeq.Accession.Number

# Remove unnecessary measures (i.e. extra label columns)
all <- all[,6:42]


#Convert to data matrix 
all_matrix <- ((as.matrix(all)))
rownames(all_matrix)<- row_names 

# Create heat map
breaks <- unique(c(seq(-5,-1,length=100),seq(-1,0.1,length=100), seq(1,5,length=100)))
my_palette <- colorRampPalette(c("yellow","black","black","purple"))(length(breaks)-1)
heatmap.2(t(all_matrix),
          Rowv = T,
          Colv = T,
          col=my_palette,
          breaks=breaks,
          density.info="none",
          trace="none",
          #main = "199 compounds; Length or Prev",
          dendrogram=c("both"), 
          symm=F,symkey=F,symbreaks=T,
          labRow=F,
          labCol=F,
          cexRow=0.8,
          cexCol=0.1,
          margins = c(8,2),
          key.title = "1" , key.xlab="Z Score", 
          #rowsep =c(0,4,12,19,31,37,43),
          #colsep =c(0,233),
          sepcolor = c("black"),sepwidth = c(0.05, 0.05),
          #ColSideColors=condition_colors, scale="none",
          distfun = function(x) dist(x, method = "euclidean"),
          hclust=function(x) hclust(x,method="ward.D2"))
```

```
Hits <- rownames(all_matrix)
write.csv(Hits, "SENCAN_Down.csv")
```

### 7\_Senescopedia.R

```
library("gplots") 
library("RColorBrewer")
library("matrixStats")
library("plyr")
library("dplyr")
library("data.table")
library("stringr")
library("ggplot2")
library("Rtsne")

# Upload your file 
all <- read.csv("All_Z_Scores.csv")
Filter <- read.csv("Senescopeida List.csv")

Filter<-Filter$converted_alias

# Define Hit Parameters - This can be Adjusted # 
all <- all %>%
  filter(RefSeq.Accession.Number %in% Filter)


all_small <- all %>%
  filter(!grepl("Blank", Gene.Symbol))

### Sanity check - NA removed in previous stage 
all_removed <- all %>%
  filter(grepl("Blank", Gene.Symbol))

# Update DF ###
all <- all_small

#Deal with duplicate Gene Entries (12 in total)#
all <- all %>%
  group_by(RefSeq.Accession.Number) %>%
  mutate(RefSeq.Accession.Number = ifelse(row_number() == 1, RefSeq.Accession.Number, paste0(RefSeq.Accession.Number, "_A")))

row_names <- all$RefSeq.Accession.Number

# Remove unnecessary measures (i.e. extra label columns)
all <- all[,6:42]


#Convert to data matrix 
all_matrix <- ((as.matrix(all)))
rownames(all_matrix)<- row_names 

# Create heat map
breaks <- unique(c(seq(-5,-1,length=100),seq(-1,0.1,length=100), seq(1,5,length=100)))
my_palette <- colorRampPalette(c("yellow","black","black","purple"))(length(breaks)-1)
heatmap.2(t(all_matrix),
          Rowv = T,
          Colv = T,
          col=my_palette,
          breaks=breaks,
          density.info="none",
          trace="none",
          #main = "199 compounds; Length or Prev",
          dendrogram=c("both"), 
          symm=F,symkey=F,symbreaks=T,
          labRow=F,
          labCol=F,
          cexRow=0.8,
          cexCol=0.1,
          margins = c(8,2),
          key.title = "1" , key.xlab="Z Score", 
          #rowsep =c(0,4,12,19,31,37,43),
          #colsep =c(0,233),
          sepcolor = c("black"),sepwidth = c(0.05, 0.05),
          #ColSideColors=condition_colors, scale="none",
          distfun = function(x) dist(x, method = "euclidean"),
          hclust=function(x) hclust(x,method="ward.D2"))
```

```
Hits <- rownames(all_matrix)
write.csv(Hits, "Senescopeida.csv")
```

### 8\_Cell Cycle.R

```
library("gplots") 
library("RColorBrewer")
library("matrixStats")
library("plyr")
library("dplyr")
library("data.table")
library("stringr")
library("ggplot2")
library("Rtsne")

# Upload your file 
all <- read.csv("All_Z_Scores.csv")
Filter <- read.csv("Cell Cycle Genes.csv")

Filter<-Filter$converted_alias

# Define Hit Parameters - This can be Adjusted # 
all <- all %>%
  filter(RefSeq.Accession.Number %in% Filter)


all_small <- all %>%
  filter(!grepl("Blank", Gene.Symbol))

### Sanity check - NA removed in previous stage 
all_removed <- all %>%
  filter(grepl("Blank", Gene.Symbol))

# Update DF ###
all <- all_small

#Deal with duplicate Gene Entries (12 in total)#
all <- all %>%
  group_by(RefSeq.Accession.Number) %>%
  mutate(RefSeq.Accession.Number = ifelse(row_number() == 1, RefSeq.Accession.Number, paste0(RefSeq.Accession.Number, "_A")))

row_names <- all$RefSeq.Accession.Number

# Remove unnecessary measures (i.e. extra label columns)
all <- all[,6:42]


#Convert to data matrix 
all_matrix <- ((as.matrix(all)))
rownames(all_matrix)<- row_names 

# Create heat map
breaks <- unique(c(seq(-5,-1,length=100),seq(-1,0.1,length=100), seq(1,5,length=100)))
my_palette <- colorRampPalette(c("yellow","black","black","purple"))(length(breaks)-1)
heatmap.2(t(all_matrix),
          Rowv = T,
          Colv = T,
          col=my_palette,
          breaks=breaks,
          density.info="none",
          trace="none",
          #main = "199 compounds; Length or Prev",
          dendrogram=c("both"), 
          symm=F,symkey=F,symbreaks=T,
          labRow=F,
          labCol=F,
          cexRow=0.8,
          cexCol=0.1,
          margins = c(8,2),
          key.title = "1" , key.xlab="Z Score", 
          #rowsep =c(0,4,12,19,31,37,43),
          #colsep =c(0,233),
          sepcolor = c("black"),sepwidth = c(0.05, 0.05),
          #ColSideColors=condition_colors, scale="none",
          distfun = function(x) dist(x, method = "euclidean"),
          hclust=function(x) hclust(x,method="ward.D2"))
```

```
Hits <- rownames(all_matrix)
write.csv(Hits, "Cell Cycle.csv")
```

### 9\_Senescence.R

```
library("gplots") 
library("RColorBrewer")
library("matrixStats")
library("plyr")
library("dplyr")
library("data.table")
library("stringr")
library("ggplot2")
library("Rtsne")

# Upload your file 
all <- read.csv("All_Z_Scores.csv")
Filter <- read.csv("Senescence Kegg.csv")

Filter<-Filter$converted_alias

# Define Hit Parameters - This can be Adjusted # 
all <- all %>%
  filter(RefSeq.Accession.Number %in% Filter)


all_small <- all %>%
  filter(!grepl("Blank", Gene.Symbol))

### Sanity check - NA removed in previous stage 
all_removed <- all %>%
  filter(grepl("Blank", Gene.Symbol))

# Update DF ###
all <- all_small

#Deal with duplicate Gene Entries (12 in total)#
all <- all %>%
  group_by(RefSeq.Accession.Number) %>%
  mutate(RefSeq.Accession.Number = ifelse(row_number() == 1, RefSeq.Accession.Number, paste0(RefSeq.Accession.Number, "_A")))

row_names <- all$RefSeq.Accession.Number

# Remove unnecessary measures (i.e. extra label columns)
all <- all[,6:42]


#Convert to data matrix 
all_matrix <- ((as.matrix(all)))
rownames(all_matrix)<- row_names 

# Create heat map
breaks <- unique(c(seq(-5,-1,length=100),seq(-1,0.1,length=100), seq(1,5,length=100)))
my_palette <- colorRampPalette(c("yellow","black","black","purple"))(length(breaks)-1)
heatmap.2(t(all_matrix),
          Rowv = T,
          Colv = T,
          col=my_palette,
          breaks=breaks,
          density.info="none",
          trace="none",
          #main = "199 compounds; Length or Prev",
          dendrogram=c("both"), 
          symm=F,symkey=F,symbreaks=T,
          labRow=F,
          labCol=F,
          cexRow=0.8,
          cexCol=0.1,
          margins = c(8,2),
          key.title = "1" , key.xlab="Z Score", 
          #rowsep =c(0,4,12,19,31,37,43),
          #colsep =c(0,233),
          sepcolor = c("black"),sepwidth = c(0.05, 0.05),
          #ColSideColors=condition_colors, scale="none",
          distfun = function(x) dist(x, method = "euclidean"),
          hclust=function(x) hclust(x,method="ward.D2"))
```

```
Hits <- rownames(all_matrix)
write.csv(Hits, "Senescence.csv")
```

### 10\_Extracting Clusters.R

```
# Load necessary libraries
library(gplots)
library(RColorBrewer)
library(matrixStats)
library(plyr)
library(dplyr)
library(data.table)
library(stringr)
library(ggplot2)

# Upload your file 
all <- read.csv("All_Z_Scores.csv")

all_small <- all %>%
  filter(!grepl("Blank", Gene.Symbol))

# Sanity check - NA removed in previous stage 
all_removed <- all %>%
  filter(grepl("Blank", Gene.Symbol))

# Update DF
all <- all_small

# Deal with duplicate Gene Entries (12 in total)
all <- all %>%
  group_by(RefSeq.Accession.Number) %>%
  mutate(RefSeq.Accession.Number = ifelse(row_number() == 1, RefSeq.Accession.Number, paste0(RefSeq.Accession.Number, "_A")))

row_names <- all$RefSeq.Accession.Number

# Remove unnecessary measures (i.e. extra label columns)
all <- all[, 6:42]

# Convert to data matrix 
all_matrix <- as.matrix(all)
rownames(all_matrix) <- row_names 

# Clustering for dendrogram
row_clust <- hclust(dist(all_matrix, method = 'euclidean'), method = 'ward.D2')

# Plotting the dendrogram
par(cex=0.1, mar=c(5, 8, 4, 1))
plot(row_clust, xlab="", ylab="", main="", sub="", axes=FALSE)
par(cex=1)
title(main="Dendrogram")
axis(2)
```

```
# Define number of clusters
k <- 3
clust <- sort(cutree(row_clust, k=k))
clust2 <- data.matrix(clust)

# Save cluster locations in an accessible csv file 
write.csv(clust2, "Clusters.csv", row.names = TRUE)

clust2 <- as.data.frame(clust2)

# Iterate through each cluster and generate heatmaps
for (i in 1:k) {
  # Filter cluster
  current_cluster <- clust2 %>% filter(clust == i)
  
  # Get the names of the current cluster
  current_cluster_names <- row.names(current_cluster)
  
  # Subset matrix for the current cluster
  current_cluster_HM <- subset(all_matrix, rownames(all_matrix) %in% rownames(current_cluster))
  
  # Transpose the matrix before saving to CSV
  write.csv(t(current_cluster_HM), paste0("Cluster_", i, ".csv"))
  
  # Define color palette and breaks
  breaks <- unique(c(seq(-5, -1, length=100), seq(-1, 0.1, length=100), seq(1, 5, length=100)))
  my_palette <- colorRampPalette(c("yellow", "black", "black", "purple"))(length(breaks) - 1)
  
  # Create heatmap
  heatmap.2(t(current_cluster_HM),
            Rowv = TRUE,
            Colv = TRUE,
            col = my_palette,
            breaks = breaks,
            density.info = "none",
            trace = "none",
            dendrogram = c("both"), 
            symm = FALSE,
            symkey = FALSE,
            symbreaks = TRUE,
            labRow = FALSE,
            labCol = FALSE,
            cexRow = 0.8,
            cexCol = 0.1,
            margins = c(8, 2),
            key.title = "1",
            key.xlab = "Z Score",
            sepcolor = c("black"),
            sepwidth = c(0.05, 0.05),
            distfun = function(x) dist(x, method = "euclidean"),
            hclust = function(x) hclust(x, method = "ward.D2"))
}
```

### 11\_K-Means Clustering

```
library("gplots") 
library("RColorBrewer")
library("matrixStats")
library("plyr")
library("dplyr")
library("data.table")
library("stringr")
library("ggplot2")
library(umap)

# Upload your file 
all <- read.csv("All_Z_Scores.csv")

all_small <- all %>%
  filter(!grepl("Blank", Gene.Symbol))

### Sanity check 
all_removed <- all %>%
  filter(grepl("Blank", Gene.Symbol))

# Update DF ###
all <- all_small

#Deal with duplicate Gene Entries (12 in total)#
all <- all %>%
  group_by(RefSeq.Accession.Number) %>%
  mutate(RefSeq.Accession.Number = ifelse(row_number() == 1, RefSeq.Accession.Number, paste0(RefSeq.Accession.Number, "_A")))

row_names <- all$RefSeq.Accession.Number

# Remove unnecessary measures (i.e. extra label columns)
all <- all[,6:42]


# Set the seed for reproducibility
set.seed(123)  # You can change the seed for different results

# Perform K-means clustering with K = 4
kmeans_result <- kmeans(all, centers = 4, nstart = 25)

# Add the cluster assignment to the original dataset
all$Cluster <- as.factor(kmeans_result$cluster)

# Run UMAP to reduce dimensionality
umap_result <- umap(all[, -ncol(all)])  # Exclude the 'Cluster' column for UMAP

# Prepare data for plotting
umap_data <- as.data.frame(umap_result$layout)
umap_data$Cluster <- all$Cluster

# Calculate the count of samples per cluster
cluster_counts <- all %>%
  count(Cluster) %>%
  mutate(Cluster = paste0(Cluster, " (n=", n, ")"))

# Update the UMAP data with the new cluster labels
umap_data$Cluster <- factor(umap_data$Cluster, levels = levels(all$Cluster), labels = cluster_counts$Cluster)


# Plot UMAP with clusters, larger text, and theme_bw
umap_plot <- ggplot(umap_data, aes(x = V1, y = V2, color = Cluster)) +
  geom_point(size = 0.2) +
  labs(title = "UMAP Plot with K-means Clusters (K=3)",
       x = "UMAP 1", y = "UMAP 2") +
  theme_bw() +  # Change to theme_bw
  theme(
    legend.position = "right",  # Adjust the legend position if needed
    text = element_text(size = 14)  # Make all text larger
  ) +
  scale_color_manual(
    values = c("#E41A1C","#377EB8", "pink", "#4DAF4A"),  # Ensure these colors correspond to your clusters
    breaks = levels(umap_data$Cluster)  # Ensure that breaks match the levels of your Cluster variable
  )

# Save the plot as a PNG file in the "k_means_umap" folder
ggsave(filename = "k_means_umap/UMAP_Kmeans_Clusters.png", plot = umap_plot, width = 8, height = 6)
```

### 12\_K-Means Heatmaps

```
library(gplots)
library(RColorBrewer)
library(matrixStats)
library(plyr)
library(dplyr)
library(data.table)
library(stringr)
library(ggplot2)

# Upload your file 
all <- read.csv("All_Z_Scores.csv")

# Filter out rows with "Blank" in Gene.Symbol
all_small <- all %>%
  filter(!grepl("Blank", Gene.Symbol))

### Sanity check
all_removed <- all %>%
  filter(grepl("Blank", Gene.Symbol))

# Update the main dataframe
all <- all_small

# Handle duplicate Gene Entries (12 in total)
all <- all %>%
  group_by(RefSeq.Accession.Number) %>%
  mutate(RefSeq.Accession.Number = ifelse(row_number() == 1, RefSeq.Accession.Number, paste0(RefSeq.Accession.Number, "_A")))

# Save RefSeq.Accession.Number for later use
refseq_numbers <- all$RefSeq.Accession.Number

# Remove unnecessary columns (keep only relevant measures for clustering)
all_for_clustering <- all[, 6:42]  # Assuming columns 6 to 42 are for clustering

# Set the seed for reproducibility
set.seed(123)

# Perform K-means clustering with K = 3
k <- 4  # Define the number of clusters
kmeans_result <- kmeans(all_for_clustering, centers = k, nstart = 25)

# Add the cluster assignment to the original dataset
all$Cluster <- as.factor(kmeans_result$cluster)

# Create a matrix for heatmap generation
all_matrix <- as.matrix(all_for_clustering)  # Convert to matrix for clustering
rownames(all_matrix) <- refseq_numbers  # Set row names for the matrix

# Create a directory for saving heatmaps
dir.create("kmeans_heatmaps", showWarnings = FALSE)

# Iterate through each cluster and generate heatmaps and CSVs
for (i in 1:k) {  # Use the number of clusters directly
  # Filter the current cluster
  current_cluster <- all %>% filter(Cluster == i)
  
  # Get the names of the current cluster
  current_cluster_names <- current_cluster$RefSeq.Accession.Number
  
  # Subset matrix for the current cluster using row names
  current_cluster_HM <- all_matrix[current_cluster_names, , drop = FALSE]  # Ensure drop = FALSE to maintain matrix dimensions
  
  # Define filename for the heatmap
  png_filename <- paste0("kmeans_heatmaps/KMeans_Cluster_", i, ".png")
  
  # Save the heatmap as a PNG file
  png(png_filename, width = 800, height = 800)
  
  # Define color palette and breaks
  breaks <- unique(c(seq(-5, -1, length=100), seq(-1, 0.1, length=100), seq(1, 5, length=100)))
  my_palette <- colorRampPalette(c("yellow", "black", "black", "purple"))(length(breaks) - 1)
  
  # Create heatmap
  heatmap.2(t(current_cluster_HM),
            Rowv = TRUE,
            Colv = TRUE,
            col = my_palette,
            breaks = breaks,
            density.info = "none",
            trace = "none",
            dendrogram = c("both"), 
            symm = FALSE,
            symkey = FALSE,
            symbreaks = TRUE,
            labRow = FALSE,
            labCol = FALSE,
            cexRow = 0.8,
            cexCol = 0.1,
            margins = c(8, 2),
            key.title = "1",
            key.xlab = "Z Score",
            sepcolor = c("black"),
            sepwidth = c(0.05, 0.05),
            distfun = function(x) dist(x, method = "euclidean"),
            hclust = function(x) hclust(x, method = "ward.D2"))
  
  dev.off()  # Close the PNG device
  
  # Save current cluster data to a CSV file in the working directory
  csv_filename <- paste0("KMeans_Cluster_", i, ".csv")  # No directory specified, save to the working directory
  write.csv(current_cluster, file = csv_filename, row.names = FALSE)  # Save the cluster data
}
```

### 13\_Extract Common Hits

```
library(VennDiagram)
```

```
## Loading required package: grid
```

```
## Loading required package: futile.logger
```

```
# Read in the data
KMeans_Cluster_1 <- read.csv("KMeans_Cluster_1.csv")
KMeans_Cluster_4 <- read.csv("KMeans_Cluster_4.csv")
Heir_Cluster_1 <- read.csv("Cluster_1.csv")

# Extract the identifiers (RefSeq.Accession.Number)
KMeans_Cluster_1_names <- KMeans_Cluster_1$RefSeq.Accession.Number
KMeans_Cluster_4_names <- KMeans_Cluster_4$RefSeq.Accession.Number
Heir_Cluster_1_names <- colnames(Heir_Cluster_1)

# Remove the first column from hierarchical cluster if needed
Heir_Cluster_1_names <- Heir_Cluster_1_names[-1]

# Convert to sets for Venn diagram
KMeans_Cluster_1_set <- unique(KMeans_Cluster_1_names)
KMeans_Cluster_4_set <- unique(KMeans_Cluster_4_names)
Heir_Cluster_1_set <- unique(Heir_Cluster_1_names)

# Print the sizes of the sets to ensure they are populated
print(paste("KMeans Cluster 1 size:", length(KMeans_Cluster_1_set)))
```

```
## [1] "KMeans Cluster 1 size: 6527"
```

```
print(paste("KMeans Cluster 4 size:", length(KMeans_Cluster_4_set)))
```

```
## [1] "KMeans Cluster 4 size: 749"
```

```
print(paste("Hierarchical Cluster 1 size:", length(Heir_Cluster_1_set)))
```

```
## [1] "Hierarchical Cluster 1 size: 5329"
```

```
# Find common elements between KMeans Cluster 1 and Hierarchical Cluster 1
common_KMeans1_Heir1 <- intersect(KMeans_Cluster_1_set, Heir_Cluster_1_set)

# Print the number of common elements to check the intersection
print(paste("Common elements between KMeans Cluster 1 and Hierarchical Cluster 1:", length(common_KMeans1_Heir1)))
```

```
## [1] "Common elements between KMeans Cluster 1 and Hierarchical Cluster 1: 4240"
```

```
# Combine the common elements from KMeans 1 and Hierarchical 1 with all elements from KMeans Cluster 4
final_hits <- union(KMeans_Cluster_4_set, common_KMeans1_Heir1)

# Check the number of elements in the final set
print(paste("Final hits size:", length(final_hits)))
```

```
## [1] "Final hits size: 4989"
```

```
# If final_hits is non-empty, create the data frame and save it
if (length(final_hits) > 0) {
  final_hits_df <- data.frame(Final_Hits = final_hits)
  
  # Save the final list to a CSV file in the working directory
  common_file <- "common_senescence_hits.csv"
  write.csv(final_hits_df, file = common_file, row.names = FALSE)
  print(paste("CSV file saved with", nrow(final_hits_df), "rows"))
} else {
  print("No final hits found.")
}
```

```
## [1] "CSV file saved with 4989 rows"
```

### 14\_Use Common Hits to Label Senescence Ground Truth

```
library(gplots)
library(RColorBrewer)
library(dplyr)

# Read your original file and the common hits file
all <- read.csv("All_Z_Scores.csv")
common_hits <- read.csv("common_senescence_hits.csv")

# Extract common hits
common_hits_set <- common_hits$Final_Hits  # Update to match the column name from the new common_hits file

# Filter out rows with "Blank" in Gene.Symbol
all_small <- all %>%
  filter(!grepl("Blank", Gene.Symbol))

# Handle duplicate Gene Entries
all <- all_small %>%
  group_by(RefSeq.Accession.Number) %>%
  mutate(RefSeq.Accession.Number = ifelse(row_number() == 1, RefSeq.Accession.Number, paste0(RefSeq.Accession.Number, "_A")))

# Create a new dataframe with "Sen" and "NonSen" labels based on common hits
all_labeled <- all %>%
  mutate(Senescence_Status = ifelse(RefSeq.Accession.Number %in% common_hits_set, "Sen", "NonSen"))

# Save RefSeq.Accession.Number for later use
refseq_numbers <- all_labeled$RefSeq.Accession.Number

# Remove unnecessary columns (keep only relevant measures for heatmap)
all_for_heatmap <- all_labeled[, 6:42]  # Assuming columns 6 to 42 are for the heatmap

# Create a matrix for heatmap generation
all_matrix <- as.matrix(all_for_heatmap)  # Convert to matrix for heatmap
rownames(all_matrix) <- refseq_numbers  # Set row names for the matrix

# Create a directory for saving heatmaps
dir.create("heatmaps", showWarnings = FALSE)

# Generate heatmaps for "Sen" and "NonSen" categories
for (label in c("Sen", "NonSen")) {
  # Filter the current label
  current_label <- all_labeled %>% filter(Senescence_Status == label)
  
  # Get the names of the current label
  current_label_names <- current_label$RefSeq.Accession.Number
  
  # Subset matrix for the current label using row names
  current_label_HM <- all_matrix[current_label_names, , drop = FALSE]  # Ensure drop = FALSE to maintain matrix dimensions
  
  # Define filename for the heatmap
  png_filename <- paste0("heatmaps/", label, "_Heatmap.png")
  
  # Save the heatmap as a PNG file
  png(png_filename, width = 800, height = 800)
  
  # Define color palette and breaks
  breaks <- unique(c(seq(-5, -1, length = 100), seq(-1, 0.1, length = 100), seq(1, 5, length = 100)))
  my_palette <- colorRampPalette(c("yellow", "black", "black", "purple"))(length(breaks) - 1)
  
  # Create heatmap
  heatmap.2(t(current_label_HM),
            Rowv = TRUE,
            Colv = TRUE,
            col = my_palette,
            breaks = breaks,
            density.info = "none",
            trace = "none",
            dendrogram = c("both"), 
            symm = FALSE,
            symkey = FALSE,
            symbreaks = TRUE,
            labRow = FALSE,
            labCol = FALSE,
            cexRow = 0.8,
            cexCol = 0.1,
            margins = c(8, 2),
            key.title = "1",
            key.xlab = "Z Score",
            sepcolor = c("black"),
            sepwidth = c(0.05, 0.05),
            distfun = function(x) dist(x, method = "euclidean"),
            hclust = function(x) hclust(x, method = "ward.D2"))
  
  dev.off()  # Close the PNG device
}

# Save the updated dataframe with "Senescence_Status" labels
write.csv(all_labeled, "All_Data_Clusters_Consensus.csv", row.names = TRUE)
```

### 15\_Make full Venn Diagram

```
library(VennDiagram)

# Read in the data
KMeans_Cluster_1 <- read.csv("KMeans_Cluster_1.csv")
KMeans_Cluster_4 <- read.csv("KMeans_Cluster_4.csv")
Heir_Cluster_1 <- read.csv("Cluster_1.csv")
Heir_Cluster_3 <- read.csv("Cluster_3.csv")  # Add Hierarchical Cluster 3

# Extract the identifiers (RefSeq.Accession.Number)
KMeans_Cluster_1_names <- KMeans_Cluster_1$RefSeq.Accession.Number
KMeans_Cluster_4_names <- KMeans_Cluster_4$RefSeq.Accession.Number
Heir_Cluster_1_names <- colnames(Heir_Cluster_1)
Heir_Cluster_3_names <- colnames(Heir_Cluster_3)  # Add identifiers for Hierarchical Cluster 3

# Remove the first column from hierarchical clusters if needed
Heir_Cluster_1_names <- Heir_Cluster_1_names[-1]
Heir_Cluster_3_names <- Heir_Cluster_3_names[-1]  # Adjust if needed

# Convert to sets for Venn diagram
KMeans_Cluster_1_set <- unique(KMeans_Cluster_1_names)
KMeans_Cluster_4_set <- unique(KMeans_Cluster_4_names)
Heir_Cluster_1_set <- unique(Heir_Cluster_1_names)
Heir_Cluster_3_set <- unique(Heir_Cluster_3_names)  # Add set for Hierarchical Cluster 3

# Create a Venn diagram with four sets
venn.plot <- venn.diagram(
  x = list(
    "KMeans Cluster 1" = KMeans_Cluster_1_set,
    "KMeans Cluster 4" = KMeans_Cluster_4_set,
    "Hierarchical Cluster 1" = Heir_Cluster_1_set,
    "Hierarchical Cluster 3" = Heir_Cluster_3_set
  ),
  category.names = c("KMeans Cluster 1", "KMeans Cluster 4", "Hierarchical Cluster 1", "Hierarchical Cluster 3"),
  filename = NULL,  # Output to R plot device
  output = TRUE
)

# Plot the Venn diagram
png(filename = "venn_diagram_4_clusters.png", width = 800, height = 800)
grid.draw(venn.plot)
dev.off()
```

```
## quartz_off_screen 
##                 2
```

### 16\_Matching Thresholds Percentage.R

```
library("gplots") 
library("RColorBrewer")
library("matrixStats")
library("plyr")
library("dplyr")
library("data.table")
library("stringr")
library("ggplot2")
library("Rtsne")

# List of file paths
file_paths <- c(
  "Hits_Nuc_Count_Only.csv",
  "Hits.csv",
  "Hits_1.96.csv",
  "Hits_1.csv",
  "SENCAN.csv",
  "SENCAN_Down.csv",
  "Senescopeida.csv",
  "Senescopeida_Sig.csv",
  "Cell Cycle.csv",
  "Senescence.csv"
)

# Create a function to load a file and extract the second column as a vector
load_and_extract_second_column <- function(file_path) {
  # Load the file into a data frame
  data <- read.csv(file_path)
  
  # Extract the second column as a vector
  vector <- data[[2]]
  
  return(vector)
}

# Use lapply to load the files and create a list of vectors
vector_list <- lapply(file_paths, load_and_extract_second_column)

# Override the original data frames with the vectors
Count_Only <- vector_list[[1]]
Hits_High <- vector_list[[2]]
Hits_Med <- vector_list[[3]]
Hits_Low <- vector_list[[4]]
SENCAN <- vector_list[[5]]
SENCAN_Neg <- vector_list[[6]]
Senescopedia_Down <- vector_list[[7]]
Senescopedia_Sig <- vector_list[[8]]
Cell_Cycle <- vector_list[[9]]
Senescence <- vector_list[[10]]

# Now for Clusters
Sen_Hits <- common_hits$Final_Hits

library(dplyr)

# Create a dataframe with the counts of matched entries
result <- data.frame(
  Vector = c("SENCAN", "SENCAN_Neg", "Senescopedia_Down", "Senescopedia_Sig", "Cell_Cycle", "Senescence", "Sen_Hits"),
  Count_Only = sapply(list(SENCAN, SENCAN_Neg, Senescopedia_Down, Senescopedia_Sig, Cell_Cycle, Senescence, Sen_Hits), function(vector) round((sum(vector %in% Count_Only) / length(vector) * 100), 1)),
  Hits_High = sapply(list(SENCAN, SENCAN_Neg, Senescopedia_Down, Senescopedia_Sig, Cell_Cycle, Senescence, Sen_Hits), function(vector) round((sum(vector %in% Hits_High) / length(vector) * 100), 1)),
  Hits_Med = sapply(list(SENCAN, SENCAN_Neg, Senescopedia_Down, Senescopedia_Sig, Cell_Cycle, Senescence, Sen_Hits), function(vector) round((sum(vector %in% Hits_Med) / length(vector) * 100), 1)),
  Hits_Low = sapply(list(SENCAN, SENCAN_Neg, Senescopedia_Down, Senescopedia_Sig, Cell_Cycle, Senescence, Sen_Hits), function(vector) round((sum(vector %in% Hits_Low) / length(vector) * 100), 1))
)

# Print the result dataframe
Test <- result

print(Test)
```

```
##              Vector Count_Only Hits_High Hits_Med Hits_Low
## 1            SENCAN        5.0       1.0      3.0     13.0
## 2        SENCAN_Neg        5.6       1.9      5.6     14.8
## 3 Senescopedia_Down        6.9       1.9      4.4     20.9
## 4  Senescopedia_Sig        7.7       2.2      5.5     16.5
## 5        Cell_Cycle        5.8       0.0      7.5     20.0
## 6        Senescence        4.1       0.0      5.4     21.8
## 7          Sen_Hits       19.6       2.0      9.3     43.1
```

### 17\_Export Threshold Percentages.R

```
library("gplots") 
library("RColorBrewer")
library("matrixStats")
library("plyr")
library("dplyr")
library("data.table")
library("stringr")
library("ggplot2")
library("Rtsne")

# List of file paths
file_paths <- c(
  "Hits_Nuc_Count_Only.csv",
  "Hits.csv",
  "Hits_1.96.csv",
  "Hits_1.csv",
  "SENCAN.csv",
  "SENCAN_Down.csv",
  "Senescopeida.csv",
  "Senescopeida_Sig.csv",
  "Cell Cycle.csv",
  "Senescence.csv"
)

# Create a function to load a file and extract the second column as a vector
load_and_extract_second_column <- function(file_path) {
  # Load the file into a data frame
  data <- read.csv(file_path)
  
  # Extract the second column as a vector
  vector <- data[[2]]
  
  return(vector)
}

# Use lapply to load the files and create a list of vectors
vector_list <- lapply(file_paths, load_and_extract_second_column)

# Override the original data frames with the vectors
Count_Only <- vector_list[[1]]
Hits_High <- vector_list[[2]]
Hits_Med <- vector_list[[3]]
Hits_Low <- vector_list[[4]]
SENCAN <- vector_list[[5]]
SENCAN_Neg <- vector_list[[6]]
Senescopedia_Down <- vector_list[[7]]
Senescopedia_Sig <- vector_list[[8]]
Cell_Cycle <- vector_list[[9]]
Senescence <- vector_list[[10]]

# Now for Clusters
Sen_Hits <- common_hits$Final_Hits


library(dplyr)


result <- data.frame(
  Vector = c("SENCAN", "SENCAN_Neg", "Senescopedia_Down", "Senescopedia_Sig", "Cell_Cycle", "Senescence", "Sen_Hits"),
  Count_Only = sapply(list(SENCAN, SENCAN_Neg, Senescopedia_Down, Senescopedia_Sig, Cell_Cycle, Senescence, Sen_Hits), function(vector) round((sum(vector %in% Count_Only) / length(Count_Only) * 100), 1)),
  Hits_High = sapply(list(SENCAN, SENCAN_Neg, Senescopedia_Down, Senescopedia_Sig, Cell_Cycle, Senescence, Sen_Hits), function(vector) round((sum(vector %in% Hits_High) / length(Hits_High) * 100), 1)),
  Hits_Med = sapply(list(SENCAN, SENCAN_Neg, Senescopedia_Down, Senescopedia_Sig, Cell_Cycle, Senescence, Sen_Hits), function(vector) round((sum(vector %in% Hits_Med) / length(Hits_Med) * 100), 1)),
  Hits_Low = sapply(list(SENCAN, SENCAN_Neg, Senescopedia_Down, Senescopedia_Sig, Cell_Cycle, Senescence, Sen_Hits), function(vector) round((sum(vector %in% Hits_Low) / length(Hits_Low) * 100), 1))
)

# Print the result dataframe
Test <- result

library(ggplot2)

# Reshape the Test dataframe for ggplot2
Test <- Test %>% filter (Vector %in% c("Sen_Hits"))
Test_melted <- melt(Test, id.vars = "Vector")
```

```
## Warning in melt(Test, id.vars = "Vector"): The melt generic in data.table has
## been passed a data.frame and will attempt to redirect to the relevant reshape2
## method; please note that reshape2 is deprecated, and this redirection is now
## deprecated as well. To continue using melt methods from reshape2 while both
## libraries are attached, e.g. melt.list, you can prepend the namespace like
## reshape2::melt(Test). In the next version, this warning will become an error.
```

```
Test_melted <- Test_melted %>% filter(grepl("Hits", variable))

# Define a custom color palette
my_colors <- c("#E41A1C", "#377EB8", "#4DAF4A")

# Create a prettier square faceted bar chart with a fixed y-axis of 100 and custom colors
ggplot(Test_melted, aes(x = Vector, y = value, fill = Vector)) +
  geom_bar(stat = "identity") +
  facet_wrap(~ variable, ncol = 2, scales = "free") +
  labs(title = "Percentage Comparisons",
       x = "Vector",
       y = "Percentage") +
  theme_minimal() +
  theme(axis.text.x = element_text(angle = 45, hjust = 1),
        legend.position = "right",
        text = element_text(face = "bold")) +
  scale_fill_manual(values = my_colors) +
  ylim(0, 100) +
  guides(fill = guide_legend(title = "Vector")) +
  scale_y_continuous(labels = scales::percent_format(scale = 1), limits = c(0, 100))
```

```
## Scale for y is already present.
## Adding another scale for y, which will replace the existing scale.
```

```
print(Test_melted)
```

```
##     Vector  variable value
## 1 Sen_Hits Hits_High  74.8
## 2 Sen_Hits  Hits_Med  88.5
## 3 Sen_Hits  Hits_Low  78.6
```

### 18\_Look\_at\_GT\_Count\_and\_Area

```
library(gplots)
library(RColorBrewer)
library(dplyr)

# Read your original file and the common hits file
all <- read.csv("All_Z_Scores.csv")
common_hits <- read.csv("common_senescence_hits.csv")

all <- all[,c(1,2,6,27)]

# Extract common hits
common_hits_set <- common_hits$Final_Hits  # Update to match the column name from the new common_hits file

# Filter out rows with "Blank" in Gene.Symbol
all_small <- all %>%
  filter(!grepl("Blank", Gene.Symbol))

# Handle duplicate Gene Entries
all <- all_small %>%
  group_by(RefSeq.Accession.Number) %>%
  mutate(RefSeq.Accession.Number = ifelse(row_number() == 1, RefSeq.Accession.Number, paste0(RefSeq.Accession.Number, "_A")))

# Create a new dataframe with "Sen" and "NonSen" labels based on common hits
all_labeled <- all %>%
  mutate(Senescence_Status = ifelse(RefSeq.Accession.Number %in% common_hits_set, "Sen", "NonSen"))

# Save RefSeq.Accession.Number for later use
refseq_numbers <- all_labeled$RefSeq.Accession.Number

# Remove unnecessary columns (keep only relevant measures for heatmap)
all_for_heatmap <- all_labeled[, 3:4]  # Assuming columns 6 to 42 are for the heatmap

# Create a matrix for heatmap generation
all_matrix <- as.matrix(all_for_heatmap)  # Convert to matrix for heatmap
rownames(all_matrix) <- refseq_numbers  # Set row names for the matrix

# Create a directory for saving heatmaps
dir.create("heatmaps", showWarnings = FALSE)

# Generate heatmaps for "Sen" and "NonSen" categories
for (label in c("Sen", "NonSen")) {
  # Filter the current label
  current_label <- all_labeled %>% filter(Senescence_Status == label)
  
  # Get the names of the current label
  current_label_names <- current_label$RefSeq.Accession.Number
  
  # Subset matrix for the current label using row names
  current_label_HM <- all_matrix[current_label_names, , drop = FALSE]  # Ensure drop = FALSE to maintain matrix dimensions
  
  # Define filename for the heatmap
  png_filename <- paste0("heatmaps/", label, "_Heatmap_Count_Area.png")
  
  # Save the heatmap as a PNG file
  png(png_filename, width = 800, height = 800)
  
  # Define color palette and breaks
  breaks <- unique(c(seq(-5, -1, length = 100), seq(-1, 0.1, length = 100), seq(1, 5, length = 100)))
  my_palette <- colorRampPalette(c("yellow", "black", "black", "purple"))(length(breaks) - 1)
  
  # Create heatmap
  heatmap.2(t(current_label_HM),
            Rowv = TRUE,
            Colv = TRUE,
            col = my_palette,
            breaks = breaks,
            density.info = "none",
            trace = "none",
            dendrogram = c("column"), 
            symm = FALSE,
            symkey = FALSE,
            symbreaks = TRUE,
            labRow = FALSE,
            labCol = FALSE,
            cexRow = 0.8,
            cexCol = 0.1,
            margins = c(8, 2),
            key.title = "1",
            key.xlab = "Z Score",
            sepcolor = c("black"),
            sepwidth = c(0.05, 0.05),
            distfun = function(x) dist(x, method = "euclidean"),
            hclust = function(x) hclust(x, method = "ward.D2"))
  
  dev.off()  # Close the PNG device
}
```
