## Supplementary material for "SAMP-Score: A morphology-based machine learning classification method for screening pro-senescence compounds in p16 positive cancer cells": HTML Walkthrough Guides for Code: Supplemental Data 1_3_Building_Models_Traditional_Threshold.html

Building-Models\_1.92\_Notebook.knit


### 1. Define Ground Truth

```
library("gplots")
```

```
## 
## Attaching package: 'gplots'
```

```
## The following object is masked from 'package:stats':
## 
##     lowess
```

```
library("RColorBrewer")
library("matrixStats")
library("plyr")
```

```
## 
## Attaching package: 'plyr'
```

```
## The following object is masked from 'package:matrixStats':
## 
##     count
```

```
library("dplyr")
```

```
## 
## Attaching package: 'dplyr'
```

```
## The following objects are masked from 'package:plyr':
## 
##     arrange, count, desc, failwith, id, mutate, rename, summarise,
##     summarize
```

```
## The following object is masked from 'package:matrixStats':
## 
##     count
```

```
## The following objects are masked from 'package:stats':
## 
##     filter, lag
```

```
## The following objects are masked from 'package:base':
## 
##     intersect, setdiff, setequal, union
```

```
library("data.table")
```

```
## 
## Attaching package: 'data.table'
```

```
## The following objects are masked from 'package:dplyr':
## 
##     between, first, last
```

```
library("stringr")
library("ggplot2")
library(glmnet)
```

```
## Loading required package: Matrix
```

```
## Loaded glmnet 4.1-8
```

```
library(caret)
```

```
## Loading required package: lattice
```

```
library(pROC)
```

```
## Type 'citation("pROC")' for a citation.
```

```
## 
## Attaching package: 'pROC'
```

```
## The following objects are masked from 'package:stats':
## 
##     cov, smooth, var
```

```
library(e1071)  # Contains functions for SVM
library(randomForest)
```

```
## randomForest 4.7-1.1
```

```
## Type rfNews() to see new features/changes/bug fixes.
```

```
## 
## Attaching package: 'randomForest'
```

```
## The following object is masked from 'package:ggplot2':
## 
##     margin
```

```
## The following object is masked from 'package:dplyr':
## 
##     combine
```

```
library(mda)
```

```
## Loading required package: class
```

```
## Loaded mda 0.5-4
```

```
library(neuralnet)
```

```
## 
## Attaching package: 'neuralnet'
```

```
## The following object is masked from 'package:dplyr':
## 
##     compute
```

```
# Upload your file 
all <- read.csv("All_Z_Scores.csv")


#Remove Blanks
all_small <- all %>%
  filter(!grepl("Blank", Gene.Symbol))

### Sanity check - NA removed in previous stage
all_removed <- all %>%
  filter(grepl("Blank", Gene.Symbol))

# Update DF ###
all <- all_small

#Deal with duplicate Gene Entries (12 in total)#
all <- all %>%
  group_by(RefSeq.Accession.Number) %>%
  mutate(RefSeq.Accession.Number = ifelse(row_number() == 1, RefSeq.Accession.Number, paste0(RefSeq.Accession.Number, "_A")))

#Add Ground Truth - Define filtering criteria
all <- all %>%
  mutate(Senescence_Status = ifelse(`Nuclei.Nuclei.Count.wv1` < -1.96 & Cells.Area.wv3 > 1.96, "Sen", "NonSen"))
```

### 2. Data Partitioning

```
# Load required libraries
library(ROSE)  # For undersampling and oversampling
```

```
## Loaded ROSE 0.0-4
```

```
library(caret)  # For creating data partitions
library(dplyr)  # For data manipulation

#Tidy Variables
All_Tidy <- all[,-c(1:5)]

# Step 1: Split the data into train and test sets (80-20 split)
set.seed(123)  # For reproducibility
trainIndex <- createDataPartition(All_Tidy$Senescence_Status, p = 0.8, list = FALSE)
trainData <- All_Tidy[trainIndex,]
testData <- All_Tidy[-trainIndex,]

# Add a unique identifier to the training data
trainData <- trainData %>%
  ungroup() %>%
  mutate(id = row_number())

# Checking the distribution of the target variable in train and test sets
cat("Distribution in Training Set:\n")
```

```
## Distribution in Training Set:
```

```
print(table(trainData$Senescence_Status))
```

```
## 
## NonSen    Sen 
##  16908    420
```

```
cat("Distribution in Test Set:\n")
```

```
## Distribution in Test Set:
```

```
print(table(testData$Senescence_Status))
```

```
## 
## NonSen    Sen 
##   4226    104
```

```
# Step 2: Perform undersampling to balance the classes
set.seed(123)
undersample_result <- ovun.sample(Senescence_Status ~ ., data = trainData, method = "under", p = 0.5)$data

# Checking the distribution after undersampling
cat("Distribution after Undersampling:\n")
```

```
## Distribution after Undersampling:
```

```
print(table(undersample_result$Senescence_Status))
```

```
## 
## NonSen    Sen 
##    408    420
```

```
# Step 3: Track and Extract Removed Observations
# Identify removed observations
# Removed IDs are those in trainData but not in undersample_result
removed_ids <- setdiff(trainData$id, undersample_result$id)

# Extract removed observations from the original training data
recycled_data <- trainData %>% filter(id %in% removed_ids)

# Remove the unique identifier column from all datasets
trainData <- trainData %>% select(-id)
undersample_result <- undersample_result %>% select(-id)
recycled_data <- recycled_data %>% select(-id)

# Combine recycled data with the test data
combined_testData <- bind_rows(testData, recycled_data)

# Checking the distribution in the combined test set
cat("Distribution in Combined Test Set:\n")
```

```
## Distribution in Combined Test Set:
```

```
print(table(combined_testData$Senescence_Status))
```

```
## 
## NonSen    Sen 
##  20726    104
```

```
# Split the Undersampled Training Data into Training and Validation
set.seed(123)
trainIndex_under <- createDataPartition(undersample_result$Senescence_Status, p = 0.5, list = FALSE)
trainData_part1 <- undersample_result[trainIndex_under,]
trainData_part2 <- undersample_result[-trainIndex_under,]

# Checking the distribution of the target variable in each part
cat("Distribution in Part 1 of Undersampled Data:\n")
```

```
## Distribution in Part 1 of Undersampled Data:
```

```
print(table(trainData_part1$Senescence_Status))
```

```
## 
## NonSen    Sen 
##    204    210
```

```
cat("Distribution in Part 2 of Undersampled Data:\n")
```

```
## Distribution in Part 2 of Undersampled Data:
```

```
print(table(trainData_part2$Senescence_Status))
```

```
## 
## NonSen    Sen 
##    204    210
```

### 3. Lasso Regularisation

```
# Load required libraries
library(glmnet)
library(caret)
library(pROC)
library(ggplot2)

# Ensure Senescence_Status is a factor (for classification)
trainData_part1$Senescence_Status <- as.factor(trainData_part1$Senescence_Status)
trainData_part2$Senescence_Status <- as.factor(trainData_part2$Senescence_Status)
combined_testData$Senescence_Status <- as.factor(combined_testData$Senescence_Status)

# Prepare training and test data for Lasso
x_train <- as.matrix(trainData_part1[, -which(names(trainData_part1) == "Senescence_Status")])
y_train <- trainData_part1$Senescence_Status
x_test <- as.matrix(combined_testData[, -which(names(combined_testData) == "Senescence_Status")])
y_test <- combined_testData$Senescence_Status

# Fit the Lasso model on trainData_part1
set.seed(123)
lasso_model <- glmnet(x_train, y_train, family = "binomial", alpha = 1)

# Use cross-validation to determine the best lambda
cv_model <- cv.glmnet(x_train, y_train, family = "binomial", alpha = 1)
best_lambda <- cv_model$lambda.min  # or cv_model$lambda.1se for simplicity

# Refit the model using the best lambda value
final_model <- glmnet(x_train, y_train, family = "binomial", alpha = 1, lambda = best_lambda)

# Make predictions on trainData_part2 using the Lasso model
predicted_probabilities_part2 <- predict(final_model, newx = as.matrix(trainData_part2[, -which(names(trainData_part2) == "Senescence_Status")]), s = best_lambda, type = "response")
predicted_classes_part2 <- ifelse(predicted_probabilities_part2 > 0.5, "Sen", "NonSen")

# Create the meta-model training data frame
lasso_meta_model_data_training <- data.frame(
  Lasso_Prob = as.vector(predicted_probabilities_part2),
  Senescence_Status = trainData_part2$Senescence_Status,
  Predicted_Status = predicted_classes_part2
)

# Save the meta-model training data for further use
write.csv(lasso_meta_model_data_training, "lasso_meta_model_data_training.csv", row.names = FALSE)

## Now we are going to apply the model to testing data and evaluate

# Make predictions on testing data using the Lasso model
predicted_probabilities_test <- predict(final_model, newx = x_test, s = best_lambda, type = "response")
predicted_classes_test <- ifelse(predicted_probabilities_test > 0.5, "Sen", "NonSen")

# Create the meta-model testing data frame
lasso_meta_model_data_testing <- data.frame(
  Lasso_Prob = as.vector(predicted_probabilities_test),
  Senescence_Status = y_test,
  Predicted_Status = predicted_classes_test
)

# Save the meta-model testing data for further use
write.csv(lasso_meta_model_data_testing, "lasso_meta_model_data_testing.csv", row.names = FALSE)

# Convert predicted_classes and actual classes to factors
combined_testData$predicted_classes <- factor(predicted_classes_test, levels = c("NonSen", "Sen"))
combined_testData$Senescence_Status <- factor(combined_testData$Senescence_Status)

# Create the confusion matrix for Lasso on combined_testData
lasso_confusion_mtx_test <- confusionMatrix(data = combined_testData$predicted_classes, reference = combined_testData$Senescence_Status)
print(lasso_confusion_mtx_test)
```

```
## Confusion Matrix and Statistics
## 
##           Reference
## Prediction NonSen   Sen
##     NonSen  20121     1
##     Sen       605   103
##                                           
##                Accuracy : 0.9709          
##                  95% CI : (0.9685, 0.9731)
##     No Information Rate : 0.995           
##     P-Value [Acc > NIR] : 1               
##                                           
##                   Kappa : 0.2471          
##                                           
##  Mcnemar's Test P-Value : <2e-16          
##                                           
##             Sensitivity : 0.9708          
##             Specificity : 0.9904          
##          Pos Pred Value : 1.0000          
##          Neg Pred Value : 0.1455          
##              Prevalence : 0.9950          
##          Detection Rate : 0.9660          
##    Detection Prevalence : 0.9660          
##       Balanced Accuracy : 0.9806          
##                                           
##        'Positive' Class : NonSen          
##
```

```
# Calculate precision, recall, F1-score, specificity, FPR
precision_test <- lasso_confusion_mtx_test$byClass["Pos Pred Value"]
recall_test <- lasso_confusion_mtx_test$byClass["Sensitivity"]
f1_score_test <- lasso_confusion_mtx_test$byClass["F1"]
specificity_test <- lasso_confusion_mtx_test$byClass["Neg Pred Value"]
fpr_test <- 1 - specificity_test

# Compute ROC curve and AUC
roc_curve_test <- roc(combined_testData$Senescence_Status, as.numeric(predicted_probabilities_test))
```

```
## Setting levels: control = NonSen, case = Sen
```

```
## Setting direction: controls < cases
```

```
auc_roc_test <- auc(roc_curve_test)

# Ensure the 'ROCs' directory exists
if (!dir.exists("ROCs")) {
  dir.create("ROCs")
}

# Create the ROC plot
roc_plot <- ggroc(roc_curve_test, legacy.axes = TRUE, alpha = 0.5, colour = "blue", linetype = 1, size = 1) +
  labs(title = "ROC Curve for Lasso Model", x = "False Positive Rate", y = "True Positive Rate") +
  geom_segment(aes(x = 0, xend = 1, y = 0, yend = 1), color = "black", linetype = "dashed")

# Save the ROC plot
ggsave("ROCs/roc_curve_lasso.png", plot = roc_plot, width = 8, height = 6)

# Store metrics and confusion matrix breakdown for combined_testData in a data frame
lasso_metrics_df_test <- data.frame(
  Model = "Lasso Model (Combined Test Data)",
  Accuracy = lasso_confusion_mtx_test$overall["Accuracy"],
  Precision = precision_test,
  Recall = recall_test,
  F1_Score = f1_score_test,
  AUC = auc_roc_test,
  Neg_Pred_Value = specificity_test,
  TP = lasso_confusion_mtx_test$table["Sen", "Sen"],
  TN = lasso_confusion_mtx_test$table["NonSen", "NonSen"],
  FN = lasso_confusion_mtx_test$table["NonSen", "Sen"],
  FP = lasso_confusion_mtx_test$table["Sen", "NonSen"]
)

# Save the metrics for combined_testData to a CSV file
write.csv(lasso_metrics_df_test, "lasso_model_metrics.csv", row.names = FALSE)

# Plot the coefficients of the Lasso model
plot_coef <- function(model, predictors, lambda) {
  # Extract coefficients
  coef <- coef(model, s = lambda)
  
  # Create dataframe for coefficients
  coef_df <- data.frame(Predictor = predictors, Coefficient = as.numeric(coef[-1]))
  
  # Plot coefficients
  ggplot(coef_df, aes(x = reorder(Predictor, Coefficient), y = Coefficient)) +
    geom_bar(stat = "identity", fill = "skyblue") +
    coord_flip() +
    labs(title = "Coefficient Plot for Lasso Logistic Regression",
         x = "Predictor", y = "Coefficient") +
    theme_minimal() +
    theme(axis.text.x = element_text(angle = 45, hjust = 1))
}

# Plot coefficients of the Lasso model
coef_plot <- plot_coef(final_model, colnames(x_train), best_lambda)

# Save the coefficient plot
ggsave("ROCs/coef_plot_lasso.png", plot = coef_plot, width = 8, height = 6)
```

### 4. Logistic Regression

```
# Load required libraries
library(caret)
library(pROC)
library(ggplot2)
library(glmnet)

# Ensure Senescence_Status is a factor (for classification)
trainData_part1$Senescence_Status <- as.factor(trainData_part1$Senescence_Status)
trainData_part2$Senescence_Status <- as.factor(trainData_part2$Senescence_Status)
combined_testData$Senescence_Status <- as.factor(combined_testData$Senescence_Status)

# Prepare training and test data for Logistic Regression
x_train <- as.matrix(trainData_part1[, -which(names(trainData_part1) == "Senescence_Status")])
y_train <- trainData_part1$Senescence_Status
x_test <- as.matrix(combined_testData[, -which(names(combined_testData) == "Senescence_Status")])
y_test <- combined_testData$Senescence_Status

# Fit the logistic regression model on trainData_part1
set.seed(123)
model_binary <- glm(Senescence_Status ~ ., data = trainData_part1, family = "binomial")
```

```
## Warning: glm.fit: algorithm did not converge
```

```
## Warning: glm.fit: fitted probabilities numerically 0 or 1 occurred
```

```
# Make predictions on trainData_part2 using the logistic regression model
predicted_probabilities_part2 <- predict(model_binary, newdata = trainData_part2, type = "response")
predicted_classes_part2 <- ifelse(predicted_probabilities_part2 > 0.5, "Sen", "NonSen")

# Create the meta-model training data frame
logistic_meta_model_data_training <- data.frame(
  logistic_Prob = as.vector(predicted_probabilities_part2),
  Senescence_Status = trainData_part2$Senescence_Status,
  Predicted_Status = predicted_classes_part2
)

# Save the meta-model training data for further use
write.csv(logistic_meta_model_data_training, "logistic_meta_model_data_training.csv", row.names = FALSE)

## Now we are going to apply the model to testing data and evaluate

# Make predictions on testing data using the logistic regression model
predicted_probabilities_test <- predict(model_binary, newdata = combined_testData, type = "response")
predicted_classes_test <- ifelse(predicted_probabilities_test > 0.5, "Sen", "NonSen")

# Create the meta-model testing data frame
logistic_meta_model_data_testing <- data.frame(
  logistic_Prob = as.vector(predicted_probabilities_test),
  Senescence_Status = y_test,
  Predicted_Status = predicted_classes_test
)

# Save the meta-model testing data for further use
write.csv(logistic_meta_model_data_testing, "logistic_meta_model_data_testing.csv", row.names = FALSE)

# Convert predicted_classes and actual classes to factors
combined_testData$predicted_classes <- factor(predicted_classes_test, levels = c("NonSen", "Sen"))
combined_testData$Senescence_Status <- factor(combined_testData$Senescence_Status)

# Create the confusion matrix for Logistic Regression on combined_testData
logistic_confusion_mtx_test <- confusionMatrix(data = combined_testData$predicted_classes, reference = combined_testData$Senescence_Status)
print(logistic_confusion_mtx_test)
```

```
## Confusion Matrix and Statistics
## 
##           Reference
## Prediction NonSen   Sen
##     NonSen  20037     8
##     Sen       689    96
##                                          
##                Accuracy : 0.9665         
##                  95% CI : (0.964, 0.9689)
##     No Information Rate : 0.995          
##     P-Value [Acc > NIR] : 1              
##                                          
##                   Kappa : 0.209          
##                                          
##  Mcnemar's Test P-Value : <2e-16         
##                                          
##             Sensitivity : 0.9668         
##             Specificity : 0.9231         
##          Pos Pred Value : 0.9996         
##          Neg Pred Value : 0.1223         
##              Prevalence : 0.9950         
##          Detection Rate : 0.9619         
##    Detection Prevalence : 0.9623         
##       Balanced Accuracy : 0.9449         
##                                          
##        'Positive' Class : NonSen         
##
```

```
# Calculate precision, recall, F1-score, specificity, FPR
precision_test <- logistic_confusion_mtx_test$byClass["Pos Pred Value"]
recall_test <- logistic_confusion_mtx_test$byClass["Sensitivity"]
f1_score_test <- logistic_confusion_mtx_test$byClass["F1"]
specificity_test <- logistic_confusion_mtx_test$byClass["Neg Pred Value"]
fpr_test <- 1 - specificity_test

# Compute ROC curve and AUC
roc_curve_test <- roc(combined_testData$Senescence_Status, as.numeric(predicted_probabilities_test))
```

```
## Setting levels: control = NonSen, case = Sen
```

```
## Setting direction: controls < cases
```

```
auc_roc_test <- auc(roc_curve_test)

# Ensure the 'ROCs' directory exists
if (!dir.exists("ROCs")) {
  dir.create("ROCs")
}

# Create the ROC plot
roc_plot <- ggroc(roc_curve_test, legacy.axes = TRUE, alpha = 0.5, colour = "blue", linetype = 1, size = 1) +
  labs(title = "ROC Curve for Logistic Regression", x = "False Positive Rate", y = "True Positive Rate") +
  geom_segment(aes(x = 0, xend = 1, y = 0, yend = 1), color = "black", linetype = "dashed")

# Save the ROC plot
ggsave("ROCs/roc_curve_logistic.png", plot = roc_plot, width = 8, height = 6)

# Store metrics and confusion matrix breakdown for combined_testData in a data frame
logistic_metrics_df_test <- data.frame(
  Model = "Logistic Model (Combined Test Data)",
  Accuracy = logistic_confusion_mtx_test$overall["Accuracy"],
  Precision = precision_test,
  Recall = recall_test,
  F1_Score = f1_score_test,
  AUC = auc_roc_test,
  Neg_Pred_Value = specificity_test,
  TP = logistic_confusion_mtx_test$table["Sen", "Sen"],
  TN = logistic_confusion_mtx_test$table["NonSen", "NonSen"],
  FN = logistic_confusion_mtx_test$table["NonSen", "Sen"],
  FP = logistic_confusion_mtx_test$table["Sen", "NonSen"]
)

# Save the metrics for combined_testData to a CSV file
write.csv(logistic_metrics_df_test, "logistic_model_metrics.csv", row.names = FALSE)
```

### 5. Elastic Nets

```
set.seed(123)

# Load required libraries
library(glmnet)
library(caret)
library(pROC)
library(ggplot2)

# Ensure Senescence_Status is a factor (for classification)
trainData_part1$Senescence_Status <- as.factor(trainData_part1$Senescence_Status)
trainData_part2$Senescence_Status <- as.factor(trainData_part2$Senescence_Status)
combined_testData$Senescence_Status <- as.factor(combined_testData$Senescence_Status)

# Prepare training and test data for Elastic Net
x_train <- as.matrix(trainData_part1[, -which(names(trainData_part1) == "Senescence_Status")])
y_train <- trainData_part1$Senescence_Status

# Ensure the test data has the same columns as the training data
x_test <- as.matrix(combined_testData[, colnames(x_train)])
y_test <- combined_testData$Senescence_Status

# Fit the Elastic Net model on trainData_part1
elastic_net_model <- cv.glmnet(x_train, y_train, family = "binomial", alpha = 0.5)

# Use cross-validation to determine the best lambda
best_lambda <- elastic_net_model$lambda.min  

# Refit the model using the best lambda value
final_elastic_net_model <- glmnet(x_train, y_train, family = "binomial", alpha = 0.5, lambda = best_lambda)

# Make predictions on trainData_part2 using the Elastic Net model
x_train_part2 <- as.matrix(trainData_part2[, colnames(x_train)])  # Ensure same columns
predicted_probabilities_part2 <- predict(final_elastic_net_model, newx = x_train_part2, s = best_lambda, type = "response")
predicted_classes_part2 <- ifelse(predicted_probabilities_part2 > 0.5, "Sen", "NonSen")

# Create the meta-model training data frame
en_meta_model_data_training <- data.frame(
  en_Probabilities = as.vector(predicted_probabilities_part2),
  Senescence_Status = trainData_part2$Senescence_Status,
  Predicted_Status = predicted_classes_part2
)

# Save the meta-model training data for further use
write.csv(en_meta_model_data_training, "en_meta_model_data_training.csv", row.names = FALSE)

## Now we are going to apply the model to testing data and evaluate

# Make predictions on testing data using the Elastic Net model
predicted_probabilities_test <- predict(final_elastic_net_model, newx = x_test, s = best_lambda, type = "response")
predicted_classes_test <- ifelse(predicted_probabilities_test > 0.5, "Sen", "NonSen")

# Create the meta-model testing data frame
en_meta_model_data_testing <- data.frame(
  en_Probabilities = as.vector(predicted_probabilities_test),
  Senescence_Status = y_test,
  Predicted_Status = predicted_classes_test
)

# Save the meta-model testing data for further use
write.csv(en_meta_model_data_testing, "en_meta_model_data_testing.csv", row.names = FALSE)

# Convert predicted_classes and actual classes to factors
combined_testData$predicted_classes <- factor(predicted_classes_test, levels = c("NonSen", "Sen"))
combined_testData$Senescence_Status <- factor(combined_testData$Senescence_Status)

# Create the confusion matrix for Elastic Net on combined_testData
en_confusion_mtx_test <- confusionMatrix(data = combined_testData$predicted_classes, reference = combined_testData$Senescence_Status)
print(en_confusion_mtx_test)
```

```
## Confusion Matrix and Statistics
## 
##           Reference
## Prediction NonSen   Sen
##     NonSen  20075     1
##     Sen       651   103
##                                          
##                Accuracy : 0.9687         
##                  95% CI : (0.9662, 0.971)
##     No Information Rate : 0.995          
##     P-Value [Acc > NIR] : 1              
##                                          
##                   Kappa : 0.2334         
##                                          
##  Mcnemar's Test P-Value : <2e-16         
##                                          
##             Sensitivity : 0.9686         
##             Specificity : 0.9904         
##          Pos Pred Value : 1.0000         
##          Neg Pred Value : 0.1366         
##              Prevalence : 0.9950         
##          Detection Rate : 0.9638         
##    Detection Prevalence : 0.9638         
##       Balanced Accuracy : 0.9795         
##                                          
##        'Positive' Class : NonSen         
##
```

```
# Calculate precision, recall, F1-score, specificity, FPR
precision_test <- en_confusion_mtx_test$byClass["Pos Pred Value"]
recall_test <- en_confusion_mtx_test$byClass["Sensitivity"]
f1_score_test <- en_confusion_mtx_test$byClass["F1"]
specificity_test <- en_confusion_mtx_test$byClass["Neg Pred Value"]
fpr_test <- 1 - specificity_test

# Compute ROC curve and AUC for Elastic Net
roc_curve_test <- roc(combined_testData$Senescence_Status, as.numeric(predicted_probabilities_test))
```

```
## Setting levels: control = NonSen, case = Sen
```

```
## Setting direction: controls < cases
```

```
auc_roc_test <- auc(roc_curve_test)

# Ensure the 'ROCs' directory exists
if (!dir.exists("ROCs")) {
  dir.create("ROCs")
}

# Create the ROC plot for Elastic Net
roc_plot <- ggroc(roc_curve_test, legacy.axes = TRUE, alpha = 0.5, colour = "blue", linetype = 1, size = 1) +
  labs(title = "ROC Curve on Combined Test Data (Elastic Net)", x = "False Positive Rate", y = "True Positive Rate") +
  geom_segment(aes(x = 0, xend = 1, y = 0, yend = 1), color = "black", linetype = "dashed")

# Save the ROC plot
ggsave("ROCs/roc_curve_elastic_net.png", plot = roc_plot, width = 8, height = 6)

# Store metrics and confusion matrix breakdown for combined_testData in a data frame
en_metrics_df_test <- data.frame(
  Model = "Elastic Net Model (Combined Test Data)",
  Accuracy = en_confusion_mtx_test$overall["Accuracy"],
  Precision = precision_test,
  Recall = recall_test,
  F1_Score = f1_score_test,
  AUC = auc_roc_test,
  Neg_Pred_Value = specificity_test,
  TP = en_confusion_mtx_test$table["Sen", "Sen"],
  TN = en_confusion_mtx_test$table["NonSen", "NonSen"],
  FN = en_confusion_mtx_test$table["NonSen", "Sen"],
  FP = en_confusion_mtx_test$table["Sen", "NonSen"]
)

# Save the metrics for combined_testData to a CSV file
write.csv(en_metrics_df_test, "en_model_metrics.csv", row.names = FALSE)
```

#6. Random Forrest

```
# Load required libraries
library(randomForest)
library(caret)
library(pROC)
library(ggplot2)

# Ensure Senescence_Status is a factor (for classification)
trainData_part1$Senescence_Status <- as.factor(trainData_part1$Senescence_Status)
trainData_part2$Senescence_Status <- as.factor(trainData_part2$Senescence_Status)
combined_testData$Senescence_Status <- as.factor(combined_testData$Senescence_Status)

# Fit the Random Forest model on trainData_part1
set.seed(123)
rf_model <- randomForest(x = trainData_part1[, -which(names(trainData_part1) == "Senescence_Status")],
                         y = trainData_part1$Senescence_Status,
                         ntree = 500,
                         type = "classification")

# Make predictions on trainData_part2 using the Random Forest model
rf_predictions_part2 <- predict(rf_model, newdata = trainData_part2)

# Get class probabilities for the positive class
rf_probabilities_part2 <- predict(rf_model, newdata = trainData_part2, type = "prob")[, "Sen"]

# Create the meta-model training data frame
rf_meta_model_data_training <- data.frame(
  rf_Prob = as.vector(rf_probabilities_part2),
  Senescence_Status = trainData_part2$Senescence_Status,
  Predicted_Status = rf_predictions_part2
)

# Save the meta-model training data for further use
write.csv(rf_meta_model_data_training, "rf_meta_model_data_training.csv", row.names = FALSE)

## Now we are going to apply the model to testing data and evaluate 

# Make predictions on testing data using the Random Forest model
rf_predictions_test <- predict(rf_model, newdata = combined_testData)

# Get class probabilities for the positive class on testing data
rf_probabilities_test <- predict(rf_model, newdata = combined_testData, type = "prob")[, "Sen"]

# Convert predicted_classes and actual classes to factors
combined_testData$predicted_classes <- rf_predictions_test
combined_testData$predicted_classes <- factor(combined_testData$predicted_classes)
combined_testData$Senescence_Status <- factor(combined_testData$Senescence_Status)

# Create the confusion matrix for Random Forest on combined_testData
rf_confusion_mtx_test <- confusionMatrix(data = combined_testData$predicted_classes, reference = combined_testData$Senescence_Status)
print(rf_confusion_mtx_test)
```

```
## Confusion Matrix and Statistics
## 
##           Reference
## Prediction NonSen   Sen
##     NonSen  20448     1
##     Sen       278   103
##                                          
##                Accuracy : 0.9866         
##                  95% CI : (0.985, 0.9881)
##     No Information Rate : 0.995          
##     P-Value [Acc > NIR] : 1              
##                                          
##                   Kappa : 0.4202         
##                                          
##  Mcnemar's Test P-Value : <2e-16         
##                                          
##             Sensitivity : 0.9866         
##             Specificity : 0.9904         
##          Pos Pred Value : 1.0000         
##          Neg Pred Value : 0.2703         
##              Prevalence : 0.9950         
##          Detection Rate : 0.9817         
##    Detection Prevalence : 0.9817         
##       Balanced Accuracy : 0.9885         
##                                          
##        'Positive' Class : NonSen         
##
```

```
# Calculate precision, recall, F1-score, specificity, FPR
precision_test <- rf_confusion_mtx_test$byClass["Pos Pred Value"]
recall_test <- rf_confusion_mtx_test$byClass["Sensitivity"]
f1_score_test <- rf_confusion_mtx_test$byClass["F1"]
specificity_test <- rf_confusion_mtx_test$byClass["Neg Pred Value"]
fpr_test <- 1 - specificity_test

# Compute ROC curve and AUC
roc_curve_test <- roc(combined_testData$Senescence_Status, rf_probabilities_test)
```

```
## Setting levels: control = NonSen, case = Sen
```

```
## Setting direction: controls < cases
```

```
auc_roc_test <- auc(roc_curve_test)

# Ensure the 'ROCs' directory exists
if (!dir.exists("ROCs")) {
  dir.create("ROCs")
}

# Create the ROC plot
roc_plot <- ggroc(roc_curve_test, legacy.axes = TRUE, alpha = 0.5, colour = "blue", linetype = 1, size = 1) +
  labs(title = "ROC Curve for Random Forest", x = "False Positive Rate", y = "True Positive Rate") +
  geom_segment(aes(x = 0, xend = 1, y = 0, yend = 1), color = "black", linetype = "dashed")

# Save the ROC plot
ggsave("ROCs/roc_curve_rf.png", plot = roc_plot, width = 8, height = 6)

# Store metrics and confusion matrix breakdown for combined_testData in a data frame
rf_metrics_df_test <- data.frame(
  Model = "Random Forest (Combined Test Data)",
  Accuracy = rf_confusion_mtx_test$overall["Accuracy"],
  Precision = precision_test,
  Recall = recall_test,
  F1_Score = f1_score_test,
  AUC = auc_roc_test,
  Neg_Pred_Value = specificity_test,
  TP = rf_confusion_mtx_test$table["Sen", "Sen"],
  TN = rf_confusion_mtx_test$table["NonSen", "NonSen"],
  FN = rf_confusion_mtx_test$table["NonSen", "Sen"],
  FP = rf_confusion_mtx_test$table["Sen", "NonSen"]
)

# Save the metrics for combined_testData to a CSV file
write.csv(rf_metrics_df_test, "rf_model_metrics.csv", row.names = FALSE)

# Plot variable importance
var_importance <- importance(rf_model)
var_importance_df <- data.frame(Variable = rownames(var_importance), Importance = var_importance[, "MeanDecreaseGini"])
var_importance_df <- var_importance_df[order(var_importance_df$Importance, decreasing = TRUE), ]

var_importance_plot <- ggplot(var_importance_df, aes(x = reorder(Variable, Importance), y = Importance)) +
  geom_bar(stat = "identity", fill = "skyblue") +
  coord_flip() +
  labs(title = "Variable Importance for Random Forest", x = "Variable", y = "Importance")

# Save the variable importance plot
ggsave("ROCs/variable_importance_rf.png", plot = var_importance_plot, width = 8, height = 6)

# Create the meta-model testing data frame
rf_meta_model_data_testing <- data.frame(
  rf_Prob = as.vector(rf_probabilities_test),
  Senescence_Status = combined_testData$Senescence_Status,
  Predicted_Status = rf_predictions_test
)

# Save the meta-model testing data for further use
write.csv(rf_meta_model_data_testing, "rf_meta_model_data_testing.csv", row.names = FALSE)
```

### 7. SVMs

```
set.seed(123)

# Load required libraries
library(e1071)
library(caret)
library(pROC)
library(ggplot2)

# Ensure Senescence_Status is a factor (for classification)
trainData_part1$Senescence_Status <- as.factor(trainData_part1$Senescence_Status)
trainData_part2$Senescence_Status <- as.factor(trainData_part2$Senescence_Status)
combined_testData$Senescence_Status <- as.factor(combined_testData$Senescence_Status)

# Prepare training and test data for SVM
x_train <- trainData_part1[, -which(names(trainData_part1) == "Senescence_Status")]
y_train <- trainData_part1$Senescence_Status
x_test <- combined_testData[, -which(names(combined_testData) == "Senescence_Status")]
y_test <- combined_testData$Senescence_Status

# Fit the SVM model on trainData_part1
svm_model <- svm(x_train, y_train, kernel = "radial", probability = TRUE)

# Ensure trainData_part2 and combined_testData have the same features as x_train
x_train_part2 <- trainData_part2[, colnames(x_train), drop = FALSE]

# Make predictions on trainData_part2 using the SVM model
svm_predictions_part2 <- predict(svm_model, newdata = x_train_part2, probability = TRUE)
svm_probabilities_part2 <- attr(svm_predictions_part2, "probabilities")[, "Sen"]
svm_predicted_classes_part2 <- ifelse(svm_probabilities_part2 > 0.5, "Sen", "NonSen")

# Create the meta-model training data frame
svm_meta_model_data_training <- data.frame(
  svm_Prob = as.vector(svm_probabilities_part2),
  Senescence_Status = as.factor(trainData_part2$Senescence_Status),
  Predicted_Status = svm_predicted_classes_part2
)

# Save the meta-model training data for further use
write.csv(svm_meta_model_data_training, "svm_meta_model_data_training.csv", row.names = FALSE)

## Now we are going to apply the model to testing data and evaluate

# Make predictions on combined_testData using the SVM model
x_test_combined <- combined_testData[, colnames(x_train), drop = FALSE]  # Ensure same columns as x_train
svm_predictions_test <- predict(svm_model, newdata = x_test_combined, probability = TRUE)
svm_probabilities_test <- attr(svm_predictions_test, "probabilities")[, "Sen"]
svm_predicted_classes_test <- ifelse(svm_probabilities_test > 0.5, "Sen", "NonSen")

# Create the meta-model testing data frame
svm_meta_model_data_testing <- data.frame(
  svm_Prob = as.vector(svm_probabilities_test),
  Senescence_Status = y_test,
  Predicted_Status = svm_predicted_classes_test
)

# Save the meta-model testing data for further use
write.csv(svm_meta_model_data_testing, "svm_meta_model_data_testing.csv", row.names = FALSE)

# Convert predicted_classes and actual classes to factors
combined_testData$predicted_classes <- factor(svm_predicted_classes_test, levels = c("NonSen", "Sen"))
combined_testData$Senescence_Status <- factor(combined_testData$Senescence_Status)

# Create the confusion matrix for SVM on combined_testData
svm_confusion_mtx_test <- confusionMatrix(data = combined_testData$predicted_classes, reference = combined_testData$Senescence_Status)
print(svm_confusion_mtx_test)
```

```
## Confusion Matrix and Statistics
## 
##           Reference
## Prediction NonSen   Sen
##     NonSen  19871     2
##     Sen       855   102
##                                           
##                Accuracy : 0.9589          
##                  95% CI : (0.9561, 0.9615)
##     No Information Rate : 0.995           
##     P-Value [Acc > NIR] : 1               
##                                           
##                   Kappa : 0.1849          
##                                           
##  Mcnemar's Test P-Value : <2e-16          
##                                           
##             Sensitivity : 0.9587          
##             Specificity : 0.9808          
##          Pos Pred Value : 0.9999          
##          Neg Pred Value : 0.1066          
##              Prevalence : 0.9950          
##          Detection Rate : 0.9540          
##    Detection Prevalence : 0.9541          
##       Balanced Accuracy : 0.9698          
##                                           
##        'Positive' Class : NonSen          
##
```

```
# Calculate precision, recall, F1-score, specificity, FPR
precision_test <- svm_confusion_mtx_test$byClass["Pos Pred Value"]
recall_test <- svm_confusion_mtx_test$byClass["Sensitivity"]
f1_score_test <- svm_confusion_mtx_test$byClass["F1"]
specificity_test <- svm_confusion_mtx_test$byClass["Neg Pred Value"]
fpr_test <- 1 - specificity_test

# Compute ROC curve and AUC for SVM
roc_curve_test <- roc(combined_testData$Senescence_Status, svm_probabilities_test)
```

```
## Setting levels: control = NonSen, case = Sen
```

```
## Setting direction: controls < cases
```

```
auc_roc_test <- auc(roc_curve_test)

# Calculate Cohen's Kappa
kappa_test <- svm_confusion_mtx_test$overall["Kappa"]

# Ensure the 'ROCs' directory exists
if (!dir.exists("ROCs")) {
  dir.create("ROCs")
}

# Create the ROC plot for SVM
roc_plot <- ggroc(roc_curve_test, legacy.axes = TRUE, alpha = 0.5, colour = "blue", linetype = 1, size = 1) +
  labs(title = "ROC Curve on Combined Test Data (SVM)", x = "False Positive Rate", y = "True Positive Rate") +
  geom_segment(aes(x = 0, xend = 1, y = 0, yend = 1), color = "black", linetype = "dashed")

# Save the ROC plot
ggsave("ROCs/roc_curve_svm.png", plot = roc_plot, width = 8, height = 6)

# Store metrics and confusion matrix breakdown for combined_testData in a data frame
SVM_metrics_df_test <- data.frame(
  Model = "SVM Model",
  Accuracy = svm_confusion_mtx_test$overall["Accuracy"],
  Precision = precision_test,
  Recall = recall_test,
  F1_Score = f1_score_test,
  AUC = auc_roc_test,
  Neg_Pred_Value = specificity_test,
  Kappa = kappa_test,
  TP = svm_confusion_mtx_test$table["Sen", "Sen"],
  TN = svm_confusion_mtx_test$table["NonSen", "NonSen"],
  FN = svm_confusion_mtx_test$table["NonSen", "Sen"],
  FP = svm_confusion_mtx_test$table["Sen", "NonSen"]
)

# Save the metrics for combined_testData to a CSV file
write.csv(SVM_metrics_df_test, "svm_model_metrics.csv", row.names = FALSE)
```

### 8. MDA

```
set.seed(123)

# Load required libraries
library(mda)
library(caret)
library(pROC)
library(ggplot2)

# Ensure Senescence_Status is a factor (for classification)
trainData_part1$Senescence_Status <- as.factor(trainData_part1$Senescence_Status)
trainData_part2$Senescence_Status <- as.factor(trainData_part2$Senescence_Status)
combined_testData$Senescence_Status <- as.factor(combined_testData$Senescence_Status)

# Prepare training data for MDA
x_train_part2 <- trainData_part2[, -which(names(trainData_part2) == "Senescence_Status")]
y_train_part2 <- trainData_part2$Senescence_Status

# Fit the MDA model on trainData_part2
mda_model <- mda(Senescence_Status ~ ., data = trainData_part2)

# Ensure test data has the same features as x_train_part2
x_test_combined <- combined_testData[, colnames(x_train_part2), drop = FALSE]

# Make predictions on trainData_part2 using the MDA model
predicted_probabilities_train_mda <- predict(mda_model, newdata = trainData_part2, type = "posterior")
predicted_classes_train_mda <- ifelse(predicted_probabilities_train_mda[, "Sen"] > 0.5, "Sen", "NonSen")

# Create and save the meta-model data frame for trainData_part2
mda_meta_model_training <- data.frame(
  mda_Prob = as.vector(predicted_probabilities_train_mda[, "Sen"]),
  Senescence_Status = as.factor(trainData_part2$Senescence_Status),
  Predicted_Status = predicted_classes_train_mda
)
write.csv(mda_meta_model_training, "mda_meta_model_training.csv", row.names = FALSE)

## Now we are going to apply the model to combined_testData and evaluate

# Make predictions on combined_testData using the MDA model
predicted_probabilities_test_mda <- predict(mda_model, newdata = combined_testData, type = "posterior")
predicted_classes_test_mda <- ifelse(predicted_probabilities_test_mda[, "Sen"] > 0.5, "Sen", "NonSen")

# Create and save the meta-model data frame for combined_testData
mda_meta_model_testing <- data.frame(
  mda_Prob = as.vector(predicted_probabilities_test_mda[, "Sen"]),
  Senescence_Status = as.factor(combined_testData$Senescence_Status),
  Predicted_Status = predicted_classes_test_mda
)
write.csv(mda_meta_model_testing, "mda_meta_model_testing.csv", row.names = FALSE)

# Calculate accuracy for MDA on combined_testData
accuracy_test_mda <- mean(predicted_classes_test_mda == combined_testData$Senescence_Status)
print(paste("MDA Accuracy on combined_testData:", accuracy_test_mda))
```

```
## [1] "MDA Accuracy on combined_testData: 0.92774843975036"
```

```
# Assign predicted classes to combined_testData
combined_testData_pred <- combined_testData
combined_testData_pred$predicted_classes <- predicted_classes_test_mda

# Convert predicted_classes and actual classes to factors
combined_testData_pred$predicted_classes <- factor(predicted_classes_test_mda, levels = c("NonSen", "Sen"))
combined_testData_pred$Senescence_Status <- factor(combined_testData_pred$Senescence_Status)

# Create confusion matrix for MDA on combined_testData
mda_confusion_mtx_test <- confusionMatrix(data = combined_testData_pred$predicted_classes, reference = combined_testData_pred$Senescence_Status)
print(mda_confusion_mtx_test)
```

```
## Confusion Matrix and Statistics
## 
##           Reference
## Prediction NonSen   Sen
##     NonSen  19223     2
##     Sen      1503   102
##                                           
##                Accuracy : 0.9277          
##                  95% CI : (0.9241, 0.9312)
##     No Information Rate : 0.995           
##     P-Value [Acc > NIR] : 1               
##                                           
##                   Kappa : 0.111           
##                                           
##  Mcnemar's Test P-Value : <2e-16          
##                                           
##             Sensitivity : 0.92748         
##             Specificity : 0.98077         
##          Pos Pred Value : 0.99990         
##          Neg Pred Value : 0.06355         
##              Prevalence : 0.99501         
##          Detection Rate : 0.92285         
##    Detection Prevalence : 0.92295         
##       Balanced Accuracy : 0.95413         
##                                           
##        'Positive' Class : NonSen          
##
```

```
# Calculate precision, recall, F1-score, specificity, FPR
precision_test <- mda_confusion_mtx_test$byClass["Pos Pred Value"]
recall_test <- mda_confusion_mtx_test$byClass["Sensitivity"]
f1_score_test <- mda_confusion_mtx_test$byClass["F1"]
specificity_test <- mda_confusion_mtx_test$byClass["Neg Pred Value"]
fpr_test <- 1 - specificity_test

# Compute ROC curve and AUC for MDA on combined_testData
roc_curve_mda_test <- roc(combined_testData_pred$Senescence_Status, predicted_probabilities_test_mda[, "Sen"])
```

```
## Setting levels: control = NonSen, case = Sen
```

```
## Setting direction: controls < cases
```

```
auc_roc_mda_test <- auc(roc_curve_mda_test)

# Calculate Cohen's Kappa
kappa_test <- mda_confusion_mtx_test$overall["Kappa"]

# Ensure the 'ROCs' directory exists
if (!dir.exists("ROCs")) {
  dir.create("ROCs")
}

# Create the ROC plot for MDA on combined_testData
roc_plot_mda <- ggroc(roc_curve_mda_test, legacy.axes = TRUE, alpha = 0.5, colour = "blue", linetype = 1, size = 1) +
  labs(title = "ROC Curve on Combined Test Data (MDA)", x = "False Positive Rate", y = "True Positive Rate") +
  geom_segment(aes(x = 0, xend = 1, y = 0, yend = 1), color = "black", linetype = "dashed")

# Save the ROC plot
ggsave("ROCs/roc_curve_mda.png", plot = roc_plot_mda, width = 8, height = 6)

# Store metrics and confusion matrix breakdown for combined_testData in a data frame
mda_metrics_df_test <- data.frame(
  Model = "MDA Model",
  Accuracy = accuracy_test_mda,
  Precision = precision_test,
  Recall = recall_test,
  F1_Score = f1_score_test,
  AUC = auc_roc_mda_test,
  Neg_Pred_Value = specificity_test,
  Kappa = kappa_test,
  TP = mda_confusion_mtx_test$table["Sen", "Sen"],
  TN = mda_confusion_mtx_test$table["NonSen", "NonSen"],
  FN = mda_confusion_mtx_test$table["NonSen", "Sen"],
  FP = mda_confusion_mtx_test$table["Sen", "NonSen"]
)

# Save the metrics for combined_testData to a CSV file
write.csv(mda_metrics_df_test, "mda_model_metrics.csv", row.names = FALSE)
```

### 9. Neural Network

```
set.seed(123)

# Load required libraries
library(neuralnet)
library(caret)
library(pROC)
library(ggplot2)

# Prepare the data
Test_Train <- trainData_part1
Test_Test <- trainData_part2
Combined_Test <- combined_testData

# Convert Senescence_Status variable to binary {0, 1}
Test_Train$Senescence_Status <- ifelse(Test_Train$Senescence_Status == "Sen", 1, 0)
Test_Test$Senescence_Status <- ifelse(Test_Test$Senescence_Status == "Sen", 1, 0)
Combined_Test$Senescence_Status <- ifelse(Combined_Test$Senescence_Status == "Sen", 1, 0)

# Define formula for the neural network
formula <- as.formula("Senescence_Status ~ .")

# Train the neural network using trainData_part1
nn_model <- neuralnet(
  formula,
  data = Test_Train,
  hidden = c(20, 10, 5, 2),  # Define the number of neurons in hidden layers
  linear.output = FALSE  # Use sigmoid activation function for binary classification
)

# Prepare meta-model training data
x_train_part2 <- Test_Test[, colnames(Test_Train), drop = FALSE]

# Make predictions on Test_Test using the NN model
predicted_probabilities_part2 <- compute(nn_model, as.matrix(x_train_part2[, -which(names(x_train_part2) == "Senescence_Status")]))$net.result
predicted_classes_part2 <- ifelse(predicted_probabilities_part2 > 0.5, 1, 0)

# Create and save the meta-model data frame for Test_Test
nn_meta_model_data_train <- data.frame(
  nn_Prob = as.vector(predicted_probabilities_part2),
  Senescence_Status = as.factor(Test_Test$Senescence_Status),
  Predicted_Status = factor(predicted_classes_part2, levels = c(0, 1), labels = c("NonSen", "Sen"))
)
write.csv(nn_meta_model_data_train, "nn_meta_model_training.csv", row.names = FALSE)

## Now we are going to apply the model to Combined_Test and evaluate

# Make predictions on Combined_Test using the NN model
x_test_combined <- Combined_Test[, colnames(Test_Train), drop = FALSE]
predicted_probabilities_test <- compute(nn_model, as.matrix(x_test_combined[, -which(names(x_test_combined) == "Senescence_Status")]))$net.result
predicted_classes_test <- ifelse(predicted_probabilities_test > 0.5, 1, 0)

# Create and save the meta-model data frame for Combined_Test
nn_meta_model_data_test <- data.frame(
  nn_Prob = as.vector(predicted_probabilities_test),
  Senescence_Status = as.factor(Combined_Test$Senescence_Status),
  Predicted_Status = factor(predicted_classes_test, levels = c(0, 1), labels = c("NonSen", "Sen"))
)
write.csv(nn_meta_model_data_test, "nn_meta_model_testing.csv", row.names = FALSE)

# Convert predicted_classes and actual classes to factors
Combined_Test$predicted_classes <- factor(predicted_classes_test, levels = c(0, 1), labels = c("NonSen", "Sen"))
Combined_Test$Senescence_Status <- factor(Combined_Test$Senescence_Status, levels = c(0, 1), labels = c("NonSen", "Sen"))

# Create confusion matrix for NN on Combined_Test
nn_confusion_mtx_test <- confusionMatrix(data = Combined_Test$predicted_classes, reference = Combined_Test$Senescence_Status)
print(nn_confusion_mtx_test)
```

```
## Confusion Matrix and Statistics
## 
##           Reference
## Prediction NonSen   Sen
##     NonSen  19379     8
##     Sen      1347    96
##                                           
##                Accuracy : 0.9349          
##                  95% CI : (0.9315, 0.9383)
##     No Information Rate : 0.995           
##     P-Value [Acc > NIR] : 1               
##                                           
##                   Kappa : 0.1159          
##                                           
##  Mcnemar's Test P-Value : <2e-16          
##                                           
##             Sensitivity : 0.93501         
##             Specificity : 0.92308         
##          Pos Pred Value : 0.99959         
##          Neg Pred Value : 0.06653         
##              Prevalence : 0.99501         
##          Detection Rate : 0.93034         
##    Detection Prevalence : 0.93072         
##       Balanced Accuracy : 0.92904         
##                                           
##        'Positive' Class : NonSen          
##
```

```
# Calculate precision, recall, F1-score, specificity, FPR
precision_nn <- nn_confusion_mtx_test$byClass["Pos Pred Value"]
recall_nn <- nn_confusion_mtx_test$byClass["Sensitivity"]
f1_score_nn <- nn_confusion_mtx_test$byClass["F1"]
specificity_nn <- nn_confusion_mtx_test$byClass["Neg Pred Value"]
fpr_nn <- 1 - specificity_nn

# Compute ROC curve and AUC for NN
roc_curve_nn <- roc(Combined_Test$Senescence_Status, predicted_probabilities_test)
```

```
## Setting levels: control = NonSen, case = Sen
```

```
## Warning in roc.default(Combined_Test$Senescence_Status,
## predicted_probabilities_test): Deprecated use a matrix as predictor. Unexpected
## results may be produced, please pass a numeric vector.
```

```
## Setting direction: controls < cases
```

```
auc_roc_nn <- auc(roc_curve_nn)

# Calculate Cohen's Kappa
kappa_nn <- nn_confusion_mtx_test$overall["Kappa"]

# Ensure the 'ROCs' directory exists
if (!dir.exists("ROCs")) {
  dir.create("ROCs")
}

# Create the ROC plot for NN
roc_plot_nn <- ggroc(roc_curve_nn, legacy.axes = TRUE, alpha = 0.5, colour = "blue", linetype = 1, size = 1) +
  labs(title = "ROC Curve on Combined Test Data (Neural Network)", x = "False Positive Rate", y = "True Positive Rate") +
  geom_segment(aes(x = 0, xend = 1, y = 0, yend = 1), color = "black", linetype = "dashed")

# Save the ROC plot
ggsave("ROCs/roc_curve_nn.png", plot = roc_plot_nn, width = 8, height = 6)

# Store metrics and confusion matrix breakdown for Combined_Test in a data frame
NN_metrics_df_test <- data.frame(
  Model = "Neural Network",
  Accuracy = nn_confusion_mtx_test$overall["Accuracy"],
  Precision = precision_nn,
  Recall = recall_nn,
  F1_Score = f1_score_nn,
  AUC = auc_roc_nn,
  Neg_Pred_Value = specificity_nn,
  FPR = fpr_nn,
  Kappa = kappa_nn,
  TP = nn_confusion_mtx_test$table["Sen", "Sen"],
  TN = nn_confusion_mtx_test$table["NonSen", "NonSen"],
  FN = nn_confusion_mtx_test$table["NonSen", "Sen"],
  FP = nn_confusion_mtx_test$table["Sen", "NonSen"]
)

# Save the metrics for Combined_Test to a CSV file
write.csv(NN_metrics_df_test, "neural_network_metrics.csv", row.names = FALSE)

# Custom plot function for neural network
custom_plot_nn <- function(nn_model, plot_title = "Neural Network") {
  plot(nn_model, rep = "best",
       show.weights = FALSE, 
       dimension = c(2000, 800), 
       radius = 3,  # Adjust the radius of the nodes
       fontsize = 18,  # Adjust the font size
       arrow.length = 0.1,  # Adjust the length of the arrows
       information = FALSE,
       main = plot_title)
}

# Plot and save the neural network with custom function
plot_name <- "ROCs/neural_network_plot.png"
png(plot_name, width = 2000, height = 1000)
custom_plot_nn(nn_model)
dev.off()
```

```
## quartz_off_screen 
##                 2
```

```
print(roc_plot_nn)
```

### 10. Meta Model

```
set.seed(123)

# Load required libraries
library(glmnet)
library(caret)
library(pROC)
library(ggplot2)

# Combine model predictions and response variable for training
meta_model_data_train <- as.data.frame(cbind(
  lasso_meta_model_data_training[,1],
  svm_meta_model_data_training[,1],
  en_meta_model_data_training[,1],
  rf_meta_model_data_training[,1],
  mda_meta_model_training[,1],
  logistic_meta_model_data_training[,1],
  nn_meta_model_data_train[,1:2]
))

colnames(meta_model_data_train) <- c("lasso", "svm", "en", "rf", "mda", "logistic", "nn", "Senescence_Status")

# Extract the predictors and response for training
train_predictors <- meta_model_data_train[, -ncol(meta_model_data_train)]
train_response <- as.factor(meta_model_data_train$Senescence_Status)

# Fit the Lasso model on the training meta-model data
lasso_model <- cv.glmnet(
  x = as.matrix(train_predictors), 
  y = as.numeric(train_response) - 1,  # glmnet requires response to be zero-indexed
  family = "binomial", 
  alpha = 1
)

# Get the best lambda
best_lambda <- lasso_model$lambda.min

# Combine model predictions and response variable for testing
meta_model_data_test <- as.data.frame(cbind(
  lasso_meta_model_data_testing[,1],
  svm_meta_model_data_testing[,1],
  en_meta_model_data_testing[,1],
  rf_meta_model_data_testing[,1],
  mda_meta_model_testing[,1],
  logistic_meta_model_data_testing[,1],
  nn_meta_model_data_test[,1:2]
))

colnames(meta_model_data_test) <- c("lasso", "svm", "en", "rf", "mda", "logistic", "nn", "Senescence_Status")

# Extract the predictors and response for testing
test_predictors <- meta_model_data_test[, -ncol(meta_model_data_test)]
test_response <- as.factor(meta_model_data_test$Senescence_Status)

# Make predictions on the meta-model test data using the trained Lasso model
predicted_probabilities_test <- predict(lasso_model, newx = as.matrix(test_predictors), s = best_lambda, type = "response")
predicted_classes_test <- ifelse(predicted_probabilities_test > 0.5, "1", "0")

# Calculate accuracy for Lasso on the test data
Lasso_accuracy_test <- mean(predicted_classes_test == meta_model_data_test$Senescence_Status)
print(paste("Lasso Accuracy on test data:", Lasso_accuracy_test))
```

```
## [1] "Lasso Accuracy on test data: 0.988190110417667"
```

```
# Prepare the data for confusion matrix
meta_model_data_test$predicted_classes <- factor(predicted_classes_test, levels = c("0", "1"))

# Convert actual classes to factor
meta_model_data_test$Senescence_Status <- factor(meta_model_data_test$Senescence_Status, levels = c("0", "1"))

# Create the confusion matrix for the test data
confusion_mtx_test <- confusionMatrix(data = meta_model_data_test$predicted_classes, reference = meta_model_data_test$Senescence_Status)
print(confusion_mtx_test)
```

```
## Confusion Matrix and Statistics
## 
##           Reference
## Prediction     0     1
##          0 20481     1
##          1   245   103
##                                           
##                Accuracy : 0.9882          
##                  95% CI : (0.9866, 0.9896)
##     No Information Rate : 0.995           
##     P-Value [Acc > NIR] : 1               
##                                           
##                   Kappa : 0.4515          
##                                           
##  Mcnemar's Test P-Value : <2e-16          
##                                           
##             Sensitivity : 0.9882          
##             Specificity : 0.9904          
##          Pos Pred Value : 1.0000          
##          Neg Pred Value : 0.2960          
##              Prevalence : 0.9950          
##          Detection Rate : 0.9832          
##    Detection Prevalence : 0.9833          
##       Balanced Accuracy : 0.9893          
##                                           
##        'Positive' Class : 0               
##
```

```
# Calculate additional metrics for the test data
precision_test <- confusion_mtx_test$byClass["Pos Pred Value"]
recall_test <- confusion_mtx_test$byClass["Sensitivity"]
f1_score_test <- confusion_mtx_test$byClass["F1"]
specificity_test <- confusion_mtx_test$byClass["Neg Pred Value"]
fpr_test <- 1 - specificity_test

# Compute ROC curve and AUC for the test data
roc_curve_test <- roc(meta_model_data_test$Senescence_Status, as.numeric(predicted_probabilities_test))
```

```
## Setting levels: control = 0, case = 1
```

```
## Setting direction: controls < cases
```

```
auc_roc_test <- auc(roc_curve_test)

# Calculate Cohen's Kappa for the test data
kappa_test <- confusion_mtx_test$overall["Kappa"]

# Create the ROC plot for the test data
roc_plot_test <- ggroc(roc_curve_test, legacy.axes = TRUE, alpha = 0.5, colour = "blue", linetype = 1, size = 1) +
  labs(title = "ROC Curve on Test Data (Meta Model)", x = "False Positive Rate", y = "True Positive Rate") +
  geom_segment(aes(x = 0, xend = 1, y = 0, yend = 1), color = "black", linetype = "dashed")

# Save the ROC plot for the test data
ggsave("roc_curve_meta_model_test.png", plot = roc_plot_test, width = 8, height = 6)

# Store metrics and confusion matrix breakdown for the test data in a data frame
Meta_metrics_df_test <- data.frame(
  Model = "Meta Model",
  Accuracy = Lasso_accuracy_test,
  Precision = precision_test,
  Recall = recall_test,
  F1_Score = f1_score_test,
  AUC = auc_roc_test,
  Neg_Pred_Value = specificity_test,
  FPR = fpr_test,
  Kappa = kappa_test,
  TP = confusion_mtx_test$table["1", "1"],
  TN = confusion_mtx_test$table["0", "0"],
  FN = confusion_mtx_test$table["0", "1"],
  FP = confusion_mtx_test$table["1", "0"]
)

# Save the metrics for the test data to a CSV file
write.csv(Meta_metrics_df_test, "Meta_Model_metrics_test.csv", row.names = FALSE)
```

### 11. Assessing Model Metrics

```
# Get the list of CSV files in the working directory
csv_files <- list.files(pattern = "\\.csv$")

# Loop through each CSV file and read it into a data frame
for (file in csv_files) {
  # Extract the name of the data frame from the file name
  df_name <- sub(".csv$", "", file)
  
  # Read the CSV file into a data frame
  assign(df_name, read.csv(file))
}


# Define the desired columns
desired_columns <- c("Model", "Accuracy", "Precision", "Recall", "F1_Score", "AUC", "Neg_Pred_Value", "TP", "TN", "FN", "FP")

# Function to subset data frames to include only desired columns
subset_columns <- function(df, columns) {
  # Check if all desired columns are present in the data frame
  if (all(columns %in% colnames(df))) {
    return(df[, columns, drop = FALSE])
  } else {
    # Return a message if some columns are missing
    warning("Some columns are missing in the data frame")
    return(df) # Return the original data frame if columns are missing
  }
}

# Subset each data frame to keep only the desired columns
logistic_model_metrics <- subset_columns(logistic_model_metrics, desired_columns)
lasso_model_metrics <- subset_columns(lasso_model_metrics, desired_columns)
svm_model_metrics <- subset_columns(svm_model_metrics, desired_columns)
en_model_metrics <- subset_columns(en_model_metrics, desired_columns)
rf_model_metrics <- subset_columns(rf_model_metrics, desired_columns)
mda_model_metrics <- subset_columns(mda_model_metrics, desired_columns)
neural_network_metrics <- subset_columns(neural_network_metrics, desired_columns)
Meta_Model_metrics <- subset_columns(Meta_Model_metrics_test, desired_columns)

# Combine the subsetted data frames into one data frame
All_Metric <- rbind(
  logistic_model_metrics,
  lasso_model_metrics,
  svm_model_metrics,
  en_model_metrics,
  rf_model_metrics,
  mda_model_metrics,
  neural_network_metrics,
  Meta_Model_metrics
)

write.csv(All_Metric, "All_Model_Metrics.csv", row.names = FALSE)


# Define a folder to save the heatmaps
output_folder <- "Heatmaps"
if (!dir.exists(output_folder)) {
  dir.create(output_folder)
}

# Function to save heatmap to file
save_heatmap <- function(matrix_data, file_name, cellnote_data) {
  # Define the file path
  file_path <- file.path(output_folder, file_name)
  
  # Open a graphics device to save the plot
  png(file_path, width = 800, height = 600)
  
  # Plot heatmap
  heatmap.2(matrix_data, Rowv = NA, Colv = NA, dendrogram = "none", trace = "none",
            col = rev(colorRampPalette(c("red","white", "turquoise"))(256)), scale = "column", key = TRUE, keysize = 1.5,
            key.title = NA, density.info = "none", cexRow = 0.8, cexCol = 0.8,
            margins = c(8, 8), cellnote = cellnote_data, notecol = "black")
  
  # Close the graphics device
  dev.off()
}

# First Heatmap - Metrics in Percentages
HM_Met <- All_Metric[, -c(1,8:11)]
rownames(HM_Met) <- All_Metric[, 1]
rownames(HM_Met) <- gsub(" \\(Combined Test Data\\)", "", rownames(HM_Met))
HM_Met <- as.matrix(HM_Met)
HM_Met_percent <- HM_Met * 100
HM_Met_percent_rounded <- round(HM_Met_percent, 1)
HM_Met_percent_char <- matrix(paste0(HM_Met_percent_rounded, "%"), nrow = nrow(HM_Met_percent_rounded))

# Save the first heatmap
save_heatmap(HM_Met, "Heatmap_Metrics_Percentages.png", HM_Met_percent_char)
```

```
## quartz_off_screen 
##                 2
```

```
# Second Heatmap - Confusion Matrix Metrics
HM_Met <- All_Metric[, -c(1:7)]
rownames(HM_Met) <- All_Metric[, 1]
rownames(HM_Met) <- gsub(" \\(Combined Test Data\\)", "", rownames(HM_Met))
HM_Met <- as.matrix(HM_Met)
HM_Met <- round(HM_Met, 1)

# Save the second heatmap
save_heatmap(HM_Met, "Heatmap_Confusion_Matrix_Metrics.png", HM_Met)
```

```
## quartz_off_screen 
##                 2
```
