## Supplementary material for "SAMP-Score: A morphology-based machine learning classification method for screening pro-senescence compounds in p16 positive cancer cells": HTML Walkthrough Guides for Code: Supplemental Data 1_4_Building-Models_Clusters.html

Building-Models\_Clusters\_Notebook.knit


### 1. Define Ground Truth

```
library("gplots")
```

```
## 
## Attaching package: 'gplots'
```

```
## The following object is masked from 'package:stats':
## 
##     lowess
```

```
library("RColorBrewer")
library("matrixStats")
library("plyr")
```

```
## 
## Attaching package: 'plyr'
```

```
## The following object is masked from 'package:matrixStats':
## 
##     count
```

```
library("dplyr")
```

```
## 
## Attaching package: 'dplyr'
```

```
## The following objects are masked from 'package:plyr':
## 
##     arrange, count, desc, failwith, id, mutate, rename, summarise,
##     summarize
```

```
## The following object is masked from 'package:matrixStats':
## 
##     count
```

```
## The following objects are masked from 'package:stats':
## 
##     filter, lag
```

```
## The following objects are masked from 'package:base':
## 
##     intersect, setdiff, setequal, union
```

```
library("data.table")
```

```
## 
## Attaching package: 'data.table'
```

```
## The following objects are masked from 'package:dplyr':
## 
##     between, first, last
```

```
library("stringr")
library("ggplot2")
library(glmnet)
```

```
## Loading required package: Matrix
```

```
## Loaded glmnet 4.1-8
```

```
library(caret)
```

```
## Loading required package: lattice
```

```
library(pROC)
```

```
## Type 'citation("pROC")' for a citation.
```

```
## 
## Attaching package: 'pROC'
```

```
## The following objects are masked from 'package:stats':
## 
##     cov, smooth, var
```

```
library(e1071)  # Contains functions for SVM
library(randomForest)
```

```
## randomForest 4.7-1.1
```

```
## Type rfNews() to see new features/changes/bug fixes.
```

```
## 
## Attaching package: 'randomForest'
```

```
## The following object is masked from 'package:ggplot2':
## 
##     margin
```

```
## The following object is masked from 'package:dplyr':
## 
##     combine
```

```
library(mda)
```

```
## Loading required package: class
```

```
## Loaded mda 0.5-4
```

```
library(neuralnet)
```

```
## 
## Attaching package: 'neuralnet'
```

```
## The following object is masked from 'package:dplyr':
## 
##     compute
```

```
# Upload your file 
all<- read.csv("All_Data_Clusters_Consensus.csv")
all <- all[,-c(1)]
```

### 2. Data Partitioning

```
# Load required libraries
library(ROSE)  # For undersampling and oversampling
```

```
## Loaded ROSE 0.0-4
```

```
library(caret)  # For creating data partitions
library(dplyr)  # For data manipulation

#Tidy Variables
All_Tidy <- all[,-c(1:5)]

# Step 1: Split the data into train and test sets (80-20 split)
set.seed(123)  # For reproducibility
trainIndex <- createDataPartition(All_Tidy$Senescence_Status, p = 0.8, list = FALSE)
trainData <- All_Tidy[trainIndex,]
testData <- All_Tidy[-trainIndex,]

# Add a unique identifier to the training data
trainData <- trainData %>%
  ungroup() %>%
  mutate(id = row_number())

# Checking the distribution of the target variable in train and test sets
cat("Distribution in Training Set:\n")
```

```
## Distribution in Training Set:
```

```
print(table(trainData$Senescence_Status))
```

```
## 
## NonSen    Sen 
##  13336   3992
```

```
cat("Distribution in Test Set:\n")
```

```
## Distribution in Test Set:
```

```
print(table(testData$Senescence_Status))
```

```
## 
## NonSen    Sen 
##   3333    997
```

```
# Step 2: Perform undersampling to balance the classes
set.seed(123)
undersample_result <- ovun.sample(Senescence_Status ~ ., data = trainData, method = "under", p = 0.5)$data

# Checking the distribution after undersampling
cat("Distribution after Undersampling:\n")
```

```
## Distribution after Undersampling:
```

```
print(table(undersample_result$Senescence_Status))
```

```
## 
## NonSen    Sen 
##   3952   3992
```

```
# Step 3: Track and Extract Removed Observations
# Identify removed observations
# Removed IDs are those in trainData but not in undersample_result
removed_ids <- setdiff(trainData$id, undersample_result$id)

# Extract removed observations from the original training data
recycled_data <- trainData %>% filter(id %in% removed_ids)

# Remove the unique identifier column from all datasets
trainData <- trainData %>% select(-id)
undersample_result <- undersample_result %>% select(-id)
recycled_data <- recycled_data %>% select(-id)

# Combine recycled data with the test data
combined_testData <- bind_rows(testData, recycled_data)

# Checking the distribution in the combined test set
cat("Distribution in Combined Test Set:\n")
```

```
## Distribution in Combined Test Set:
```

```
print(table(combined_testData$Senescence_Status))
```

```
## 
## NonSen    Sen 
##  12717    997
```

```
# Split the Undersampled Training Data into Training and Validation
set.seed(123)
trainIndex_under <- createDataPartition(undersample_result$Senescence_Status, p = 0.5, list = FALSE)
trainData_part1 <- undersample_result[trainIndex_under,]
trainData_part2 <- undersample_result[-trainIndex_under,]

# Checking the distribution of the target variable in each part
cat("Distribution in Part 1 of Undersampled Data:\n")
```

```
## Distribution in Part 1 of Undersampled Data:
```

```
print(table(trainData_part1$Senescence_Status))
```

```
## 
## NonSen    Sen 
##   1976   1996
```

```
cat("Distribution in Part 2 of Undersampled Data:\n")
```

```
## Distribution in Part 2 of Undersampled Data:
```

```
print(table(trainData_part2$Senescence_Status))
```

```
## 
## NonSen    Sen 
##   1976   1996
```

### 3. Lasso Regularisation

```
# Load required libraries
library(glmnet)
library(caret)
library(pROC)
library(ggplot2)

# Ensure Senescence_Status is a factor (for classification)
trainData_part1$Senescence_Status <- as.factor(trainData_part1$Senescence_Status)
trainData_part2$Senescence_Status <- as.factor(trainData_part2$Senescence_Status)
combined_testData$Senescence_Status <- as.factor(combined_testData$Senescence_Status)

# Prepare training and test data for Lasso
x_train <- as.matrix(trainData_part1[, -which(names(trainData_part1) == "Senescence_Status")])
y_train <- trainData_part1$Senescence_Status
x_test <- as.matrix(combined_testData[, -which(names(combined_testData) == "Senescence_Status")])
y_test <- combined_testData$Senescence_Status

# Fit the Lasso model on trainData_part1
set.seed(123)
lasso_model <- glmnet(x_train, y_train, family = "binomial", alpha = 1)

# Use cross-validation to determine the best lambda
cv_model <- cv.glmnet(x_train, y_train, family = "binomial", alpha = 1)
```

```
## Warning: from glmnet C++ code (error code -48); Convergence for 48th lambda
## value not reached after maxit=100000 iterations; solutions for larger lambdas
## returned
```

```
## Warning: from glmnet C++ code (error code -50); Convergence for 50th lambda
## value not reached after maxit=100000 iterations; solutions for larger lambdas
## returned
```

```
## Confusion Matrix and Statistics
## 
##           Reference
## Prediction NonSen   Sen
##     NonSen  11707    57
##     Sen      1010   940
##                                           
##                Accuracy : 0.9222          
##                  95% CI : (0.9176, 0.9266)
##     No Information Rate : 0.9273          
##     P-Value [Acc > NIR] : 0.9892          
##                                           
##                   Kappa : 0.5994          
##                                           
##  Mcnemar's Test P-Value : <2e-16          
##                                           
##             Sensitivity : 0.9206          
##             Specificity : 0.9428          
##          Pos Pred Value : 0.9952          
##          Neg Pred Value : 0.4821          
##              Prevalence : 0.9273          
##          Detection Rate : 0.8537          
##    Detection Prevalence : 0.8578          
##       Balanced Accuracy : 0.9317          
##                                           
##        'Positive' Class : NonSen          
##
```

```
## Confusion Matrix and Statistics
## 
##           Reference
## Prediction NonSen   Sen
##     NonSen  11711    58
##     Sen      1006   939
##                                           
##                Accuracy : 0.9224          
##                  95% CI : (0.9178, 0.9268)
##     No Information Rate : 0.9273          
##     P-Value [Acc > NIR] : 0.9862          
##                                           
##                   Kappa : 0.5999          
##                                           
##  Mcnemar's Test P-Value : <2e-16          
##                                           
##             Sensitivity : 0.9209          
##             Specificity : 0.9418          
##          Pos Pred Value : 0.9951          
##          Neg Pred Value : 0.4828          
##              Prevalence : 0.9273          
##          Detection Rate : 0.8539          
##    Detection Prevalence : 0.8582          
##       Balanced Accuracy : 0.9314          
##                                           
##        'Positive' Class : NonSen          
##
```

```
## Confusion Matrix and Statistics
## 
##           Reference
## Prediction NonSen   Sen
##     NonSen  11714    59
##     Sen      1003   938
##                                         
##                Accuracy : 0.9226        
##                  95% CI : (0.918, 0.927)
##     No Information Rate : 0.9273        
##     P-Value [Acc > NIR] : 0.9837        
##                                         
##                   Kappa : 0.6001        
##                                         
##  Mcnemar's Test P-Value : <2e-16        
##                                         
##             Sensitivity : 0.9211        
##             Specificity : 0.9408        
##          Pos Pred Value : 0.9950        
##          Neg Pred Value : 0.4833        
##              Prevalence : 0.9273        
##          Detection Rate : 0.8542        
##    Detection Prevalence : 0.8585        
##       Balanced Accuracy : 0.9310        
##                                         
##        'Positive' Class : NonSen        
##
```

```
## Confusion Matrix and Statistics
## 
##           Reference
## Prediction NonSen   Sen
##     NonSen  11840    31
##     Sen       877   966
##                                           
##                Accuracy : 0.9338          
##                  95% CI : (0.9295, 0.9379)
##     No Information Rate : 0.9273          
##     P-Value [Acc > NIR] : 0.001604        
##                                           
##                   Kappa : 0.647           
##                                           
##  Mcnemar's Test P-Value : < 2.2e-16       
##                                           
##             Sensitivity : 0.9310          
##             Specificity : 0.9689          
##          Pos Pred Value : 0.9974          
##          Neg Pred Value : 0.5241          
##              Prevalence : 0.9273          
##          Detection Rate : 0.8634          
##    Detection Prevalence : 0.8656          
##       Balanced Accuracy : 0.9500          
##                                           
##        'Positive' Class : NonSen          
##
```

```
## Confusion Matrix and Statistics
## 
##           Reference
## Prediction NonSen   Sen
##     NonSen  11961    34
##     Sen       756   963
##                                           
##                Accuracy : 0.9424          
##                  95% CI : (0.9384, 0.9462)
##     No Information Rate : 0.9273          
##     P-Value [Acc > NIR] : 1.076e-12       
##                                           
##                   Kappa : 0.6797          
##                                           
##  Mcnemar's Test P-Value : < 2.2e-16       
##                                           
##             Sensitivity : 0.9406          
##             Specificity : 0.9659          
##          Pos Pred Value : 0.9972          
##          Neg Pred Value : 0.5602          
##              Prevalence : 0.9273          
##          Detection Rate : 0.8722          
##    Detection Prevalence : 0.8747          
##       Balanced Accuracy : 0.9532          
##                                           
##        'Positive' Class : NonSen          
##
```

```
## [1] "MDA Accuracy on combined_testData: 0.923581741286277"
```

```
# Assign predicted classes to combined_testData
combined_testData_pred <- combined_testData
combined_testData_pred$predicted_classes <- predicted_classes_test_mda

# Convert predicted_classes and actual classes to factors
combined_testData_pred$predicted_classes <- factor(predicted_classes_test_mda, levels = c("NonSen", "Sen"))
combined_testData_pred$Senescence_Status <- factor(combined_testData_pred$Senescence_Status)

# Create confusion matrix for MDA on combined_testData
mda_confusion_mtx_test <- confusionMatrix(data = combined_testData_pred$predicted_classes, reference = combined_testData_pred$Senescence_Status)
print(mda_confusion_mtx_test)
```

```
## Confusion Matrix and Statistics
## 
##           Reference
## Prediction NonSen   Sen
##     NonSen  11724    55
##     Sen       993   942
##                                         
##                Accuracy : 0.9236        
##                  95% CI : (0.919, 0.928)
##     No Information Rate : 0.9273        
##     P-Value [Acc > NIR] : 0.954         
##                                         
##                   Kappa : 0.6046        
##                                         
##  Mcnemar's Test P-Value : <2e-16        
##                                         
##             Sensitivity : 0.9219        
##             Specificity : 0.9448        
##          Pos Pred Value : 0.9953        
##          Neg Pred Value : 0.4868        
##              Prevalence : 0.9273        
##          Detection Rate : 0.8549        
##    Detection Prevalence : 0.8589        
##       Balanced Accuracy : 0.9334        
##                                         
##        'Positive' Class : NonSen        
##
```

```
## Confusion Matrix and Statistics
## 
##           Reference
## Prediction NonSen   Sen
##     NonSen  11760    54
##     Sen       957   943
##                                           
##                Accuracy : 0.9263          
##                  95% CI : (0.9218, 0.9306)
##     No Information Rate : 0.9273          
##     P-Value [Acc > NIR] : 0.6846          
##                                           
##                   Kappa : 0.6142          
##                                           
##  Mcnemar's Test P-Value : <2e-16          
##                                           
##             Sensitivity : 0.9247          
##             Specificity : 0.9458          
##          Pos Pred Value : 0.9954          
##          Neg Pred Value : 0.4963          
##              Prevalence : 0.9273          
##          Detection Rate : 0.8575          
##    Detection Prevalence : 0.8615          
##       Balanced Accuracy : 0.9353          
##                                           
##        'Positive' Class : NonSen          
##
```

```
# Calculate precision, recall, F1-score, specificity, FPR
precision_nn <- nn_confusion_mtx_test$byClass["Pos Pred Value"]
recall_nn <- nn_confusion_mtx_test$byClass["Sensitivity"]
f1_score_nn <- nn_confusion_mtx_test$byClass["F1"]
specificity_nn <- nn_confusion_mtx_test$byClass["Neg Pred Value"]
fpr_nn <- 1 - specificity_nn

# Compute ROC curve and AUC for NN
roc_curve_nn <- roc(Combined_Test$Senescence_Status, predicted_probabilities_test)
```

```
## Setting levels: control = NonSen, case = Sen
```

```
## Warning in roc.default(Combined_Test$Senescence_Status,
## predicted_probabilities_test): Deprecated use a matrix as predictor. Unexpected
## results may be produced, please pass a numeric vector.
```

```
## Setting direction: controls < cases
```

```
auc_roc_nn <- auc(roc_curve_nn)

# Calculate Cohen's Kappa
kappa_nn <- nn_confusion_mtx_test$overall["Kappa"]

# Ensure the 'ROCs' directory exists
if (!dir.exists("ROCs")) {
  dir.create("ROCs")
}

# Create the ROC plot for NN
roc_plot_nn <- ggroc(roc_curve_nn, legacy.axes = TRUE, alpha = 0.5, colour = "blue", linetype = 1, size = 1) +
  labs(title = "ROC Curve on Combined Test Data (Neural Network)", x = "False Positive Rate", y = "True Positive Rate") +
  geom_segment(aes(x = 0, xend = 1, y = 0, yend = 1), color = "black", linetype = "dashed")

# Save the ROC plot
ggsave("ROCs/roc_curve_nn.png", plot = roc_plot_nn, width = 8, height = 6)

# Store metrics and confusion matrix breakdown for Combined_Test in a data frame
NN_metrics_df_test <- data.frame(
  Model = "Neural Network",
  Accuracy = nn_confusion_mtx_test$overall["Accuracy"],
  Precision = precision_nn,
  Recall = recall_nn,
  F1_Score = f1_score_nn,
  AUC = auc_roc_nn,
  Neg_Pred_Value = specificity_nn,
  FPR = fpr_nn,
  Kappa = kappa_nn,
  TP = nn_confusion_mtx_test$table["Sen", "Sen"],
  TN = nn_confusion_mtx_test$table["NonSen", "NonSen"],
  FN = nn_confusion_mtx_test$table["NonSen", "Sen"],
  FP = nn_confusion_mtx_test$table["Sen", "NonSen"]
)

# Save the metrics for Combined_Test to a CSV file
write.csv(NN_metrics_df_test, "neural_network_metrics.csv", row.names = FALSE)

# Custom plot function for neural network
custom_plot_nn <- function(nn_model, plot_title = "Neural Network") {
  plot(nn_model, rep = "best",
       show.weights = TRUE, 
       dimension = c(2000, 800), 
       radius = 3,  # Adjust the radius of the nodes
       fontsize = 18,  # Adjust the font size
       arrow.length = 0.1,  # Adjust the length of the arrows
       information = FALSE,
       main = plot_title)
}

# Plot and save the neural network with custom function
plot_name <- "ROCs/neural_network_plot.png"
png(plot_name, width = 2000, height = 1000)
custom_plot_nn(nn_model)
dev.off()
```

```
## quartz_off_screen 
##                 2
```

### 10. Meta Model

```
set.seed(123)

# Load required libraries
library(glmnet)
library(caret)
library(pROC)
library(ggplot2)

# Combine model predictions and response variable for training
meta_model_data_train <- as.data.frame(cbind(
  lasso_meta_model_data_training[,1],
  svm_meta_model_data_training[,1],
  en_meta_model_data_training[,1],
  rf_meta_model_data_training[,1],
  mda_meta_model_training[,1],
  logistic_meta_model_data_training[,1],
  nn_meta_model_data_train[,1:2]
))

colnames(meta_model_data_train) <- c("lasso", "svm", "en", "rf", "mda", "logistic", "nn", "Senescence_Status")

# Extract the predictors and response for training
train_predictors <- meta_model_data_train[, -ncol(meta_model_data_train)]
train_response <- as.factor(meta_model_data_train$Senescence_Status)

# Fit the Lasso model on the training meta-model data
lasso_model <- cv.glmnet(
  x = as.matrix(train_predictors), 
  y = as.numeric(train_response) - 1,  # glmnet requires response to be zero-indexed
  family = "binomial", 
  alpha = 1
)

# Get the best lambda
best_lambda <- lasso_model$lambda.min

# Combine model predictions and response variable for testing
meta_model_data_test <- as.data.frame(cbind(
  lasso_meta_model_data_testing[,1],
  svm_meta_model_data_testing[,1],
  en_meta_model_data_testing[,1],
  rf_meta_model_data_testing[,1],
  mda_meta_model_testing[,1],
  logistic_meta_model_data_testing[,1],
  nn_meta_model_data_test[,1:2]
))

colnames(meta_model_data_test) <- c("lasso", "svm", "en", "rf", "mda", "logistic", "nn", "Senescence_Status")

# Extract the predictors and response for testing
test_predictors <- meta_model_data_test[, -ncol(meta_model_data_test)]
test_response <- as.factor(meta_model_data_test$Senescence_Status)

# Make predictions on the meta-model test data using the trained Lasso model
predicted_probabilities_test <- predict(lasso_model, newx = as.matrix(test_predictors), s = best_lambda, type = "response")
predicted_classes_test <- ifelse(predicted_probabilities_test > 0.50, "1", "0")

Lasso_accuracy_test <- mean(predicted_classes_test == meta_model_data_test$Senescence_Status)
print(paste("Lasso Accuracy on test data:", Lasso_accuracy_test))
```

```
## [1] "Lasso Accuracy on test data: 0.948228088085168"
```

```
# Prepare the data for confusion matrix
meta_model_data_test$predicted_classes <- factor(predicted_classes_test, levels = c("0", "1"))

# Convert actual classes to factor
meta_model_data_test$Senescence_Status <- factor(meta_model_data_test$Senescence_Status, levels = c("0", "1"))

# Create the confusion matrix for the test data
confusion_mtx_test <- confusionMatrix(data = meta_model_data_test$predicted_classes, reference = meta_model_data_test$Senescence_Status)
print(confusion_mtx_test)
```

```
## Confusion Matrix and Statistics
## 
##           Reference
## Prediction     0     1
##          0 12037    30
##          1   680   967
##                                           
##                Accuracy : 0.9482          
##                  95% CI : (0.9444, 0.9519)
##     No Information Rate : 0.9273          
##     P-Value [Acc > NIR] : < 2.2e-16       
##                                           
##                   Kappa : 0.7047          
##                                           
##  Mcnemar's Test P-Value : < 2.2e-16       
##                                           
##             Sensitivity : 0.9465          
##             Specificity : 0.9699          
##          Pos Pred Value : 0.9975          
##          Neg Pred Value : 0.5871          
##              Prevalence : 0.9273          
##          Detection Rate : 0.8777          
##    Detection Prevalence : 0.8799          
##       Balanced Accuracy : 0.9582          
##                                           
##        'Positive' Class : 0               
##
```

```
# Calculate additional metrics for the test data - Note Caret will mistakenly swap Pos Pred Value and Neg Pred Values - these are corrected in manuscript figure
precision_test <- confusion_mtx_test$byClass["Pos Pred Value"]
recall_test <- confusion_mtx_test$byClass["Sensitivity"]
f1_score_test <- confusion_mtx_test$byClass["F1"]
specificity_test <- confusion_mtx_test$byClass["Neg Pred Value"]
fpr_test <- 1 - specificity_test

# Compute ROC curve and AUC for the test data
roc_curve_test <- roc(meta_model_data_test$Senescence_Status, as.numeric(predicted_probabilities_test))
```

```
## quartz_off_screen 
##                 2
```
