## Supplementary material for "SAMP-Score: A morphology-based machine learning classification method for screening pro-senescence compounds in p16 positive cancer cells": HTML Walkthrough Guides for Code: Supplemental Data 1_5_Wrangle_DDU_Screen_1.html

5\_Wrangle\_DDU\_Screen\_1.knit


### 1\_Import Data.R

```
# Load the libraries
library("gplots")
library("RColorBrewer")
library("dplyr")
library("matrixStats")

# Batch 1 P585-P599
PL585 <- read.csv("DDU0517B1P585.csv", fileEncoding = 'UTF-8-BOM')
PL586 <- read.csv("DDU0517B1P586.csv", fileEncoding = 'UTF-8-BOM')
PL587 <- read.csv("DDU0517B1P587.csv", fileEncoding = 'UTF-8-BOM')
PL588 <- read.csv("DDU0517B1P588.csv", fileEncoding = 'UTF-8-BOM')
PL589 <- read.csv("DDU0517B1P589.csv", fileEncoding = 'UTF-8-BOM')
PL590 <- read.csv("DDU0517B1P590.csv", fileEncoding = 'UTF-8-BOM')
PL591 <- read.csv("DDU0517B1P591.csv", fileEncoding = 'UTF-8-BOM')
PL592 <- read.csv("DDU0517B1P592.csv", fileEncoding = 'UTF-8-BOM')
PL593 <- read.csv("DDU0517B1P593.csv", fileEncoding = 'UTF-8-BOM')
PL594 <- read.csv("DDU0517B1P594.csv", fileEncoding = 'UTF-8-BOM')
PL595 <- read.csv("DDU0517B1P595.csv", fileEncoding = 'UTF-8-BOM')
PL596 <- read.csv("DDU0517B1P596.csv", fileEncoding = 'UTF-8-BOM')
PL597 <- read.csv("DDU0517B1P597.csv", fileEncoding = 'UTF-8-BOM')
PL598 <- read.csv("DDU0517B1P598.csv", fileEncoding = 'UTF-8-BOM')
PL599 <- read.csv("DDU0517B1P599.csv", fileEncoding = 'UTF-8-BOM')

# Batch 2 P601-P614
PL601 <- read.csv("DDU0517B2P601.csv", fileEncoding = 'UTF-8-BOM')
PL602 <- read.csv("DDU0517B2P602.csv", fileEncoding = 'UTF-8-BOM')
PL603 <- read.csv("DDU0517B2P603.csv", fileEncoding = 'UTF-8-BOM')
PL604 <- read.csv("DDU0517B2P604.csv", fileEncoding = 'UTF-8-BOM')
PL605 <- read.csv("DDU0517B2P605.csv", fileEncoding = 'UTF-8-BOM')
PL606 <- read.csv("DDU0517B2P606.csv", fileEncoding = 'UTF-8-BOM')
PL607 <- read.csv("DDU0517B2P607.csv", fileEncoding = 'UTF-8-BOM')
PL608 <- read.csv("DDU0517B2P608.csv", fileEncoding = 'UTF-8-BOM')
PL609 <- read.csv("DDU0517B2P609.csv", fileEncoding = 'UTF-8-BOM')
PL610 <- read.csv("DDU0517B2P610.csv", fileEncoding = 'UTF-8-BOM')
PL611 <- read.csv("DDU0517B2P611.csv", fileEncoding = 'UTF-8-BOM')
PL612 <- read.csv("DDU0517B2P612.csv", fileEncoding = 'UTF-8-BOM')
PL613 <- read.csv("DDU0517B2P613.csv", fileEncoding = 'UTF-8-BOM')
PL614 <- read.csv("DDU1517B2P614.csv", fileEncoding = 'UTF-8-BOM')

# Batch 3 P615-P629
PL615 <- read.csv("DDU1517B3P615.csv", fileEncoding = 'UTF-8-BOM')
PL616 <- read.csv("DDU1517B3P616.csv", fileEncoding = 'UTF-8-BOM')
PL617 <- read.csv("DDU1517B3P617.csv", fileEncoding = 'UTF-8-BOM')
PL618 <- read.csv("DDU1517B3P618.csv", fileEncoding = 'UTF-8-BOM')
PL619 <- read.csv("DDU1517B3P619.csv", fileEncoding = 'UTF-8-BOM')
PL620 <- read.csv("DDU1517B3P620.csv", fileEncoding = 'UTF-8-BOM')
PL621 <- read.csv("DDU1517B3P621.csv", fileEncoding = 'UTF-8-BOM')
PL622 <- read.csv("DDU1517B3P622.csv", fileEncoding = 'UTF-8-BOM')
PL623 <- read.csv("DDU1517B3P623.csv", fileEncoding = 'UTF-8-BOM')
PL624 <- read.csv("DDU1517B3P624.csv", fileEncoding = 'UTF-8-BOM')
PL625 <- read.csv("DDU1517B3P625.csv", fileEncoding = 'UTF-8-BOM')
PL626 <- read.csv("DDU1517B3P626.csv", fileEncoding = 'UTF-8-BOM')
PL627 <- read.csv("DDU1517B3P627.csv", fileEncoding = 'UTF-8-BOM')
PL628 <- read.csv("DDU1517B3P628.csv", fileEncoding = 'UTF-8-BOM')
PL629 <- read.csv("DDU1517B3P629.csv", fileEncoding = 'UTF-8-BOM')

# Batch 4 P630-P644
PL630 <- read.csv("DDU1517B4P630.csv", fileEncoding = 'UTF-8-BOM')
PL631 <- read.csv("DDU1517B4P631.csv", fileEncoding = 'UTF-8-BOM')
PL632 <- read.csv("DDU1517B4P632.csv", fileEncoding = 'UTF-8-BOM')
PL633 <- read.csv("DDU1517B4P633.csv", fileEncoding = 'UTF-8-BOM')
PL634 <- read.csv("DDU1517B4P634.csv", fileEncoding = 'UTF-8-BOM')
PL635 <- read.csv("DDU1517B4P635.csv", fileEncoding = 'UTF-8-BOM')
PL636 <- read.csv("DDU1517B4P636.csv", fileEncoding = 'UTF-8-BOM')
PL637 <- read.csv("DDU1517B4P637.csv", fileEncoding = 'UTF-8-BOM')
PL638 <- read.csv("DDU1517B4P638.csv", fileEncoding = 'UTF-8-BOM')
PL639 <- read.csv("DDU1517B4P639.csv", fileEncoding = 'UTF-8-BOM')
PL640 <- read.csv("DDU1517B4P640.csv", fileEncoding = 'UTF-8-BOM')
PL641 <- read.csv("DDU1517B4P641.csv", fileEncoding = 'UTF-8-BOM')
PL642 <- read.csv("DDU1517B4P642.csv", fileEncoding = 'UTF-8-BOM')
PL643 <- read.csv("DDU1517B4P643.csv", fileEncoding = 'UTF-8-BOM')
PL644 <- read.csv("DDU1517B4P644.csv", fileEncoding = 'UTF-8-BOM')

# Batch 5 P600
PL600 <- read.csv("DDU1517B5P600P1071.csv", fileEncoding = 'UTF-8-BOM')
```

### 2\_Add Colnames.R

```
library("readxl")

# Import Colnames
colnames<- read.csv("Colnames_Long.csv")
colnames <- colnames(colnames)

# Assign column names to each data frame
colnames(PL585) <- colnames
colnames(PL586) <- colnames
colnames(PL587) <- colnames
colnames(PL588) <- colnames
colnames(PL589) <- colnames
colnames(PL590) <- colnames
colnames(PL591) <- colnames
colnames(PL592) <- colnames
colnames(PL593) <- colnames
colnames(PL594) <- colnames
colnames(PL595) <- colnames
colnames(PL596) <- colnames
colnames(PL597) <- colnames
colnames(PL598) <- colnames
colnames(PL599) <- colnames

colnames(PL601) <- colnames
colnames(PL602) <- colnames
colnames(PL603) <- colnames
colnames(PL604) <- colnames
colnames(PL605) <- colnames
colnames(PL606) <- colnames
colnames(PL607) <- colnames
colnames(PL608) <- colnames
colnames(PL609) <- colnames
colnames(PL610) <- colnames
colnames(PL611) <- colnames
colnames(PL612) <- colnames
colnames(PL613) <- colnames
colnames(PL614) <- colnames

colnames(PL615) <- colnames
colnames(PL616) <- colnames
colnames(PL617) <- colnames
colnames(PL618) <- colnames
colnames(PL619) <- colnames
colnames(PL620) <- colnames
colnames(PL621) <- colnames
colnames(PL622) <- colnames
colnames(PL623) <- colnames
colnames(PL624) <- colnames
colnames(PL625) <- colnames
colnames(PL626) <- colnames
colnames(PL627) <- colnames
colnames(PL628) <- colnames
colnames(PL629) <- colnames

colnames(PL630) <- colnames
colnames(PL631) <- colnames
colnames(PL632) <- colnames
colnames(PL633) <- colnames
colnames(PL634) <- colnames
colnames(PL635) <- colnames
colnames(PL636) <- colnames
colnames(PL637) <- colnames
colnames(PL638) <- colnames
colnames(PL639) <- colnames
colnames(PL640) <- colnames
colnames(PL641) <- colnames
colnames(PL642) <- colnames
colnames(PL643) <- colnames
colnames(PL644) <- colnames

colnames(PL600) <- colnames
```

### 3\_Tidy Measures.R

```
# Column Number for Measures
Measure_Names <- read.csv("Measure_Names.csv", fileEncoding = 'UTF-8-BOM')
Measure_Names <- colnames(Measure_Names)

data_frame_names <- mget(paste0("PL", 585:644))

# Update each dataframe in the list
lapply(names(data_frame_names), function(name) {
  assign(name, value = data_frame_names[[name]] %>%
           select(any_of(Measure_Names)), envir = .GlobalEnv)
})
```

### 4\_Import Plate Maps.R

```
# Read Excel files with the first row included as data
PM1 <- read_excel("DDU_17AP0585_17AP0615_MAP_updated.xlsx")
PM2 <- read_excel("DDU_17AP0586_17AP0616_MAP_updated.xlsx")
PM3 <- read_excel("DDU_17AP0587_17AP0617_MAP_updated.xlsx")
PM4 <- read_excel("DDU_17AP0588_17AP0618_MAP_updated.xlsx")
PM5 <- read_excel("DDU_17AP0589_17AP0619_MAP_updated.xlsx")
PM6 <- read_excel("DDU_17AP0590_17AP0620_MAP_updated.xlsx")
PM7 <- read_excel("DDU_17AP0591_17AP0621_MAP_updated.xlsx")
PM8 <- read_excel("DDU_17AP0592_17AP0622_MAP_updated.xlsx")
PM9 <- read_excel("DDU_17AP0593_17AP0623_MAP_updated.xlsx")
PM10 <- read_excel("DDU_17AP0594_17AP0624_MAP_updated.xlsx")
PM11 <- read_excel("DDU_17AP0595_17AP0625_MAP_updated.xlsx")
PM12 <- read_excel("DDU_17AP0596_17AP0626_MAP_updated.xlsx")
PM13 <- read_excel("DDU_17AP0597_17AP0627_MAP_updated.xlsx")
PM14 <- read_excel("DDU_17AP0598_17AP0628_MAP_updated.xlsx")
PM15 <- read_excel("DDU_17AP0599_17AP0629_MAP_updated.xlsx")

#Only Keep whats needed
PM1 <- PM1[, c(1, 6)]
PM2 <- PM2[, c(1, 6)]
PM3 <- PM3[, c(1, 6)]
PM4 <- PM4[, c(1, 6)]
PM5 <- PM5[, c(1, 6)]
PM6 <- PM6[, c(1, 6)]
PM7 <- PM7[, c(1, 6)]
PM8 <- PM8[, c(1, 6)]
PM9 <- PM9[, c(1, 6)]
PM10 <- PM10[, c(1, 6)]
PM11 <- PM11[, c(1, 6)]
PM12 <- PM12[, c(1, 6)]
PM13 <- PM13[, c(1, 6)]
PM14 <- PM14[, c(1, 6)]
PM15 <- PM15[, c(1, 6)]

# Import Second Set

PM16 <- read_excel("DDU_17AP0600_17AP0630_MAP_updated.xlsx", col_names = FALSE)
PM17 <- read_excel("DDU_17AP0601_17AP0631_MAP_updated.xlsx", col_names = FALSE)
PM18 <- read_excel("DDU_17AP0602_17AP0632_MAP_updated.xlsx", col_names = FALSE)
PM19 <- read_excel("DDU_17AP0603_17AP0633_MAP_updated.xlsx", col_names = FALSE)
PM20 <- read_excel("DDU_17AP0604_17AP0634_MAP_updated.xlsx", col_names = FALSE)
PM21 <- read_excel("DDU_17AP0605_17AP0635_MAP_updated.xlsx", col_names = FALSE)
PM22 <- read_excel("DDU_17AP0606_17AP0636_MAP_updated.xlsx", col_names = FALSE)
PM23 <- read_excel("DDU_17AP0607_17AP0637_MAP_updated.xlsx", col_names = FALSE)
PM24 <- read_excel("DDU_17AP0608_17AP0638_MAP_updated.xlsx", col_names = FALSE)
PM25 <- read_excel("DDU_17AP0609_17AP0639_MAP_updated.xlsx", col_names = FALSE)
PM26 <- read_excel("DDU_17AP0610_17AP0640_MAP_updated.xlsx", col_names = FALSE)
PM27 <- read_excel("DDU_17AP0611_17AP0641_MAP_updated.xlsx", col_names = FALSE)
PM28 <- read_excel("DDU_17AP0612_17AP0642_MAP_updated.xlsx", col_names = FALSE)
PM29 <- read_excel("DDU_17AP0613_17AP0643_MAP_updated.xlsx", col_names = FALSE)
PM30 <- read_excel("DDU_17AP0614_17AP0644_MAP_updated.xlsx", col_names = FALSE)

#Keep whats needed

PM16 <- PM16[, c(2, 7)]
PM17 <- PM17[, c(2, 7)]
PM18 <- PM18[, c(2, 7)]
PM19 <- PM19[, c(2, 7)]
PM20 <- PM20[, c(2, 7)]
PM21 <- PM21[, c(2, 7)]
PM22 <- PM22[, c(2, 7)]
PM23 <- PM23[, c(2, 7)]
PM24 <- PM24[, c(2, 7)]
PM25 <- PM25[, c(2, 7)]
PM26 <- PM26[, c(2, 7)]
PM27 <- PM27[, c(2, 7)]
PM28 <- PM28[, c(2, 7)]
PM29 <- PM29[, c(2, 7)]
PM30 <- PM30[, c(2, 7)]

# Set column names of the last 15 dataframes to be the same as PM1
colnames(PM16) <- colnames(PM1)
colnames(PM17) <- colnames(PM1)
colnames(PM18) <- colnames(PM1)
colnames(PM19) <- colnames(PM1)
colnames(PM20) <- colnames(PM1)
colnames(PM21) <- colnames(PM1)
colnames(PM22) <- colnames(PM1)
colnames(PM23) <- colnames(PM1)
colnames(PM24) <- colnames(PM1)
colnames(PM25) <- colnames(PM1)
colnames(PM26) <- colnames(PM1)
colnames(PM27) <- colnames(PM1)
colnames(PM28) <- colnames(PM1)
colnames(PM29) <- colnames(PM1)
colnames(PM30) <- colnames(PM1)
```

### 5\_Update Well Labels.R

```
# Create a function to convert the PTODWELLREFERENCE values
convert_PTODWELLREFERENCE <- function(value) {
  letter <- substr(value, 1, 1)  # Extract the first character (letter)
  number <- as.numeric(substr(value, 2, nchar(value)))  # Extract the numbers
  return(paste(letter, "-", number, sep = " "))  # Concatenate letter, "-", and number
}

# Convert PTODWELLREFERENCE column for each dataframe
PM1$PTODWELLREFERENCE <- convert_PTODWELLREFERENCE(PM1$PTODWELLREFERENCE)
PM2$PTODWELLREFERENCE <- convert_PTODWELLREFERENCE(PM2$PTODWELLREFERENCE)
PM3$PTODWELLREFERENCE <- convert_PTODWELLREFERENCE(PM3$PTODWELLREFERENCE)
PM4$PTODWELLREFERENCE <- convert_PTODWELLREFERENCE(PM4$PTODWELLREFERENCE)
PM5$PTODWELLREFERENCE <- convert_PTODWELLREFERENCE(PM5$PTODWELLREFERENCE)
PM6$PTODWELLREFERENCE <- convert_PTODWELLREFERENCE(PM6$PTODWELLREFERENCE)
PM7$PTODWELLREFERENCE <- convert_PTODWELLREFERENCE(PM7$PTODWELLREFERENCE)
PM8$PTODWELLREFERENCE <- convert_PTODWELLREFERENCE(PM8$PTODWELLREFERENCE)
PM9$PTODWELLREFERENCE <- convert_PTODWELLREFERENCE(PM9$PTODWELLREFERENCE)
PM10$PTODWELLREFERENCE <- convert_PTODWELLREFERENCE(PM10$PTODWELLREFERENCE)
PM11$PTODWELLREFERENCE <- convert_PTODWELLREFERENCE(PM11$PTODWELLREFERENCE)
PM12$PTODWELLREFERENCE <- convert_PTODWELLREFERENCE(PM12$PTODWELLREFERENCE)
PM13$PTODWELLREFERENCE <- convert_PTODWELLREFERENCE(PM13$PTODWELLREFERENCE)
PM14$PTODWELLREFERENCE <- convert_PTODWELLREFERENCE(PM14$PTODWELLREFERENCE)
PM15$PTODWELLREFERENCE <- convert_PTODWELLREFERENCE(PM15$PTODWELLREFERENCE)
PM16$PTODWELLREFERENCE <- convert_PTODWELLREFERENCE(PM16$PTODWELLREFERENCE)
PM17$PTODWELLREFERENCE <- convert_PTODWELLREFERENCE(PM17$PTODWELLREFERENCE)
PM18$PTODWELLREFERENCE <- convert_PTODWELLREFERENCE(PM18$PTODWELLREFERENCE)
PM19$PTODWELLREFERENCE <- convert_PTODWELLREFERENCE(PM19$PTODWELLREFERENCE)
PM20$PTODWELLREFERENCE <- convert_PTODWELLREFERENCE(PM20$PTODWELLREFERENCE)
PM21$PTODWELLREFERENCE <- convert_PTODWELLREFERENCE(PM21$PTODWELLREFERENCE)
PM22$PTODWELLREFERENCE <- convert_PTODWELLREFERENCE(PM22$PTODWELLREFERENCE)
PM23$PTODWELLREFERENCE <- convert_PTODWELLREFERENCE(PM23$PTODWELLREFERENCE)
PM24$PTODWELLREFERENCE <- convert_PTODWELLREFERENCE(PM24$PTODWELLREFERENCE)
PM25$PTODWELLREFERENCE <- convert_PTODWELLREFERENCE(PM25$PTODWELLREFERENCE)
PM26$PTODWELLREFERENCE <- convert_PTODWELLREFERENCE(PM26$PTODWELLREFERENCE)
PM27$PTODWELLREFERENCE <- convert_PTODWELLREFERENCE(PM27$PTODWELLREFERENCE)
PM28$PTODWELLREFERENCE <- convert_PTODWELLREFERENCE(PM28$PTODWELLREFERENCE)
PM29$PTODWELLREFERENCE <- convert_PTODWELLREFERENCE(PM29$PTODWELLREFERENCE)
PM30$PTODWELLREFERENCE <- convert_PTODWELLREFERENCE(PM30$PTODWELLREFERENCE)

#Update to Match
colnames(PM1) <- c("Compound", "WELL.LABEL")
colnames(PM2) <- c("Compound", "WELL.LABEL")
colnames(PM3) <- c("Compound", "WELL.LABEL")
colnames(PM4) <- c("Compound", "WELL.LABEL")
colnames(PM5) <- c("Compound", "WELL.LABEL")
colnames(PM6) <- c("Compound", "WELL.LABEL")
colnames(PM7) <- c("Compound", "WELL.LABEL")
colnames(PM8) <- c("Compound", "WELL.LABEL")
colnames(PM9) <- c("Compound", "WELL.LABEL")
colnames(PM10) <- c("Compound", "WELL.LABEL")
colnames(PM11) <- c("Compound", "WELL.LABEL")
colnames(PM12) <- c("Compound", "WELL.LABEL")
colnames(PM13) <- c("Compound", "WELL.LABEL")
colnames(PM14) <- c("Compound", "WELL.LABEL")
colnames(PM15) <- c("Compound", "WELL.LABEL")
colnames(PM16) <- c("Compound", "WELL.LABEL")
colnames(PM17) <- c("Compound", "WELL.LABEL")
colnames(PM18) <- c("Compound", "WELL.LABEL")
colnames(PM19) <- c("Compound", "WELL.LABEL")
colnames(PM20) <- c("Compound", "WELL.LABEL")
colnames(PM21) <- c("Compound", "WELL.LABEL")
colnames(PM22) <- c("Compound", "WELL.LABEL")
colnames(PM23) <- c("Compound", "WELL.LABEL")
colnames(PM24) <- c("Compound", "WELL.LABEL")
colnames(PM25) <- c("Compound", "WELL.LABEL")
colnames(PM26) <- c("Compound", "WELL.LABEL")
colnames(PM27) <- c("Compound", "WELL.LABEL")
colnames(PM28) <- c("Compound", "WELL.LABEL")
colnames(PM29) <- c("Compound", "WELL.LABEL")
colnames(PM30) <- c("Compound", "WELL.LABEL")
```

### 6\_Add Compounds.R

```
# Loop through the range of PL dataframes
for (i in 585:614) {
  # Join PL dataframe with corresponding PM dataframe
  df_name <- paste0("PL", i)
  pm_name <- paste0("PM", (i - 584))
  assign(df_name, merge(get(df_name), get(pm_name), by = "WELL.LABEL", all.x = TRUE))
  
  # Add "DMSO" to Compound if it's "LowControl" or "HighControl"
  df <- get(df_name)
  df$Compound[df$Compound %in% c("LowControl", "HighControl")] <- "DMSO"
  
  # Remove rows with NA values
  df <- na.omit(df)
  
  assign(df_name, df)
}

# Loop through the range of PL dataframes from 615 to 644
for (i in 615:644) {
  # Join PL dataframe with corresponding PM dataframe
  df_name <- paste0("PL", i)
  pm_name <- paste0("PM", 30 - (644 - i))
  assign(df_name, merge(get(df_name), get(pm_name), by = "WELL.LABEL", all.x = TRUE))
  
  # Add "DMSO" to Compound if it's "LowControl" or "HighControl"
  df <- get(df_name)
  df$Compound[df$Compound %in% c("LowControl", "HighControl")] <- "DMSO"
  
  # Remove rows with NA values
  df <- na.omit(df)
  
  assign(df_name, df)
}
```

### 7\_Extract Controls.R

```
# Define a function to filter dataframe for DMSO
filter_dmsopm <- function(df) {
  df_dmsopm <- df[df$Compound == "DMSO", ]
  df_pm1 <- df[df$Compound != "DMSO", ]
  return(list(pm1 = df_pm1, pm1_dmsopm = df_dmsopm))
}

# Filter PL dataframes for DMSO
for (i in 585:614) {
  df_name <- paste0("PL", i)
  df_filtered <- filter_dmsopm(get(df_name))
  assign(df_name, df_filtered$pm1)
  assign(paste0(df_name, "_DMSO"), df_filtered$pm1_dmsopm)
}

# Filter PL dataframes from 615 to 644
for (i in 615:644) {
  df_name <- paste0("PL", i)
  df_filtered <- filter_dmsopm(get(df_name))
  assign(df_name, df_filtered$pm1)
  assign(paste0(df_name, "_DMSO"), df_filtered$pm1_dmsopm)
}

# Filter DMSO dataframes
for (i in 585:614) {
  df_name <- paste0("PL", i, "_DMSO")
  df <- get(df_name)
  df <- df[df$WELL.LABEL %in% c("M - 24", "N - 24", "O - 24", "P - 24"), ]
  assign(df_name, df)
}

for (i in 615:644) {
  df_name <- paste0("PL", i, "_DMSO")
  df <- get(df_name)
  df <- df[df$WELL.LABEL %in% c( "M - 24", "N - 24", "O - 24", "P - 24"), ]
  assign(df_name, df)
}
```

### 8\_Get Batches.R

```
# Create batches
Batch_1 <- do.call(rbind, mget(paste0("PL", 585:599)))
Batch_2 <- do.call(rbind, mget(paste0("PL", 601:614)))
Batch_3 <- do.call(rbind, mget(paste0("PL", 615:629)))
Batch_4 <- do.call(rbind, mget(paste0("PL", 630:644)))
Batch_5 <- PL600

# Combine DMSO batches
Batch_1_DMSO <- do.call(rbind, mget(paste0("PL", 585:599, "_DMSO")))
Batch_2_DMSO <- do.call(rbind, mget(paste0("PL", 601:614, "_DMSO")))
Batch_3_DMSO <- do.call(rbind, mget(paste0("PL", 615:629, "_DMSO")))
Batch_4_DMSO <- do.call(rbind, mget(paste0("PL", 630:644, "_DMSO")))
Batch_5_DMSO <- PL600_DMSO
```

### 9\_Batch Mean and SDs.R

```
Batch_1_Means <- colMeans(Batch_1_DMSO[,-c(1,2,40)])
Batch_1_Means <- c("Mean","Mean", Batch_1_Means,"Mean")
Batch_1_Means_DF <- rbind(Batch_1_DMSO,Batch_1_Means, stringsAsFactors = F)

Batch_1_SD <- Batch_1_DMSO %>%
  select_if(is.numeric) %>%
  summarise_all(sd, na.rm = TRUE) %>%
  unlist(use.names = FALSE) %>%
  c("SD","SD", .,"SD")
Batch_1_DMSO_Complete <- rbind(Batch_1_Means_DF,Batch_1_SD, stringsAsFactors = F)


# Calculate means for Batch 2 DMSO data
Batch_2_Means <- colMeans(Batch_2_DMSO[, -c(1, 2, 40)])
Batch_2_Means <- c("Mean", "Mean", Batch_2_Means, "Mean")
Batch_2_Means_DF <- rbind(Batch_2_DMSO, Batch_2_Means, stringsAsFactors = FALSE)

# Calculate standard deviation for Batch 2 DMSO data
Batch_2_SD <- Batch_2_DMSO %>%
  select_if(is.numeric) %>%
  summarise_all(sd, na.rm = TRUE) %>%
  unlist(use.names = FALSE) %>%
  c("SD", "SD", ., "SD")
Batch_2_DMSO_Complete <- rbind(Batch_2_Means_DF, Batch_2_SD, stringsAsFactors = FALSE)


# Calculate means for Batch 3 DMSO data
Batch_3_Means <- colMeans(Batch_3_DMSO[, -c(1, 2, 40)])
Batch_3_Means <- c("Mean", "Mean", Batch_3_Means, "Mean")
Batch_3_Means_DF <- rbind(Batch_3_DMSO, Batch_3_Means, stringsAsFactors = FALSE)

# Calculate standard deviation for Batch 3 DMSO data
Batch_3_SD <- Batch_3_DMSO %>%
  select_if(is.numeric) %>%
  summarise_all(sd, na.rm = TRUE) %>%
  unlist(use.names = FALSE) %>%
  c("SD", "SD", ., "SD")
Batch_3_DMSO_Complete <- rbind(Batch_3_Means_DF, Batch_3_SD, stringsAsFactors = FALSE)

# Calculate means for Batch 4 DMSO data
Batch_4_Means <- colMeans(Batch_4_DMSO[, -c(1, 2, 40)])
Batch_4_Means <- c("Mean", "Mean", Batch_4_Means, "Mean")
Batch_4_Means_DF <- rbind(Batch_4_DMSO, Batch_4_Means, stringsAsFactors = FALSE)

# Calculate standard deviation for Batch 4 DMSO data
Batch_4_SD <- Batch_4_DMSO %>%
  select_if(is.numeric) %>%
  summarise_all(sd, na.rm = TRUE) %>%
  unlist(use.names = FALSE) %>%
  c("SD", "SD", ., "SD")
Batch_4_DMSO_Complete <- rbind(Batch_4_Means_DF, Batch_4_SD, stringsAsFactors = FALSE)

# Calculate means for Batch 5 DMSO data
Batch_5_Means <- colMeans(Batch_5_DMSO[, -c(1, 2, 40)])
Batch_5_Means <- c("Mean", "Mean", Batch_5_Means, "Mean")
Batch_5_Means_DF <- rbind(Batch_5_DMSO, Batch_5_Means, stringsAsFactors = FALSE)

# Calculate standard deviation for Batch 5 DMSO data
Batch_5_SD <- Batch_5_DMSO %>%
  select_if(is.numeric) %>%
  summarise_all(sd, na.rm = TRUE) %>%
  unlist(use.names = FALSE) %>%
  c("SD", "SD", ., "SD")
Batch_5_DMSO_Complete <- rbind(Batch_5_Means_DF, Batch_5_SD, stringsAsFactors = FALSE)
```

### 10\_Add\_Means.R

```
Batch_1 <- rbind(Batch_1, Batch_1_Means,Batch_1_SD)
Batch_2 <- rbind(Batch_2, Batch_2_Means,Batch_2_SD)
Batch_3 <- rbind(Batch_3, Batch_3_Means,Batch_3_SD)
Batch_4 <- rbind(Batch_4, Batch_4_Means,Batch_4_SD)
Batch_5 <- rbind(Batch_5, Batch_5_Means,Batch_5_SD)
```

### 11\_Z\_Score\_Function.R

```
transform_dataframe <- function(input_df) {
  # Remove the first column and convert the remaining columns to numeric
  df_new <- input_df[, -c(1, 2, 40)]
  df_new <- df_new %>% mutate_all(as.numeric)
  
  # Determine the row indices for transformation
  last_row_index <- nrow(input_df)
  second_last_row_index <- nrow(input_df) - 1
  
  # Create a new dataframe for the transformed values
  df_normalized <- df_new  # Create a copy of the original dataframe
  
  # Divide each column by its corresponding value in the second last row
  for (i in 1:ncol(df_normalized)) {
    df_normalized[, i] <- df_normalized[, i] - df_new[second_last_row_index, i]
  }
  
  for (i in 1:ncol(df_normalized)) {
    df_normalized[, i] <- df_normalized[, i] / df_new[last_row_index, i]
  }
  
  # Add back the "Compound" column
  df_normalized <- cbind("Plate.ID" = deparse(substitute(input_df)), WELL.LABEL = input_df[, 1], Compound = input_df[, 40], df_normalized)
  
  return(df_normalized)
}
```

### 12\_Export Data.R

```
Batch_1_PhenoMeasures <- transform_dataframe(Batch_1)
Batch_2_PhenoMeasures <- transform_dataframe(Batch_2)
Batch_3_PhenoMeasures <- transform_dataframe(Batch_3)
Batch_4_PhenoMeasures <- transform_dataframe(Batch_4)
Batch_5_PhenoMeasures <- transform_dataframe(Batch_5)

Batch_1_Z_Score <- Batch_1_PhenoMeasures[-c(nrow(Batch_1_PhenoMeasures)-1, nrow(Batch_1_PhenoMeasures)), ]
Batch_2_Z_Score <- Batch_2_PhenoMeasures[-c(nrow(Batch_2_PhenoMeasures)-1, nrow(Batch_2_PhenoMeasures)), ]
Batch_3_Z_Score <- Batch_3_PhenoMeasures[-c(nrow(Batch_3_PhenoMeasures)-1, nrow(Batch_3_PhenoMeasures)), ]
Batch_4_Z_Score <- Batch_4_PhenoMeasures[-c(nrow(Batch_4_PhenoMeasures)-1, nrow(Batch_4_PhenoMeasures)), ]
Batch_5_Z_Score <- Batch_5_PhenoMeasures[-c(nrow(Batch_5_PhenoMeasures)-1, nrow(Batch_5_PhenoMeasures)), ]


All_Data <- rbind(Batch_1_Z_Score,Batch_2_Z_Score,Batch_3_Z_Score,Batch_4_Z_Score,Batch_5_Z_Score)


# Save All_Z_Scores to CSV
write.csv(All_Data, "DDU_Screen_1_Z_Score.csv", row.names = FALSE)
```

### 13\_Extract Hits.R

```
Hits <- All_Data %>% 
  mutate(Senescence_Status = ifelse(`Nuclei.Count.wv1` < -3 & Cells.Area.wv2 > 3, "Sen", "NonSen")) %>%
  filter(Senescence_Status == "Sen") 

duplicates <- All_Data %>% 
  mutate(Senescence_Status = ifelse(`Nuclei.Count.wv1` < -3 & Cells.Area.wv2 > 3, "Sen", "NonSen")) %>%
  filter(Senescence_Status == "Sen") %>%
  group_by(Compound) %>%  # Group by the columns you want to check for duplicates
  filter(n() > 1)  # Keep only rows that appear more than once  

write.csv(Hits$Compound, "Hit.csv" )
```
