## Supplementary material for "SAMP-Score: A morphology-based machine learning classification method for screening pro-senescence compounds in p16 positive cancer cells": HTML Walkthrough Guides for Code: Supplemental Data 1_6_Testing_Compound_Screen_1_Traditional.html

Building-Models\_Clusters\_Notebook.knit


### 1. Define Ground Truth

```
library("gplots")
```

```
## 
## Attaching package: 'gplots'
```

```
## The following object is masked from 'package:stats':
## 
##     lowess
```

```
library("RColorBrewer")
library("matrixStats")
library("plyr")
```

```
## 
## Attaching package: 'plyr'
```

```
## The following object is masked from 'package:matrixStats':
## 
##     count
```

```
library("dplyr")
```

```
## 
## Attaching package: 'dplyr'
```

```
## The following objects are masked from 'package:plyr':
## 
##     arrange, count, desc, failwith, id, mutate, rename, summarise,
##     summarize
```

```
## The following object is masked from 'package:matrixStats':
## 
##     count
```

```
## The following objects are masked from 'package:stats':
## 
##     filter, lag
```

```
## The following objects are masked from 'package:base':
## 
##     intersect, setdiff, setequal, union
```

```
library("data.table")
```

```
## 
## Attaching package: 'data.table'
```

```
## The following objects are masked from 'package:dplyr':
## 
##     between, first, last
```

```
library("stringr")
library("ggplot2")
library(glmnet)
```

```
## Loading required package: Matrix
```

```
## Loaded glmnet 4.1-8
```

```
library(caret)
```

```
## Loading required package: lattice
```

```
library(pROC)
```

```
## Type 'citation("pROC")' for a citation.
```

```
## 
## Attaching package: 'pROC'
```

```
## The following objects are masked from 'package:stats':
## 
##     cov, smooth, var
```

```
library(e1071)  # Contains functions for SVM
library(randomForest)
```

```
## randomForest 4.7-1.1
```

```
## Type rfNews() to see new features/changes/bug fixes.
```

```
## 
## Attaching package: 'randomForest'
```

```
## The following object is masked from 'package:ggplot2':
## 
##     margin
```

```
## The following object is masked from 'package:dplyr':
## 
##     combine
```

```
library(mda)
```

```
## Loading required package: class
```

```
## Loaded mda 0.5-4
```

```
library(neuralnet)
```

```
## 
## Attaching package: 'neuralnet'
```

```
## The following object is masked from 'package:dplyr':
## 
##     compute
```

```
# Upload your file 
all <- read.csv("All_Data_Clusters.csv")

#Remove Blanks
all_small <- all %>%
  filter(!grepl("Blank", Gene.Symbol))

### Sanity check - NA removed in previous stage 
all_removed <- all %>%
  filter(grepl("Blank", Gene.Symbol))

# Update DF ###
all <- all_small

#Deal with duplicate Gene Entries (12 in total)#
all <- all %>%
  group_by(RefSeq.Accession.Number) %>%
  mutate(RefSeq.Accession.Number = ifelse(row_number() == 1, RefSeq.Accession.Number, paste0(RefSeq.Accession.Number, "_A")))

#Add Ground Truth - Define filtering criteria
all <- all %>%
  mutate(Senescence_Status = ifelse(`Nuclei.Nuclei.Count.wv1` < -1.96 & Cells.Area.wv3 > 1.96, "Sen", "NonSen"))

all <- all[,-c(1,44)]
```

```
## 
## NonSen    Sen 
##    204    210
```

```
#Replace Test Data With New Unseen Data
combined_testData <- read.csv("DDU_Screen_1_Z_Score.csv")
combined_testData <- combined_testData %>%
  filter(across(where(is.numeric), ~ !is.na(.) & is.finite(.)))
```

```
## Warning: Using `across()` in `filter()` was deprecated in dplyr 1.0.8.
## ℹ Please use `if_any()` or `if_all()` instead.
## Call `lifecycle::last_lifecycle_warnings()` to see where this warning was
## generated.
```

```
Indexed_Compounds <- combined_testData[,c(1:3)]
combined_testData <- combined_testData [,-c(1:3)]

column_names<- colnames(trainData_part1)

colnames(combined_testData) <- column_names[-38]
```

### 3. Lasso Regularisation

```
# Load required libraries
library(glmnet)
library(caret)
library(pROC)
library(ggplot2)

# Ensure Senescence_Status is a factor (for classification)
trainData_part1$Senescence_Status <- as.factor(trainData_part1$Senescence_Status)
trainData_part2$Senescence_Status <- as.factor(trainData_part2$Senescence_Status)

# Prepare training and test data for Lasso
x_train <- as.matrix(trainData_part1[, -which(names(trainData_part1) == "Senescence_Status")])
y_train <- trainData_part1$Senescence_Status
x_test <- as.matrix(combined_testData)

# Fit the Lasso model on trainData_part1
set.seed(123)
lasso_model <- glmnet(x_train, y_train, family = "binomial", alpha = 1)

# Use cross-validation to determine the best lambda
cv_model <- cv.glmnet(x_train, y_train, family = "binomial", alpha = 1)
best_lambda <- cv_model$lambda.min  # or cv_model$lambda.1se for simplicity

# Refit the model using the best lambda value
final_model <- glmnet(x_train, y_train, family = "binomial", alpha = 1, lambda = best_lambda)

# Make predictions on trainData_part2 using the Lasso model
predicted_probabilities_part2 <- predict(final_model, newx = as.matrix(trainData_part2[, -which(names(trainData_part2) == "Senescence_Status")]), s = best_lambda, type = "response")
predicted_classes_part2 <- ifelse(predicted_probabilities_part2 > 0.5, "Sen", "NonSen")

# Create the meta-model training data frame
lasso_meta_model_data_training <- data.frame(
  Lasso_Prob = as.vector(predicted_probabilities_part2),
  Senescence_Status = trainData_part2$Senescence_Status,
  Predicted_Status = predicted_classes_part2
)

# Save the meta-model training data for further use
write.csv(lasso_meta_model_data_training, "lasso_meta_model_data_training.csv", row.names = FALSE)

## Now we are going to apply the model to testing data and evaluate

# Make predictions on testing data using the Lasso model
predicted_probabilities_test <- predict(final_model, newx = x_test, s = best_lambda, type = "response")
predicted_classes_test <- ifelse(predicted_probabilities_test > 0.5, "Sen", "NonSen")

# Create the meta-model testing data frame
lasso_meta_model_data_testing <- data.frame(
  Lasso_Prob = as.vector(predicted_probabilities_test),
  Predicted_Status = predicted_classes_test
)

# Save the meta-model testing data for further use
write.csv(lasso_meta_model_data_testing, "lasso_meta_model_data_testing.csv", row.names = FALSE)
```

### 10. Meta Model

```
set.seed(123)

# Load required libraries
library(glmnet)
library(caret)
library(pROC)
library(ggplot2)

# Combine model predictions and response variable for training
meta_model_data_train <- as.data.frame(cbind(
  lasso_meta_model_data_training[,1],
  svm_meta_model_data_training[,1],
  en_meta_model_data_training[,1],
  rf_meta_model_data_training[,1],
  mda_meta_model_training[,1],
  logistic_meta_model_data_training[,1],
  nn_meta_model_data_train[,1:2]
))

colnames(meta_model_data_train) <- c("lasso", "svm", "en", "rf", "mda", "logistic", "nn", "Senescence_Status")

# Extract the predictors and response for training
train_predictors <- meta_model_data_train[, -ncol(meta_model_data_train)]
train_response <- as.factor(meta_model_data_train$Senescence_Status)

# Fit the Lasso model on the training meta-model data
lasso_model <- cv.glmnet(
  x = as.matrix(train_predictors), 
  y = as.numeric(train_response) - 1,  # glmnet requires response to be zero-indexed
  family = "binomial", 
  alpha = 1
)

# Get the best lambda
best_lambda <- lasso_model$lambda.min

# Combine model predictions and response variable for testing
meta_model_data_test <- as.data.frame(cbind(
  lasso_meta_model_data_testing[,1],
  svm_meta_model_data_testing[,1],
  en_meta_model_data_testing[,1],
  rf_meta_model_data_testing[,1],
  mda_meta_model_testing[,1],
  logistic_meta_model_data_testing[,1],
  nn_meta_model_data_test[,1:2]
))

colnames(meta_model_data_test) <- c("lasso", "svm", "en", "rf", "mda", "logistic", "nn", "Senescence_Status")

# Extract the predictors and response for testing
test_predictors <- meta_model_data_test[, -ncol(meta_model_data_test)]
test_response <- as.factor(meta_model_data_test$Senescence_Status)

# Make predictions on the meta-model test data using the trained Lasso model
predicted_probabilities_test <- predict(lasso_model, newx = as.matrix(test_predictors), s = best_lambda, type = "response")
predicted_classes_test <- ifelse(predicted_probabilities_test > 0.5, "Sen", "NonSen")

predicted_classes_DDU_Screen <- cbind(Indexed_Compounds,predicted_classes_test,combined_testData)


write.csv(predicted_classes_DDU_Screen, "DDU_Screen_1_Predictions.csv")

# Logarithmic scatter plot
Log_Scatter_Plot <- ggplot(predicted_classes_DDU_Screen, aes(x = Nuclei.Nuclei.Count.wv1, y = Cells.Area.wv3, color = s1)) +
  geom_point(size = 0.1) +
  scale_y_log10() +
  labs(x = "Nuclei Count (Z Score)", y = "Cells Area (Log10 Z Score)", color = "Senescence Prediction") +
  theme_bw() +
  theme(text = element_text(size = 14))

# Save the plot
ggsave("Log_Scatter_Plot.png", plot = Log_Scatter_Plot, width = 8, height = 6)
```

```
## Warning in self$trans$transform(x): NaNs produced
```

```
## Warning: Transformation introduced infinite values in continuous y-axis
```

```
## Warning: Removed 4354 rows containing missing values (`geom_point()`).
```

```
# Non-logarithmic scatter plot
Scatter_Plot <- ggplot(predicted_classes_DDU_Screen, aes(x = Nuclei.Nuclei.Count.wv1, y = Cells.Area.wv3, color = s1)) +
  geom_point(size = 0.1) +
  labs(x = "Nuclei Count (Z Score)", y = "Cells Area (Z Score)", color = "Senescence Prediction") +
  theme_bw() +
  theme(text = element_text(size = 14))

# Save the plot
ggsave("Scatter_Plot.png", plot = Scatter_Plot, width = 8, height = 6)

# Count the total occurrences of each category in s1
total_counts <- predicted_classes_DDU_Screen %>%
  count(s1) %>%
  rename(Hit_Count = n)

# Count the occurrences of observations that satisfy the original conditions
conditional_counts_1_92 <- predicted_classes_DDU_Screen %>%
  filter(Nuclei.Nuclei.Count.wv1 <= -1.92, Cells.Area.wv3 >= 1.92) %>%
  count(s1) %>%
  rename(Threshold_Count_1_92 = n)

# Count the occurrences of observations that satisfy the stricter conditions (threshold of 3)
conditional_counts_3 <- predicted_classes_DDU_Screen %>%
  filter(Nuclei.Nuclei.Count.wv1 <= -3, Cells.Area.wv3 >= 3) %>%
  count(s1) %>%
  rename(Threshold_Count_3 = n)

# Merge the total counts with both sets of conditional counts
merged_counts <- total_counts %>%
  left_join(conditional_counts_1_92, by = "s1") %>%
  left_join(conditional_counts_3, by = "s1") %>%
  mutate(
    Threshold_Count_1_92 = ifelse(is.na(Threshold_Count_1_92), 0, Threshold_Count_1_92),
    Threshold_Count_3 = ifelse(is.na(Threshold_Count_3), 0, Threshold_Count_3)
  )

# Print the merged counts
print(merged_counts)
```

```
##       s1 Hit_Count Threshold_Count_1_92 Threshold_Count_3
## 1 NonSen     15319                   43                 0
## 2    Sen      2884                 2621              1195
```

```
write.csv(merged_counts, "Compare_Methods.csv")
```
