## Supplementary material for "SAMP-Score: A morphology-based machine learning classification method for screening pro-senescence compounds in p16 positive cancer cells": HTML Walkthrough Guides for Code: Supplemental Data 1_7_Testing_Compound_Screen_1_Clusters.html

### 10. Meta Model

```
set.seed(123)

# Combine model predictions and response variable for training
meta_model_data_train <- as.data.frame(cbind(
  lasso_meta_model_data_training[,1],
  svm_meta_model_data_training[,1],
  en_meta_model_data_training[,1],
  rf_meta_model_data_training[,1],
  mda_meta_model_training[,1],
  logistic_meta_model_data_training[,1],
  nn_meta_model_data_train[,1:2]
))

colnames(meta_model_data_train) <- c("lasso", "svm", "en", "rf", "mda", "logistic", "nn", "Senescence_Status")

# Extract the predictors and response for training
train_predictors <- meta_model_data_train[, -ncol(meta_model_data_train)]
train_response <- as.factor(meta_model_data_train$Senescence_Status)

# Fit the Lasso model on the training meta-model data
lasso_model <- cv.glmnet(
  x = as.matrix(train_predictors), 
  y = as.numeric(train_response) - 1,  # glmnet requires response to be zero-indexed
  family = "binomial", 
  alpha = 1
)

# Get the best lambda
best_lambda <- lasso_model$lambda.min

# Combine model predictions and response variable for testing
meta_model_data_test <- as.data.frame(cbind(
  lasso_meta_model_data_testing[,1],
  svm_meta_model_data_testing[,1],
  en_meta_model_data_testing[,1],
  rf_meta_model_data_testing[,1],
  mda_meta_model_testing[,1],
  logistic_meta_model_data_testing[,1],
  nn_meta_model_data_test[,1:2]
))

colnames(meta_model_data_test) <- c("lasso", "svm", "en", "rf", "mda", "logistic", "nn", "Senescence_Status")

# Extract the predictors and response for testing
test_predictors <- meta_model_data_test[, -ncol(meta_model_data_test)]
test_response <- as.factor(meta_model_data_test$Senescence_Status)

# Make predictions on the meta-model test data using the trained Lasso model
predicted_probabilities_test <- predict(lasso_model, newx = as.matrix(test_predictors), s = best_lambda, type = "response")
predicted_classes_test <- ifelse(predicted_probabilities_test > 0.9, "Sen", "NonSen")

# Combine predictions with the probability into the final data frame
predicted_classes_DDU_Screen <- as.data.frame(cbind(
  Indexed_Compounds,
  predicted_classes_test, 
  predicted_probabilities_test,
  combined_testData
))

# Rename columns for clarity
colnames(predicted_classes_DDU_Screen)[4:5] <- c("Prediction", "Probability")

# Save the results to a CSV file
write.csv(predicted_classes_DDU_Screen, "DDU_Screen_1_Predictions.csv")

# Logarithmic scatter plot
Log_Scatter_Plot <- ggplot(predicted_classes_DDU_Screen, aes(x = Nuclei.Nuclei.Count.wv1, y = Cells.Area.wv3, color = Prediction)) +
  geom_point(size = 0.1) +
  scale_y_log10() +
  labs(x = "Nuclei Count (Z Score)", y = "Cells Area (Log10 Z Score)", color = "Senescence Prediction") +
  theme_bw() +
  theme(text = element_text(size = 14))

# Save the plot
ggsave("Log_Scatter_Plot.png", plot = Log_Scatter_Plot, width = 8, height = 6)
```

```
## Warning in self$trans$transform(x): NaNs produced
```

```
## Warning: Transformation introduced infinite values in continuous y-axis
```

```
## Warning: Removed 4354 rows containing missing values (`geom_point()`).
```

```
# Non-logarithmic scatter plot
Scatter_Plot <- ggplot(predicted_classes_DDU_Screen, aes(x = Nuclei.Nuclei.Count.wv1, y = Cells.Area.wv3, color = Prediction)) +
  geom_point(size = 0.1) +
  labs(x = "Nuclei Count (Z Score)", y = "Cells Area (Z Score)", color = "Senescence Prediction") +
  theme_bw() +
  theme(text = element_text(size = 14))

# Save the plot
ggsave("Scatter_Plot.png", plot = Scatter_Plot, width = 8, height = 6)

# Count the total occurrences of each category in Prediction
total_counts <- predicted_classes_DDU_Screen %>%
  count(Prediction) %>%
  rename(Hit_Count = n)

# Count the occurrences of observations that satisfy the original conditions
conditional_counts_1_92 <- predicted_classes_DDU_Screen %>%
  filter(Nuclei.Nuclei.Count.wv1 <= -1.92, Cells.Area.wv3 >= 1.92) %>%
  count(Prediction) %>%
  rename(Threshold_Count_1_92 = n)

# Count the occurrences of observations that satisfy the stricter conditions (threshold of 3)
conditional_counts_3 <- predicted_classes_DDU_Screen %>%
  filter(Nuclei.Nuclei.Count.wv1 <= -3, Cells.Area.wv3 >= 3) %>%
  count(Prediction) %>%
  rename(Threshold_Count_3 = n)

# Merge the total counts with both sets of conditional counts
merged_counts <- total_counts %>%
  left_join(conditional_counts_1_92, by = "Prediction") %>%
  left_join(conditional_counts_3, by = "Prediction") %>%
  mutate(
    Threshold_Count_1_92 = ifelse(is.na(Threshold_Count_1_92), 0, Threshold_Count_1_92),
    Threshold_Count_3 = ifelse(is.na(Threshold_Count_3), 0, Threshold_Count_3)
  )

# Print the merged counts
print(merged_counts)
```

```
##   Prediction Hit_Count Threshold_Count_1_92 Threshold_Count_3
## 1     NonSen     14862                  694               470
## 2        Sen      3341                 1970               725
```

```
write.csv(merged_counts, "Compare_Methods.csv")


# Find unique Sen Compounds 

# Step 1: Identify compounds that appear as "Sen" only once in the entire dataset
single_sen_compounds <- predicted_classes_DDU_Screen %>%
  group_by(Compound) %>%
  filter(sum(Prediction == "Sen") == 1) %>%
  pull(Compound)  # Extract the list of such compounds

# Step 2: Filter the entire dataset to include only those compounds that are "Sen" only once
final_results <- predicted_classes_DDU_Screen %>%
  filter(Compound %in% single_sen_compounds, Prediction == "Sen")

write.csv(final_results, "Unique Compounds.csv")
```

12. Morphology Heatmaps

```
# Create a folder to store the heatmaps if it doesn't already exist
dir.create("Morphology_Heatmaps", showWarnings = FALSE)

# Combine data and add Dose column
Heatmap_Data <- cbind(Indexed_Compounds, predicted_classes_test, combined_testData)

# Add Dose column based on Plate.ID
Heatmap_Data <- Heatmap_Data %>%
  mutate(Dose = case_when(
    Plate.ID %in% c("Batch_1", "Batch_2", "Batch_5") ~ 10,
    Plate.ID %in% c("Batch_3", "Batch_4") ~ 50,
    TRUE ~ NA_real_ # In case there are Plate.IDs not listed, assign NA
  ))

# Remove Plate.ID column and replace with Dose in the data used for heatmaps
Heatmap <- Heatmap_Data[,-which(names(Heatmap_Data) == "Plate.ID")]

# List of unique Dose values and s1 categories
Doses <- unique(Heatmap$Dose)
s1_categories <- unique(Heatmap$s1)

# Loop through each Dose and s1 category to create and save a heatmap
for (dose in Doses) {
  for (category in s1_categories) {
    
    # Filter the data for the current Dose and s1 category
    Heatmap_adjusted <- Heatmap[Heatmap$Dose == dose & Heatmap$s1 == category, ]
    
    # Set row names to Compound and remove unnecessary columns
    rownames(Heatmap_adjusted) <- Heatmap_adjusted$Compound
    Heatmap_adjusted <- Heatmap_adjusted[,-c(1,2,3,41)]  # Remove Dose and s1 columns, keep Compound as row names
    
    # Convert to matrix
    all_matrix <- as.matrix(Heatmap_adjusted)  # Convert to matrix
    
    # Define color palette and breaks
    breaks <- unique(c(seq(-5, -1, length=100), seq(-1, 0.1, length=100), seq(1, 5, length=100)))
    my_palette <- colorRampPalette(c("yellow", "black", "black", "purple"))(length(breaks) - 1)
    
    # Save the heatmap as a PNG file
    png_filename <- paste0("Morphology_Heatmaps/Heatmap_Dose_", dose, "_", category, ".png")
    png(png_filename, width = 800, height = 800)
    
    # Create heatmap
    heatmap.2(t(all_matrix),
              Rowv = TRUE,
              Colv = TRUE,
              col = my_palette,
              breaks = breaks,
              density.info = "none",
              trace = "none",
              dendrogram = c("both"), 
              symm = FALSE, symkey = FALSE, symbreaks = TRUE,
              labRow = FALSE,
              labCol = FALSE,
              cexRow = 0.8,
              cexCol = 0.1,
              margins = c(8, 2),
              key.title = "1", 
              key.xlab = "Z Score", 
              main = paste("Heatmap for Dose", dose, " - ", category),  # Add a title
              sepcolor = c("black"), sepwidth = c(0.05, 0.05),
              distfun = function(x) dist(x, method = "euclidean"),
              hclust = function(x) hclust(x, method = "ward.D2"))
    
    dev.off()  # Close the PNG device
  }
}
```
