## Supplementary material for "SAMP-Score: A morphology-based machine learning classification method for screening pro-senescence compounds in p16 positive cancer cells": HTML Walkthrough Guides for Code: Supplemental Data 1_8_Wrangle_Compound_Screen_2.html

Wrangle-DDU-Screen-2.knit


### 1\_Import Data.R

```
# Load the libraries 
library("gplots")
```

```
## 
## Attaching package: 'gplots'
```

```
## The following object is masked from 'package:stats':
## 
##     lowess
```

```
library("RColorBrewer")
library("dplyr")
```

```
## 
## Attaching package: 'dplyr'
```

```
## The following objects are masked from 'package:stats':
## 
##     filter, lag
```

```
## The following objects are masked from 'package:base':
## 
##     intersect, setdiff, setequal, union
```

```
library("matrixStats")
```

```
## 
## Attaching package: 'matrixStats'
```

```
## The following object is masked from 'package:dplyr':
## 
##     count
```

```
B1_P1 <- read.csv("B1_P1.csv", fileEncoding = 'UTF-8-BOM')
B1_P2 <- read.csv("B1_P2.csv", fileEncoding = 'UTF-8-BOM')
B1_P3 <- read.csv("B1_P3.csv", fileEncoding = 'UTF-8-BOM')
B1_P4 <- read.csv("B1_P4.csv", fileEncoding = 'UTF-8-BOM')
B1_P5 <- read.csv("B1_P5.csv", fileEncoding = 'UTF-8-BOM')
B1_P6 <- read.csv("B1_P6.csv", fileEncoding = 'UTF-8-BOM')
B1_P7 <- read.csv("B1_P7.csv", fileEncoding = 'UTF-8-BOM')
B1_P8 <- read.csv("B1_P8.csv", fileEncoding = 'UTF-8-BOM')
B1_P9 <- read.csv("B1_P9.csv", fileEncoding = 'UTF-8-BOM')
B1_P10 <- read.csv("B1_P10.csv", fileEncoding = 'UTF-8-BOM')
B1_P11 <- read.csv("B1_P11.csv", fileEncoding = 'UTF-8-BOM')
B1_P12 <- read.csv("B1_P12.csv", fileEncoding = 'UTF-8-BOM')
B1_P13 <- read.csv("B1_P13.csv", fileEncoding = 'UTF-8-BOM')
B1_P14 <- read.csv("B1_P14.csv", fileEncoding = 'UTF-8-BOM')
B1_P15 <- read.csv("B1_P15.csv", fileEncoding = 'UTF-8-BOM')

B2_P1 <- read.csv("B2_P1.csv", fileEncoding = 'UTF-8-BOM')
B2_P2 <- read.csv("B2_P2.csv", fileEncoding = 'UTF-8-BOM')
B2_P3 <- read.csv("B2_P3.csv", fileEncoding = 'UTF-8-BOM')
B2_P4 <- read.csv("B2_P4.csv", fileEncoding = 'UTF-8-BOM')
B2_P5 <- read.csv("B2_P5.csv", fileEncoding = 'UTF-8-BOM')
B2_P6 <- read.csv("B2_P6.csv", fileEncoding = 'UTF-8-BOM')
B2_P7 <- read.csv("B2_P7.csv", fileEncoding = 'UTF-8-BOM')
B2_P8 <- read.csv("B2_P8.csv", fileEncoding = 'UTF-8-BOM')
B2_P9 <- read.csv("B2_P9.csv", fileEncoding = 'UTF-8-BOM')
B2_P10 <- read.csv("B2_P10.csv", fileEncoding = 'UTF-8-BOM')
B2_P11 <- read.csv("B2_P11.csv", fileEncoding = 'UTF-8-BOM')
B2_P12 <- read.csv("B2_P12.csv", fileEncoding = 'UTF-8-BOM')
B2_P13 <- read.csv("B2_P13.csv", fileEncoding = 'UTF-8-BOM')
B2_P14 <- read.csv("B2_P14.csv", fileEncoding = 'UTF-8-BOM')
B2_P15 <- read.csv("B2_P15.csv", fileEncoding = 'UTF-8-BOM')
```

### 2\_Tidy Measures.R

```
# Column Number for Measures
Measure_Names <-read.csv("Measure_Names.csv", fileEncoding = 'UTF-8-BOM')
```

```
## Warning in read.table(file = file, header = header, sep = sep, quote = quote, :
## incomplete final line found by readTableHeader on 'Measure_Names.csv'
```

```
Measure_Names <- colnames(Measure_Names)

B1_P1 <- B1_P1%>%
  select(one_of(Measure_Names))
# List of data frame names
data_frame_names <- c("B1_P1", "B1_P2", "B1_P3", "B1_P4", "B1_P5", "B1_P6", "B1_P7", "B1_P8", "B1_P9", "B1_P10",
                      "B1_P11", "B1_P12", "B1_P13", "B1_P14", "B1_P15", "B2_P1", "B2_P2", "B2_P3", "B2_P4", "B2_P5",
                      "B2_P6", "B2_P7", "B2_P8", "B2_P9", "B2_P10", "B2_P11", "B2_P12", "B2_P13", "B2_P14", "B2_P15")

# Create a list of data frames
data_frames_list <- lapply(data_frame_names, function(df_name) {
  read.csv(paste0(df_name, ".csv"), fileEncoding = 'UTF-8-BOM') %>%
    select(one_of(Measure_Names))
})

# Optionally, if you want to assign the modified data frames back to the original variables
# Note: This will overwrite the original data frames
for (i in seq_along(data_frame_names)) {
  assign(data_frame_names[i], data_frames_list[[i]])
}
```

### 3\_Extract siGlo.R

```
data_frame_names <- c("B1_P1", "B1_P2", "B1_P3", "B1_P4", "B1_P5", "B1_P6", "B1_P7", "B1_P8", "B1_P9", "B1_P10",
                      "B1_P11", "B1_P12", "B1_P13", "B1_P14", "B1_P15", "B2_P1", "B2_P2", "B2_P3", "B2_P4", "B2_P5",
                      "B2_P6", "B2_P7", "B2_P8", "B2_P9", "B2_P10", "B2_P11", "B2_P12", "B2_P13", "B2_P14", "B2_P15")


well_labels <- c("A - 23", "B - 23", "C - 23", "D - 23", "E - 23", "F - 23", "G - 23", "H - 23")

# Loop through each data frame and apply the filter
for (df_name in data_frame_names) {
  assign(paste0(df_name, "_siGlo"), get(df_name) %>%
           filter(WELL.LABEL %in% well_labels))
}
```

### 4\_Create Batches.R

```
# Update Batch_1
Batch_1_siGlo <- bind_rows(B1_P1_siGlo, B1_P2_siGlo, B1_P3_siGlo, B1_P4_siGlo, B1_P5_siGlo,
                           B1_P6_siGlo, B1_P7_siGlo, B1_P8_siGlo, B1_P9_siGlo, B1_P10_siGlo,
                           B1_P11_siGlo, B1_P12_siGlo, B1_P13_siGlo, B1_P14_siGlo, B1_P15_siGlo)

# Update Batch_2
Batch_2_siGlo <- bind_rows(B2_P1_siGlo, B2_P2_siGlo, B2_P3_siGlo, B2_P4_siGlo, B2_P5_siGlo,
                           B2_P6_siGlo, B2_P7_siGlo, B2_P8_siGlo, B2_P9_siGlo, B2_P10_siGlo,
                           B2_P11_siGlo, B2_P12_siGlo, B2_P13_siGlo, B2_P14_siGlo, B2_P15_siGlo)
```

### 5\_Calculate Batch Means.R

```
#Batch1
Batch_1_Means <- colMeans(Batch_1_siGlo[,-c(1,2)])
Batch_1_Means <- c("Mean","Mean", Batch_1_Means)
Batch_1_Means_DF <- rbind(Batch_1_siGlo,Batch_1_Means, stringsAsFactors = F)

Batch_1_SD <- Batch_1_siGlo %>%
  select_if(is.numeric) %>%
  summarise_all(sd, na.rm = TRUE) %>%
  unlist(use.names = FALSE) %>%
  c("SD","SD", .)
Batch_1_siGlo_Complete <- rbind(Batch_1_Means_DF,Batch_1_SD, stringsAsFactors = F)


#Batch2
Batch_2_Means <- colMeans(Batch_2_siGlo[,-c(1,2)])
Batch_2_Means <- c("Mean","Mean", Batch_2_Means)
Batch_2_Means_DF <- rbind(Batch_2_siGlo,Batch_2_Means, stringsAsFactors = F)

Batch_2_SD <- Batch_2_siGlo %>%
  select_if(is.numeric) %>%
  summarise_all(sd, na.rm = TRUE) %>%
  unlist(use.names = FALSE) %>%
  c("SD","SD", .)
Batch_2_siGlo_Complete <- rbind(Batch_2_Means_DF,Batch_2_SD, stringsAsFactors = F)
```

### 6\_Write Batch siGlo Controls.R

```
write.csv(Batch_1_siGlo_Complete,"Batch_1_Complete.csv", row.names = FALSE)

write.csv(Batch_2_siGlo_Complete,"Batch_2_Complete.csv", row.names = FALSE)
```

### 7\_Add Batch Means.R

```
#Add Batch 1 Mean/SD
B1_P1 <- rbind(B1_P1, Batch_1_Means,Batch_1_SD)
B1_P2 <- rbind(B1_P2, Batch_1_Means,Batch_1_SD)
B1_P3 <- rbind(B1_P3, Batch_1_Means,Batch_1_SD)
B1_P4 <- rbind(B1_P4, Batch_1_Means,Batch_1_SD)
B1_P5 <- rbind(B1_P5, Batch_1_Means,Batch_1_SD)
B1_P6 <- rbind(B1_P6, Batch_1_Means,Batch_1_SD)
B1_P7 <- rbind(B1_P7, Batch_1_Means,Batch_1_SD)
B1_P8 <- rbind(B1_P8, Batch_1_Means,Batch_1_SD)
B1_P9 <- rbind(B1_P9, Batch_1_Means,Batch_1_SD)
B1_P10 <- rbind(B1_P10, Batch_1_Means,Batch_1_SD)
B1_P11 <- rbind(B1_P11, Batch_1_Means,Batch_1_SD)
B1_P12 <- rbind(B1_P12, Batch_1_Means,Batch_1_SD)
B1_P13 <- rbind(B1_P13, Batch_1_Means,Batch_1_SD)
B1_P14 <- rbind(B1_P14, Batch_1_Means,Batch_1_SD)
B1_P15 <- rbind(B1_P15, Batch_1_Means,Batch_1_SD)


#Add Batch 2 Mean/SD
B2_P1 <- rbind(B2_P1, Batch_1_Means,Batch_1_SD)
B2_P2 <- rbind(B2_P2, Batch_1_Means,Batch_1_SD)
B2_P3 <- rbind(B2_P3, Batch_1_Means,Batch_1_SD)
B2_P4 <- rbind(B2_P4, Batch_1_Means,Batch_1_SD)
B2_P5 <- rbind(B2_P5, Batch_1_Means,Batch_1_SD)
B2_P6 <- rbind(B2_P6, Batch_1_Means,Batch_1_SD)
B2_P7 <- rbind(B2_P7, Batch_1_Means,Batch_1_SD)
B2_P8 <- rbind(B2_P8, Batch_1_Means,Batch_1_SD)
B2_P9 <- rbind(B2_P9, Batch_1_Means,Batch_1_SD)
B2_P10 <- rbind(B2_P10, Batch_1_Means,Batch_1_SD)
B2_P11 <- rbind(B2_P11, Batch_1_Means,Batch_1_SD)
B2_P12 <- rbind(B2_P12, Batch_1_Means,Batch_1_SD)
B2_P13 <- rbind(B2_P13, Batch_1_Means,Batch_1_SD)
B2_P14 <- rbind(B2_P14, Batch_1_Means,Batch_1_SD)
B2_P15 <- rbind(B2_P15, Batch_1_Means,Batch_1_SD)
```

### 8\_Create\_Z\_Function.R

```
transform_dataframe <- function(input_df) {
  # Remove the first column and convert the remaining columns to numeric
  B1_P1_New <- input_df[,-c(1,2)]
  B1_P1_New <- B1_P1_New %>% mutate_all(as.numeric)
  
  # Create a new dataframe for the transformed values
  B1_P1_Normalized <- B1_P1_New  # Create a copy of the original dataframe
  
  # Divide each column by its corresponding value in row 385
  for (i in 1:ncol(B1_P1_Normalized)) {
    B1_P1_Normalized[, i] <- B1_P1_Normalized[, i] - B1_P1_New[385, i]
  }
  
  for (i in 1:ncol(B1_P1_Normalized)) {
    B1_P1_Normalized[, i] <- B1_P1_Normalized[, i] / B1_P1_New[386, i]
  }
  
  # Add the first column back to the transformed dataframe with the name "WELL.LABEL"
  B1_P1_Normalized <- cbind("Plate Name" = deparse(substitute(input_df)), WELL.LABEL = input_df[, 2], B1_P1_Normalized)
  
  return(B1_P1_Normalized)
}

# Example usage:
# Assuming you have a dataframe named B1_P1
# B1_P1_PhenoMeasures <- transform_dataframe(B1_P1)
```

### 9\_Z\_Score\_Transformation.R

```
B1_P1_PhenoMeasures <- transform_dataframe(B1_P1)
B1_P2_PhenoMeasures <- transform_dataframe(B1_P2)
B1_P3_PhenoMeasures <- transform_dataframe(B1_P3)
B1_P4_PhenoMeasures <- transform_dataframe(B1_P4)
B1_P5_PhenoMeasures <- transform_dataframe(B1_P5)
B1_P6_PhenoMeasures <- transform_dataframe(B1_P6)
B1_P7_PhenoMeasures <- transform_dataframe(B1_P7)
B1_P8_PhenoMeasures <- transform_dataframe(B1_P8)
B1_P9_PhenoMeasures <- transform_dataframe(B1_P9)
B1_P10_PhenoMeasures <- transform_dataframe(B1_P10)
B1_P11_PhenoMeasures <- transform_dataframe(B1_P11)
B1_P12_PhenoMeasures <- transform_dataframe(B1_P12)
B1_P13_PhenoMeasures <- transform_dataframe(B1_P13)
B1_P14_PhenoMeasures <- transform_dataframe(B1_P14)
B1_P15_PhenoMeasures <- transform_dataframe(B1_P15)

# Repeat the process for B2_P1 to B2_P15
B2_P1_PhenoMeasures <- transform_dataframe(B2_P1)
B2_P2_PhenoMeasures <- transform_dataframe(B2_P2)
B2_P3_PhenoMeasures <- transform_dataframe(B2_P3)
B2_P4_PhenoMeasures <- transform_dataframe(B2_P4)
B2_P5_PhenoMeasures <- transform_dataframe(B2_P5)
B2_P6_PhenoMeasures <- transform_dataframe(B2_P6)
B2_P7_PhenoMeasures <- transform_dataframe(B2_P7)
B2_P8_PhenoMeasures <- transform_dataframe(B2_P8)
B2_P9_PhenoMeasures <- transform_dataframe(B2_P9)
B2_P10_PhenoMeasures <- transform_dataframe(B2_P10)
B2_P11_PhenoMeasures <- transform_dataframe(B2_P11)
B2_P12_PhenoMeasures <- transform_dataframe(B2_P12)
B2_P13_PhenoMeasures <- transform_dataframe(B2_P13)
B2_P14_PhenoMeasures <- transform_dataframe(B2_P14)
B2_P15_PhenoMeasures <- transform_dataframe(B2_P15)
```

### 10\_Export\_All\_Data.R

```
# List of PhenoMeasures dataframe names
df_names <- c("B1_P1_PhenoMeasures", "B1_P2_PhenoMeasures", "B1_P3_PhenoMeasures", "B1_P4_PhenoMeasures", "B1_P5_PhenoMeasures",
              "B1_P6_PhenoMeasures", "B1_P7_PhenoMeasures", "B1_P8_PhenoMeasures", "B1_P9_PhenoMeasures", "B1_P10_PhenoMeasures",
              "B1_P11_PhenoMeasures", "B1_P12_PhenoMeasures", "B1_P13_PhenoMeasures", "B1_P14_PhenoMeasures", "B1_P15_PhenoMeasures",
              "B2_P1_PhenoMeasures", "B2_P2_PhenoMeasures", "B2_P3_PhenoMeasures", "B2_P4_PhenoMeasures", "B2_P5_PhenoMeasures",
              "B2_P6_PhenoMeasures", "B2_P7_PhenoMeasures", "B2_P8_PhenoMeasures", "B2_P9_PhenoMeasures", "B2_P10_PhenoMeasures",
              "B2_P11_PhenoMeasures", "B2_P12_PhenoMeasures", "B2_P13_PhenoMeasures", "B2_P14_PhenoMeasures", "B2_P15_PhenoMeasures")

# Combine the dataframes into one large dataframe
All_Z_Scores <- do.call(rbind, lapply(df_names, get))

library(stringr)
filtered_data <- All_Z_Scores  %>%
  filter(!str_detect(WELL.LABEL, "- 11|- 12|- 23|- 24"))

filtered_data <- filtered_data[-c(9660,9659),]

filtered_data <- filtered_data  %>%
  filter(!str_detect(WELL.LABEL, "Mean|SD"))

Compounds <- read.csv("DDU_Locations.csv", fileEncoding = 'UTF-8-BOM')
Compounds<- rbind(Compounds,Compounds)

All_Compound_Scores <- cbind(Compounds , filtered_data)

All_Compound_Scores <-All_Compound_Scores %>%
  filter(!PlateBarcode == "X")

All_Compound_Scores <- All_Compound_Scores[,-c(1,3,5:8)]

All_Compound_Scores <- All_Compound_Scores  %>%
  group_by(DDDNumber, Concentration) %>%
  summarize_all(mean, na.rm = TRUE)

# View the resulting dataframe

# Save All_Z_Scores to CSV
write.csv(All_Compound_Scores , "All_Compound_Scores.csv", row.names = FALSE)
```
