## Supplementary material for "SAMP-Score: A morphology-based machine learning classification method for screening pro-senescence compounds in p16 positive cancer cells": HTML Walkthrough Guides for Code: Supplemental Data 1_9_Testing_Compound_Screen_2_Clusters.html

```
print(table(trainData_part2$Senescence_Status))
```

```
## 
## NonSen    Sen 
##   1976   1996
```

```
#Replace Test Data With New Unseen Data
combined_testData <- read.csv("All_Compound_Scores.csv")
combined_testData <- combined_testData %>%
  filter(across(where(is.numeric), ~ !is.na(.) & is.finite(.)))
```

```
## Warning: Using `across()` in `filter()` was deprecated in dplyr 1.0.8.
## ℹ Please use `if_any()` or `if_all()` instead.
## Call `lifecycle::last_lifecycle_warnings()` to see where this warning was
## generated.
```

```
Indexed_Compounds <- combined_testData[,c(1:2)]
combined_testData <- combined_testData [,-c(1:2)]

column_names<- colnames(trainData_part1)

colnames(combined_testData) <- column_names[-38]
```

### 10. Meta Model

```
set.seed(123)

# Load required libraries
library(glmnet)
library(caret)
library(pROC)
library(ggplot2)

# Combine model predictions and response variable for training
meta_model_data_train <- as.data.frame(cbind(
  lasso_meta_model_data_training[,1],
  svm_meta_model_data_training[,1],
  en_meta_model_data_training[,1],
  rf_meta_model_data_training[,1],
  mda_meta_model_training[,1],
  logistic_meta_model_data_training[,1],
  nn_meta_model_data_train[,1:2]
))

colnames(meta_model_data_train) <- c("lasso", "svm", "en", "rf", "mda", "logistic", "nn", "Senescence_Status")

# Extract the predictors and response for training
train_predictors <- meta_model_data_train[, -ncol(meta_model_data_train)]
train_response <- as.factor(meta_model_data_train$Senescence_Status)

# Fit the Lasso model on the training meta-model data
lasso_model <- cv.glmnet(
  x = as.matrix(train_predictors), 
  y = as.numeric(train_response) - 1,  # glmnet requires response to be zero-indexed
  family = "binomial", 
  alpha = 1
)

# Get the best lambda
best_lambda <- lasso_model$lambda.min

# Combine model predictions and response variable for testing
meta_model_data_test <- as.data.frame(cbind(
  lasso_meta_model_data_testing[,1],
  svm_meta_model_data_testing[,1],
  en_meta_model_data_testing[,1],
  rf_meta_model_data_testing[,1],
  mda_meta_model_testing[,1],
  logistic_meta_model_data_testing[,1],
  nn_meta_model_data_test[,1:2]
))

colnames(meta_model_data_test) <- c("lasso", "svm", "en", "rf", "mda", "logistic", "nn", "Senescence_Status")

# Extract the predictors and response for testing
test_predictors <- meta_model_data_test[, -ncol(meta_model_data_test)]
test_response <- as.factor(meta_model_data_test$Senescence_Status)

# Make predictions on the meta-model test data using the trained Lasso model
predicted_probabilities_test <- predict(lasso_model, newx = as.matrix(test_predictors), s = best_lambda, type = "response")
predicted_classes_test <- ifelse(predicted_probabilities_test > 0.9, "Sen", "NonSen")

predicted_classes_DDU_Screen <- cbind(Indexed_Compounds,predicted_classes_test,combined_testData)


write.csv(predicted_classes_DDU_Screen, "DDU_Screen_1_Predictions.csv")

# Logarithmic scatter plot
Log_Scatter_Plot <- ggplot(predicted_classes_DDU_Screen, aes(x = Nuclei.Nuclei.Count.wv1, y = Cells.Area.wv3, color = s1)) +
  geom_point(size = 0.1) +
  scale_y_log10() +
  labs(x = "Nuclei Count (Z Score)", y = "Cells Area (Log10 Z Score)", color = "Senescence Prediction") +
  geom_vline(xintercept = -1.92, linetype = "dotted", color = "blue", size = 1) +  # Vertical line at x = 1.92
  geom_hline(yintercept = 1.92, linetype = "dotted", color = "black", size = 1) +   # Horizontal line at y = 1.92
  theme_bw() +
  theme(text = element_text(size = 14))
```

```
## Warning: Using `size` aesthetic for lines was deprecated in ggplot2 3.4.0.
## ℹ Please use `linewidth` instead.
## This warning is displayed once every 8 hours.
## Call `lifecycle::last_lifecycle_warnings()` to see where this warning was
## generated.
```

```
# Save the plot
ggsave("Log_Scatter_Plot.png", plot = Log_Scatter_Plot, width = 8, height = 6)
```

```
## Warning in self$trans$transform(x): NaNs produced
```

```
## Warning: Transformation introduced infinite values in continuous y-axis
```

```
## Warning: Removed 1490 rows containing missing values (`geom_point()`).
```

```
# Non-logarithmic scatter plot
Scatter_Plot <- ggplot(predicted_classes_DDU_Screen, aes(x = Nuclei.Nuclei.Count.wv1, y = Cells.Area.wv3, color = s1)) +
  geom_point(size = 0.1) +
  labs(x = "Nuclei Count (Z Score)", y = "Cells Area (Z Score)", color = "Senescence Prediction") +
  theme_bw() +
  theme(text = element_text(size = 14))

# Save the plot
ggsave("Scatter_Plot.png", plot = Scatter_Plot, width = 8, height = 6)
```

### 11. Plot SAMP Scores

```
# Create a folder to store the outputs if it doesn't already exist
output_dir <- "SAMP_Score_Plots"
dir.create(output_dir, showWarnings = FALSE)

library(tidyr)
```

```
## 
## Attaching package: 'tidyr'
```

```
## The following objects are masked from 'package:Matrix':
## 
##     expand, pack, unpack
```

```
library(gplots)
library(ggplot2)

Heatmap_Data <- cbind(Indexed_Compounds,predicted_probabilities_test, combined_testData)

Heatmap <- Heatmap_Data[,c(1,2,3)]

Heatmap_adjusted <- spread(Heatmap, key = Concentration, value = s1)
Heatmap_adjusted <- Heatmap_adjusted %>%
  mutate(across(everything(), ~ifelse(is.na(.), 0, .)))

rownames(Heatmap_adjusted) <- Heatmap_adjusted[,1]
Heatmap_adjusted <- Heatmap_adjusted[,-1]

all_matrix <- as.matrix(Heatmap_adjusted)

# Create heatmap
breaks <- unique(c(seq(0,0.87,length=100),seq(0.87,0.89,length=100), seq(0.9,1,length=100)))
my_palette <- colorRampPalette(c("black","black","#D3EC01","#46D100"))(length(breaks)-1)

# Save heatmap to file
heatmap_filename <- file.path(output_dir, "Heatmap.png")
png(heatmap_filename, width = 800, height = 800)

heatmap.2(all_matrix,
          Rowv = TRUE,
          Colv = FALSE,
          col = my_palette,
          breaks = breaks,
          density.info = "none",
          trace = "none",
          dendrogram = c("row"), 
          symm = FALSE, symkey = FALSE, symbreaks = TRUE,
          labRow = FALSE,
          labCol = colnames(all_matrix),
          cexRow = 0.1,
          cexCol = 2,
          margins = c(8,2),
          key.title = "1", 
          key.xlab = "Z Score", 
          sepcolor = c("black"), 
          sepwidth = c(0.05, 0.05),
          distfun = function(x) dist(x, method = "euclidean"),
          hclust = function(x) hclust(x, method = "ward.D2"))

dev.off() # Close the PNG device
```

```
## quartz_off_screen 
##                 2
```

```
# Create scatter plot and save it to file
plot <- ggplot(Heatmap, aes(x = Concentration, y = s1)) +
  geom_point() +
  labs(x = "Variables", y = "SAMP-Score", title = "Scatter Plot of All Doses / Compounds") +
  theme(axis.text.x = element_text(angle = 45, hjust = 1))

plot_filename <- file.path(output_dir, "Scatter_Plot.png")
ggsave(plot_filename, plot = plot, width = 8, height = 6)
```

#12. Morphology Heatmaps

```
Heatmap_Data <- cbind(Indexed_Compounds,predicted_probabilities_test, combined_testData)

library(tidyr)
library(gplots) # For heatmap.2 function

# Create a folder to store the heatmaps if it doesn't already exist
dir.create("Morphology_Heatmaps", showWarnings = FALSE)

# Remove the s1 column (column 3) but keep columns 1 and 2
Heatmap <- Heatmap_Data[,-3]

# List of unique concentrations
concentrations <- unique(Heatmap$Concentration)

# Loop through each concentration to create and save a heatmap
for (conc in concentrations) {
  
  # Filter the data for the current concentration
  Heatmap_adjusted <- Heatmap[Heatmap$Concentration == conc, ]
  
  # Set row names to DDDNumber and remove the Concentration column
  rownames(Heatmap_adjusted) <- Heatmap_adjusted$DDDNumber
  Heatmap_adjusted <- Heatmap_adjusted[,-1]  # Remove Concentration column, keep DDDNumber as row names
  
  # Convert to matrix
  all_matrix <- as.matrix(Heatmap_adjusted[,-1])  # Exclude DDDNumber from the matrix
  
  # Define color palette and breaks
  breaks <- unique(c(seq(-5,-1,length=100),seq(-1,0.1,length=100), seq(1,5,length=100)))
  my_palette <- colorRampPalette(c("yellow","black","black","purple"))(length(breaks)-1)
  
  # Save the heatmap as a PNG file
  png_filename <- paste0("Morphology_Heatmaps/Heatmap_Concentration_", conc, ".png")
  png(png_filename, width = 800, height = 800)
  
  # Create heat map
  heatmap.2(t(all_matrix),
            Rowv = T,
            Colv = T,
            col=my_palette,
            breaks=breaks,
            density.info="none",
            trace="none",
            #main = "199 compounds; Length or Prev",
            dendrogram=c("both"), 
            symm=F,symkey=F,symbreaks=T,
            labRow=F,
            labCol=F,
            cexRow=0.8,
            cexCol=0.1,
            margins = c(8,2),
            key.title = "1" , key.xlab="Z Score", 
            #rowsep =c(0,4,12,19,31,37,43),
            #colsep =c(0,233),
            sepcolor = c("black"),sepwidth = c(0.05, 0.05),
            #ColSideColors=condition_colors, scale="none",
            distfun = function(x) dist(x, method = "euclidean"),
            hclust=function(x) hclust(x,method="ward.D2"))
  
  dev.off()  # Close the PNG device
}
```

#13 Morphology of QM0005928

```
Heatmap_Data <- cbind(Indexed_Compounds, predicted_probabilities_test, combined_testData)

library(tidyr)
library(gplots) # For heatmap.2 function

# Create a folder to store the heatmaps if it doesn't already exist
dir.create("Morphology_Heatmaps", showWarnings = FALSE)

# Remove the s1 column (column 3) but keep columns 1 and 2
Heatmap <- Heatmap_Data[,-3]

# Specify the compound you're interested in
compound_of_interest <- "QM0005928"  # Replace with the actual compound name

# Filter the data for the specified compound
Heatmap_adjusted <- Heatmap[Heatmap$DDDNumber == compound_of_interest, ]

# Check if the filtering resulted in data
if (nrow(Heatmap_adjusted) == 0) {
  stop("No data found for the specified compound.")
}

# Update the Concentration values and ensure it is numeric
Heatmap_adjusted$Concentration <- as.numeric(Heatmap_adjusted$Concentration) * 1000000

# Sort by Concentration
Heatmap_adjusted <- Heatmap_adjusted[order(Heatmap_adjusted$Concentration), ]

# Set row names to Concentration and remove the Concentration column
rownames(Heatmap_adjusted) <- Heatmap_adjusted$Concentration
Heatmap_adjusted <- Heatmap_adjusted[,-1]  # Remove Concentration column, keep it as row names

# Convert to matrix
all_matrix <- as.matrix(Heatmap_adjusted)

# Define color palette and breaks
breaks <- unique(c(seq(-5, -1, length=100), seq(-1, 0.1, length=100), seq(1, 5, length=100)))
my_palette <- colorRampPalette(c("yellow", "black", "black", "purple"))(length(breaks) - 1)

# Save the heatmap as a PNG file
png_filename <- paste0("Morphology_Heatmaps/Heatmap_Compound_", compound_of_interest, ".png")
png(png_filename, width = 800, height = 800)

# Create heat map
heatmap.2(t(all_matrix),
          Rowv = TRUE,
          Colv = NA,  # Turn off column clustering
          col = my_palette,
          breaks = breaks,
          density.info = "none",
          trace = "none",
          dendrogram = c("row"),  # Only cluster rows
          symm = FALSE, symkey = FALSE, symbreaks = TRUE,
          labRow = FALSE,
          labCol = rownames(all_matrix),
          cexRow = 0.8,
          cexCol = 2,
          margins = c(8, 2),
          key.title = "1", key.xlab = "Z Score",
          sepcolor = c("black"), sepwidth = c(0.05, 0.05),
          distfun = function(x) dist(x, method = "euclidean"),
          hclust = function(x) hclust(x, method = "ward.D2"))

dev.off()  # Close the PNG device
```

```
## quartz_off_screen 
##                 2
```

### Line Chart QM0005928

```
# Load necessary libraries
library(ggplot2)
library(dplyr)

# Create a folder to store the line graphs if it doesn't already exist
dir.create("Morphology_LineGraphs", showWarnings = FALSE)

# Remove the s1 column (column 3) but keep columns 1 and 2
Line_Chart_Data <- Heatmap_Data

# Specify the compound you're interested in
compound_of_interest <- "QM0005928"  # Replace with the actual compound name

# Filter the data for the specified compound
Line_Chart_Data_Adjusted<- Line_Chart_Data[Line_Chart_Data$DDDNumber == compound_of_interest, ]

# Check if the filtering resulted in data
if (nrow(Line_Chart_Data_Adjusted) == 0) {
  stop("No data found for the specified compound.")
}

# Update the Concentration values and ensure it is numeric
Line_Chart_Data_Adjusted$Concentration <- as.numeric(Line_Chart_Data_Adjusted$Concentration) * 1000000

# Sort by Concentration
Line_Chart_Data_Adjusted<- Line_Chart_Data_Adjusted[order(Line_Chart_Data_Adjusted$Concentration), ]

# Extract dose and s1 columns for plotting
dose_vs_s1 <- Line_Chart_Data_Adjusted %>%
  select(Concentration, s1)  # Ensure s1 is included

# Create a line plot of Dose vs s1
png_filename <- paste0("Morphology_LineGraphs/LineGraph_Compound_", compound_of_interest, ".png")
png(png_filename, width = 800, height = 600)

ggplot(dose_vs_s1, aes(x = log10(Concentration), y = s1)) +
  geom_line() +
  geom_point() +
  labs(title = paste("Dose vs SAMP-Score for Compound", compound_of_interest),
       x = "Log10 Dose (Scaled Concentration)",
       y = "SAMP-Score") +
  theme_bw() +  # Use black and white theme
  theme(
    text = element_text(size = 16),  # Increase text size for titles and labels
    axis.title = element_text(size = 14),  # Increase text size for axis titles
    axis.text = element_text(size = 12)   # Increase text size for axis labels
  )

dev.off()  # Close the PNG device
```

```
## quartz_off_screen 
##                 2
```
