## Supplementary material for "SAMP-Score: A morphology-based machine learning classification method for screening pro-senescence compounds in p16 positive cancer cells": HTML Walkthrough Guides for Code: Supplemental Data 1_10_Testing_Other_Cancer_Lines.html

```
print(table(trainData_part2$Senescence_Status))
```

```
## 
## NonSen    Sen 
##   1976   1996
```

```
#Replace Test Data With New Unseen Data
combined_testData <- read.csv("Cell_Line_Z_Scores.csv")
```

```
## Warning in read.table(file = file, header = header, sep = sep, quote = quote, :
## incomplete final line found by readTableHeader on 'Cell_Line_Z_Scores.csv'
```

### 10. Meta Model

```
set.seed(123)

# Load required libraries
library(glmnet)
library(caret)
library(pROC)
library(ggplot2)

# Combine model predictions and response variable for training
meta_model_data_train <- as.data.frame(cbind(
  lasso_meta_model_data_training[,1],
  svm_meta_model_data_training[,1],
  en_meta_model_data_training[,1],
  rf_meta_model_data_training[,1],
  mda_meta_model_training[,1],
  logistic_meta_model_data_training[,1],
  nn_meta_model_data_train[,1:2]
))

colnames(meta_model_data_train) <- c("lasso", "svm", "en", "rf", "mda", "logistic", "nn", "Senescence_Status")

# Extract the predictors and response for training
train_predictors <- meta_model_data_train[, -ncol(meta_model_data_train)]
train_response <- as.factor(meta_model_data_train$Senescence_Status)

# Fit the Lasso model on the training meta-model data
lasso_model <- cv.glmnet(
  x = as.matrix(train_predictors), 
  y = as.numeric(train_response) - 1,  # glmnet requires response to be zero-indexed
  family = "binomial", 
  alpha = 1
)

# Get the best lambda
best_lambda <- lasso_model$lambda.min

# Combine model predictions and response variable for testing
meta_model_data_test <- as.data.frame(cbind(
  lasso_meta_model_data_testing[,1],
  svm_meta_model_data_testing[,1],
  en_meta_model_data_testing[,1],
  rf_meta_model_data_testing[,1],
  mda_meta_model_testing[,1],
  logistic_meta_model_data_testing[,1],
  nn_meta_model_data_test[,1:2]
))

colnames(meta_model_data_test) <- c("lasso", "svm", "en", "rf", "mda", "logistic", "nn", "Senescence_Status")

# Extract the predictors and response for testing
test_predictors <- meta_model_data_test[, -ncol(meta_model_data_test)]
test_response <- as.factor(meta_model_data_test$Senescence_Status)

# Make predictions on the meta-model test data using the trained Lasso model
predicted_probabilities_test <- predict(lasso_model, newx = as.matrix(test_predictors), s = best_lambda, type = "response")
predicted_classes_test <- ifelse(predicted_probabilities_test > 0.9, "Sen", "NonSen")

predicted_classes_DDU_Screen <- cbind(Indexed_Compounds,predicted_classes_test,combined_testData)


write.csv(predicted_classes_DDU_Screen, "Predictions.csv")
```

### 11. Plot SAMP Scores

```
# Create a folder to store the outputs if it doesn't already exist
output_dir <- "SAMP_Score_Plots"
dir.create(output_dir, showWarnings = FALSE)

library(tidyr)
```

```
## 
## Attaching package: 'tidyr'
```

```
## The following objects are masked from 'package:Matrix':
## 
##     expand, pack, unpack
```

```
library(gplots)
library(ggplot2)

Heatmap_Data <- cbind(Indexed_Compounds,predicted_probabilities_test, combined_testData)

Heatmap <- Heatmap_Data[,-c(2,3)]

rownames(Heatmap) <- Heatmap[,1]
Heatmap <- Heatmap[,-1]

all_matrix <- as.matrix(Heatmap)

# Create heatmap
breaks <- unique(c(seq(-5, -1, length=100), seq(-1, 0.1, length=100), seq(1, 5, length=100)))
my_palette <- colorRampPalette(c("yellow", "black", "black", "purple"))(length(breaks) - 1)

# Save heatmap to file
heatmap_filename <- file.path(output_dir, "Heatmap.png")
png(heatmap_filename, width = 800, height = 800)

heatmap.2(t(all_matrix),
          Rowv = TRUE,
          Colv = T,
          col = my_palette,
          breaks = breaks,
          density.info = "none",
          trace = "none",
          dendrogram = c("both"), 
          symm = FALSE, symkey = FALSE, symbreaks = TRUE,
          labRow = FALSE,
          labCol = rownames(all_matrix),
          cexRow = 0.1,
          cexCol = 1,
          margins = c(8,5),
          key.title = "1", 
          key.xlab = "Z Score", 
          sepcolor = c("black"), 
          sepwidth = c(0.05, 0.05),
          distfun = function(x) dist(x, method = "euclidean"),
          hclust = function(x) hclust(x, method = "ward.D2"))

dev.off() # Close the PNG device
```

```
## quartz_off_screen 
##                 2
```

```
write.csv(Heatmap_Data[,1:4], "Pred_Coeffs.csv")
```
